# Mapping active cis-regulatory elements from transcription initiation events

**DOI:** 10.64898/2026.05.11.724207

**Authors:** Hjörleifur Einarsson, Natsuda Navamajiti, Christian Skov Vaagensø, Stefano Pellegrini, Lykke Jirström, Nicolas Alcaraz, Malte Thodberg, Daniel A. Solvie, Wei-Lin Qiu, Maya U. Sheth, Marta Greedy Escudero, Bram L. Gorissen, Marco Salvatore, Takeya Kasukawa, Hazuki Takahashi, Piero Carninci, Bradley E. Bernstein, Albin Sandelin, Jesse M. Engreitz, Robert Krautz, Robin Andersson

**Affiliations:** Section for Computational and RNA Biology, Department of Biology, University of Copenhagen, Denmark; Novo Nordisk Foundation Center for Genomic Mechanisms of Disease, Broad Institute of MIT and Harvard, Cambridge, MA, USA; Current address: Institute for Research in Biomedicine, The Barcelona Institute of Science and Technology, Spain; Novo Nordisk Foundation Center for Protein Research, Department of Cellular and Molecular Medicine, University of Copenhagen, Denmark; Center for Epigenetic Cell Memory (EpiC), Danish Cancer Institute, Danish Cancer Society, Denmark; Novo Nordisk Foundation Center for Basic Metabolic Research, University of Copenhagen, Denmark; Department of Cancer Biology, Dana Farber Cancer Institute, Boston, MA, USA; Departments of Cell Biology and Pathology, Harvard Medical School, Boston, MA, USA; Department of Bioengineering, Stanford University, Stanford, CA, USA; Department of Genetics, Stanford University School of Medicine, Stanford, CA, USA; Basic Sciences and Engineering Initiative, Betty Irene Moore Children’s Heart Center, Lucile Packard Children’s Hospital, Stanford, CA, USA; Broad Institute of MIT and Harvard, Cambridge, MA, USA; ScienTek ApS, Silkeborg, Denmark & Hikari Solutions SRL, Italy; RIKEN Center for Integrative Medical Sciences, Yokohama, Japan; Human Technopole, Milan, Italy; Ludwig Center at Harvard Medical School, Boston, MA, USA; Gene Regulation Observatory, Broad Institute of MIT and Harvard, Cambridge, MA, USA; Biotech Research & Innovation Centre, University of Copenhagen, Denmark; Stanford Cardiovascular Institute, Stanford University School of Medicine, Stanford, CA, USA

**Author notes:** &. Equal contribution as first authors (Hjörleifur Einarsson, Natsuda Navamajiti). Authors have agreed that either can be listed first. Equal contribution as last authors (Robert Krautz, Robin Andersson). Authors have agreed that either can be listed last.

## Abstract

Determining the activity of cis-regulatory elements (CREs) is essential for modeling gene regulation and interpreting genetic variation. Yet, current methods often lack the specificity to distinguish active regulation from permissive chromatin, the sensitivity to detect unstable enhancer RNAs, or the scalability required to profile limited input material and primary cells. Here, we introduce nucCAGE, a transcription start site (TSS) assay for profiling nuclear, capped RNAs, and PRIME, a computational framework for identifying active CREs from TSS data. Together, these methods increase sensitivity to low-abundance RNAs and enable robust detection of active regulatory elements across diverse contexts. Across multiple orthogonal functional and genetic benchmarks, including fine-mapped eQTLs, ClinVar variants, GWAS loci, and CRISPRi-tested elements, nucCAGE-derived PRIME predictions achieve superior recall compared to state-of-the-art methods while maintaining strong enrichment for phenotype-associated variation. Applying PRIME to the FANTOM5 dataset yields a comprehensive, cell-type-resolved atlas of active CREs that recapitulates known tissue-trait relationships. We demonstrate how this atlas can be used to nominate causal noncoding variants, linking immune-cell enhancer regulation of *SMAD3* to asthma and *NCOR2* to premature separation of placenta. Together, nucCAGE and PRIME provide a framework for high-sensitivity genome-wide discovery of active CREs and a resource for variant-to-function studies.

## Introduction

Cis-regulatory elements (CREs), including promoters and enhancers, ensure precise spatiotemporal activation of genes by integrating external signals with internal cell states^1–5^. CREs are essential for development and homeostasis, and disruptions caused by genetic variants in these elements are major contributors to human disease^6^. Yet reliably identifying causal CREs and the cell types in which variant effects manifest remains a fundamental challenge in genomics^7^, underscoring the need for sensitive methods for genome-wide mapping of CRE activity.

Large-scale epigenomic efforts have catalogued millions of putative CREs using chromatin accessibility and histone modifications^8,9^. These maps have been instrumental in linking noncoding genetic variants to CREs, yet epigenomic signatures may reflect permissive rather than functionally active chromatin states^10–12^. Consequently, identifying which candidate enhancers are active in a specific cellular context based on epigenomic data alone remains a major challenge^2^.

Transcription start site (TSS) assays, such as Cap Analysis of Gene Expression (CAGE)^13^, offer a complementary readout by enriching for the 5’ ends of m7G-capped RNAs originating from both promoters and enhancers, reflecting their shared architecture^2,14^ (**Supplementary Note 1**). Similar to mRNA gene promoters, which produce reverse strand promoter upstream transcripts (PROMPTs)^15,16^, active enhancers initiate divergent transcription of enhancer RNAs (eRNAs)^17–20^. This phenomenon enabled the construction of CAGE-derived enhancer atlases across human cell types and tissues in the FANTOM projects^17,21,22^.

Although CAGE-defined enhancers are relatively sparse (63,285 enhancers identified across 1,829 FANTOM5 libraries^23^), they exhibit disproportionately strong enrichment for complex trait heritability. For example, FANTOM5 enhancers show a 39-fold enrichment for immunological traits compared to a 3-fold enrichment for putative enhancers marked by H3K27ac^24^. While their total heritability contribution is constrained by their limited genomic coverage, this high concentration of signal indicates that transcription initiation provides a more specific and functionally refined readout of regulatory activity than chromatin signatures. Improving the sensitivity of TSS assays or computational strategies to identify a more comprehensive set of CREs from TSS data will substantially benefit variant interpretation by increasing the proportion of heritability that can be mapped to high-confidence CREs.

Unlike mRNAs, most eRNAs and PROMPTs lack poly(A) tails and are rapidly degraded^25–31^, limiting their detectability by TSS assays^32,33^. Nascent transcript assays, such as GRO-cap^34^, address the low abundance and instability of eRNAs^35^ by measuring transcriptional activity directly at DNA loci. However, these assays cannot distinguish polymerase pausing from re-initiation events^36–39^ and require substantial starting material^40^, which limits their utility. Furthermore, the degree to which TSS assays capture nascent transcripts remains unclear. TSS assays that enrich for nascent transcripts through nuclear fractionation^41^ or subcellular fractionation of chromatin^42^ (NET-CAGE) show promise. Recent ENCODE benchmarking datasets^43^ enable systematic evaluation of both experimental and computational methods for CRE mapping.

Here, we present nucCAGE (**Fig. 1A-B**), a TSS assay optimized for profiling nascent transcripts, together with PRIME (**Fig. 1C**), a computational framework for the accurate mapping of CREs from TSS data. Leveraging the shared transcriptional architecture of promoters and enhancers^2,14^, PRIME predicts CREs as a unified class from TSS data rather than explicitly distinguishing between these element types. Together, these methods overcome key limitations of existing approaches by improving the detection of low-abundance eRNAs and achieving higher recall and functional enrichment of CREs across diverse benchmarks relative to existing methods. We compare predictions from nucCAGE with those from whole-cell CAGE, assess the trade-off between their recall and precision, and derive optimal experimental parameters for CRE profiling. Finally, we demonstrate PRIME’s general versatility in detecting CREs from other TSS assay data by applying it to CAGE libraries from the FANTOM5^44^ consortium to build a greatly expanded atlas of CRE activity across 141 human cell types, tissues, organs, and cell lines, and demonstrate how this atlas can guide functional interpretation of noncoding genetic variants.

**Figure 1.**
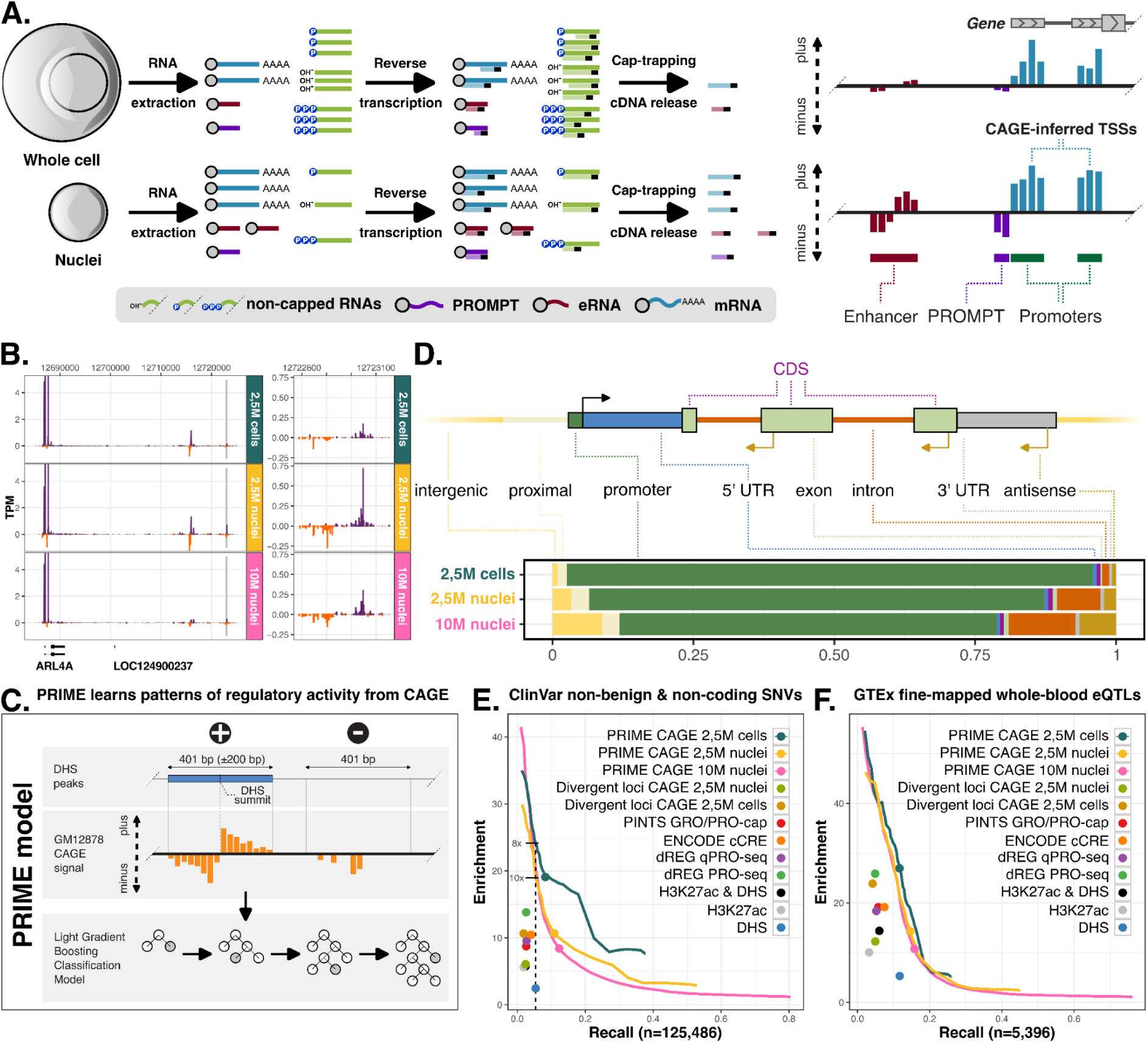
nucCAGE and PRIME increase the sensitivity and functional relevance of active CRE detection. **A**. Conceptual overview of the nucCAGE protocol. By extracting nuclei prior to cap-trapping, nucCAGE aims to deplete abundant, stable cytoplasmic RNA species (e.g., rRNAs) (left) and increase the sensitivity for unstable, nuclear-retained RNA species, such as eRNAs and PROMPTs (right). **B**. TPM normalized K562 CTSS signal (vertical axes) at the *ARL4A* promoter (left) and its known enhancer (right, marked in grey in the left panel), defined from CRISPRi^46^ and massively parallel reporter assay (MPRA)^47^ data. CTSS signal on the minus strand is displayed as negative TPM values. **C**. Schematic of the PRIME computational framework. A LightGBM-based classifier was trained to learn characteristic TSS signal patterns of active CREs in a data-driven manner, using GM12878 nuclei and whole-cell CTSS signals within DHSs as positive training examples (proxies for active CREs), excluding annotated exonic regions, and CTSS signals distal to DHSs as negative training examples. **D.** Genomic distribution of identified TSSs, visualized as stacked bars reflecting the proportion of detected TSSs across annotation categories. Shown for pooled K562 2.5 million (M) whole-cell samples and 2.5M and 10M nuclei samples. For concordance across replicates, see **Fig. S2B**. **E**. Enrichment-recall curves for predicting pathogenic and likely pathogenic ClinVar single nucleotide variants (SNVs) as well as variants of unknown significance. Enrichment is defined as the ratio of noncoding, non-benign ClinVar SNVs to noncoding common variants overlapping predicted CREs. Recall is the fraction of noncoding, non-benign ClinVar SNVs overlapping predicted CREs. Curves represent performance at thresholds corresponding to recall increments of 0.01; points indicate performance at recommended thresholds. The dashed vertical line illustrates the relative increase in enrichment of PRIME predictions compared to DHSs. For results stratified by promoter distance, see **Fig. S3A**. **F.** Enrichment-recall curves for predicting fine-mapped eQTLs. Enrichment is defined as the ratio of fine-mapped noncoding eQTLs (PIP > 50%) to noncoding common variants overlapping predicted CREs. Recall is the fraction of eQTL variants overlapping CREs. Curves and points are defined as in **E**. For results stratified by promoter distance, see **Fig. S3B**.

## Results

### nucCAGE and PRIME increase the sensitivity of CRE detection

To improve the detection of active CREs, we developed nucCAGE, a TSS assay that enriches eRNAs and other unstable, nuclear-retained transcripts by selectively extracting nuclei prior to cap-trapping (**Fig. 1A**). While cellular subfractionation was used in combination with CAGE library preparation before^42,45^, we optimised nuclei extraction by adding controlled mechanical shear forces (see Methods) to detach the endoplasmic reticulum from the nuclear envelope, reducing the carryover of mature ribosomal RNAs (rRNAs) (**Fig. S1**). Because abundant cytoplasmic RNAs, such as rRNAs, occupy resources during reverse transcription, their depletion increases the recovery of capped RNAs with complete second-strand cDNAs. This increases the relative proportion of unstable, non-polyadenylated transcripts, including eRNAs in the nucCAGE libraries^19,42^. Consistent with this, nucCAGE libraries from K562 cells exhibited stronger signal at CAGE-derived transcription start sites (CTSSs) than whole-cell CAGE at known enhancer loci and PROMPTs, while maintaining comparable CTSS signal at mRNA promoters (**Fig. 1B, Fig. S2A**).

Across samples and replicates, nucCAGE increased the relative fraction of CTSSs mapping to intronic and intergenic regions, which contain putative enhancers, while retaining the expected enrichment of mRNA-associated TSSs (**Fig. 1D**, **Fig. S2B**). This reflects improved detection of unstable RNAs relative to whole-cell samples, as exemplified by increased CTSS signal from eRNAs at the known *ARL4A* enhancer^46,47^ (**Fig. 1B**) and at the PROMPT upstream of the *KLF2* promoter (**Fig. S2A**). Furthermore, nucCAGE libraries displayed a high proportion of CTSSs near DNase I hypersensitive sites (DHSs)^9^, mirroring whole-cell CAGE (**Fig. S2C**) and consistent with the expected chromatin accessibility of active CREs. Together, these results demonstrate that nucCAGE enriches for unstable RNAs and increases the sensitivity of active CRE mapping.

In parallel with nucCAGE development, we sought to improve the computational prediction of active CREs. We developed PRIME, a LightGBM-based classifier that predicts active CREs directly from TSS signal profiles (**Fig. 1C**) along with a computational framework for analyzing TSS assay data. The PRIME model was trained to learn characteristic TSS signal patterns derived from GM12878 CAGE data that are indicative of active CREs, using DHSs as a proxy for regulatory activity, similar to previous work^48–50^. The model does not explicitly distinguish between promoter and enhancer activity, thereby circumventing strictly defined heuristic properties of CREs, such as balanced divergent transcription^17^, which are sensitive to library quality and sequencing depth.

We benchmarked the GM12878-derived PRIME model by applying it to K562 cells and comparing predictions against existing CRE detection methods based on CAGE-derived divergent transcription (FANTOM5-style^17^ divergent loci), nascent transcription^35,48^ (dREG, PINTS), histone modifications^8,51^ (H3K27ac, ENCODE cCREs), or DHSs^9^ alone (all in K562 cells). PRIME achieved substantially higher recall and enrichment of ClinVar^52^ noncoding, pathogenic variants (pathogenic, likely pathogenic) and variants of unknown significance (VUSs) relative to common variants (**Fig. 1E**), regardless of gene promoter distance (**Fig. S3A**), providing a measure of functional relevance through disease-associated variation. For example, at a PRIME score threshold chosen to match the highest recall achieved by alternative methods (0.053 for DHSs), PRIME achieved 19-fold and 24-fold enrichment of ClinVar variants when applied to nucCAGE and whole-cell CAGE, respectively, outperforming all other methods (corresponding to 8-fold and 10-fold higher enrichment than DHSs, respectively; **Fig. 1E**, dashed vertical line). These findings were corroborated using fine-mapped eQTLs from GTEx^53^, which provide an independent benchmark linking regulatory activity to gene expression, in which PRIME again showed superior recall and enrichment (**Fig. 1F**, **Fig. S3B**). We note that, although nucCAGE-based predictions provide higher recall than those generated from whole-cell CAGE, the increased sensitivity is accompanied by a slight reduction in enrichment.

In conclusion, PRIME applied to nucCAGE libraries delivers the highest sensitivity of all methods tested, enabling more comprehensive detection of active enhancers and improved interpretation of noncoding variants.

### Library complexity delineates CRE detection capacity

Given the increased sensitivity of nucCAGE for detecting active CREs, we next sought to define the limits of library complexity and the upper bound of CREs detectable by TSS assays. Using GM12878 cells, we evaluated the influence of input material on nucCAGE performance.

As expected, nuclear RNA extraction yielded substantially less RNA than whole-cell equivalents (3.5–13-fold reduction; **Fig. 2A**, **Fig. S4A**). This decrease reflects both the intentional removal of abundant cytoplasmic RNAs and a ∼50% loss of nuclei during extraction (**Fig. S4B-C**). Because CAGE requires 5 µg of input RNA for optimal library construction to ensure proper scaling of the reverse-transcription reaction and efficient synthesis of second strand cDNA^13,54,55^, carrier RNA is required when starting materials fall below this range^56^. Consequently, two opposing factors shape nucCAGE library complexity: reduced RNA input lowers detectable complexity, while depletion of abundant cytoplasmic RNAs increases the relative representation of transient, nascent transcripts. To account for these counteracting influences, we compared GM12878 samples from 500,000 cells with 1 million nuclei (compensating for nuclei loss) and 5 million nuclei (compensating for loss of total RNA). We also included 1 million whole-cell and 10 million nuclei samples for reference.

**Figure 2.**
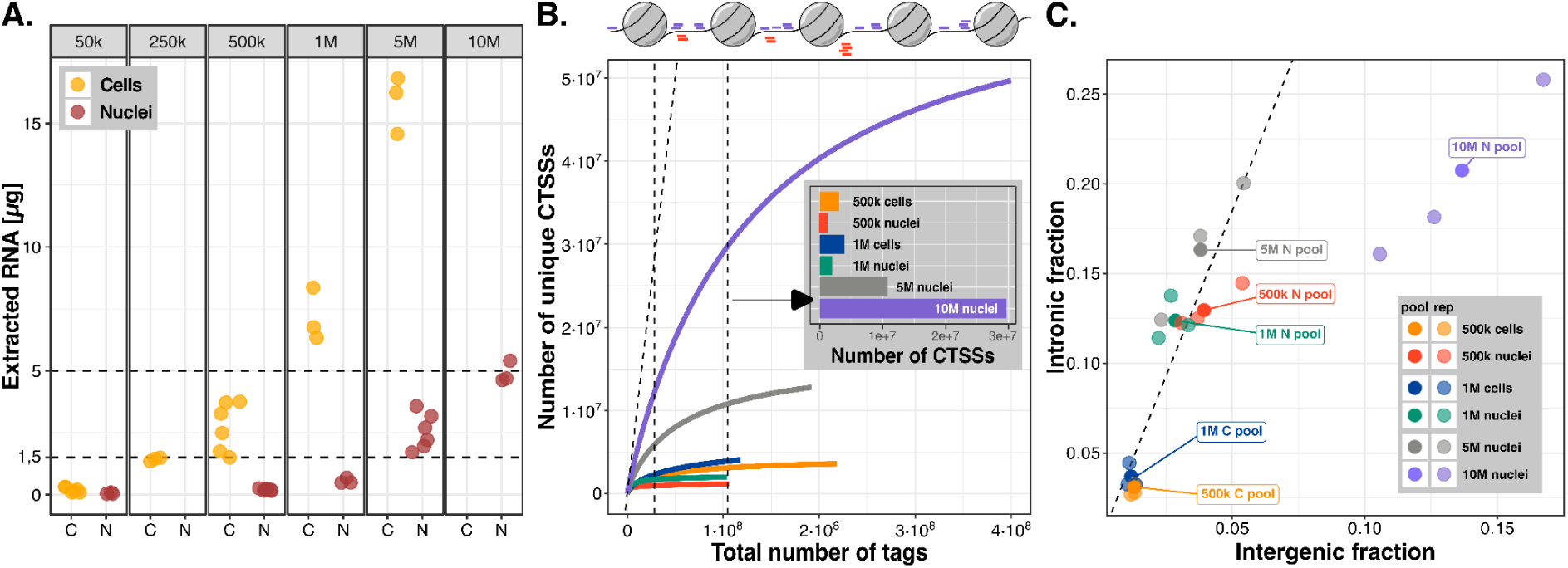
Library complexity and noise estimates define optimal CAGE input parameters for CRE detection. **A.** Total RNA yield (vertical axes) from whole-cell (C, yellow) and nuclei (N, red) extractions, stratified by GM12878 input cell number (horizontal panels). Each dot represents a separate RNA extraction experiment. **B.** Library complexity curves for pooled samples, showing the number of unique CTSSs against the total number of CAGE reads (library size) to estimate the transcriptional diversity and signal-to-noise ratio. **C.** Fractions of intergenic versus intronic CTSSs for pooled samples and individual replicates. All samples exhibit a linear, scaled increase in these fractions commensurate with input material, with the exception of the 10 million nuclei samples, which deviate from this trend.

As expected, higher input cell numbers increased the number of unique CTSSs detected, and whole-cell libraries showed greater complexity than nucCAGE libraries at matched input cell numbers (e.g., 1 million cells vs. 1 million nuclei; **Fig. 2B, Fig. S5A-C**), while showing similar noise levels (**Fig. S5D**). Notably, increasing input to 5 million nuclei markedly increased the number of genomic bins with CTSS coverage (**Fig. S5A**). Further increasing input to 10 million nuclei while keeping all reagent concentrations resulted in substantial noise accumulation (**Fig. S5D**). This was accompanied by clear deviations in the proportions of intergenic, intronic, and exonic CTSSs, with intronic fractions growing at the expense of exonic signal and intergenic fractions increasing at the expense of intronic signal (**Fig. 2E, Fig. S5E**). This is indicative of an accumulation of spurious reads at the expense of true TSS signals, likely driven by less effective cap-trapping of capped RNAs in crowded reactions. Similar patterns were observed in 10 million K562 nuclei samples (**Fig. 1C**, **Fig. S2B**).

Consistent with these observations, nucCAGE input affected the number of detected CTSS-associated DHSs (**Fig. S6A**). At lower inputs (500,000 and 1 million nuclei), nucCAGE identified fewer CTSS-associated DHSs in intronic and intergenic regions than whole-cell CAGE. In contrast, increasing input to 5 million nuclei significantly increased the number of expressed DHSs, approaching or exceeding the levels identified by GRO-cap.

We next investigated the CTSS complexity of CAGE-inferred core promoters (also known as “tag clusters”), derived from positional clusters of same-strand CTSSs^57^. Low-input nucCAGE samples (500,000 and 1 million nuclei), showed (similarly to GRO-cap samples) lower CTSS positional complexity, with reduced CTSS positional distribution width (10-90% IQR) and shape entropy^44,57^ (**Fig. S6B**), while 5 million nucCAGE samples closely resembled the positional complexities of 500,000 and 1 million whole-cell CAGE libraries. Of note, 5 million and 10 million nucCAGE samples also displayed a higher expression of canonical^58^ (YC) initiator-associated TSSs than low-input samples (**Fig. S6C**), suggesting that higher promoter TSS complexity is not simply driven by increased noise.

Taken together, our results indicate that more than 1 million but fewer than 10 million GM12878 nuclei, corresponding to approximately 1.5–2.5 µg nuclear RNA (**Fig. 2A**), achieve a favorable balance between library complexity and noise. Within this range, nucCAGE displays high TSS complexity and achieves markedly greater complexity than 500,000 whole-cell samples, even though they originate from similar amounts of RNA starting material.

### nucCAGE identifies both nascent and steady-state transcripts

The increased complexity in nucCAGE libraries and enrichment for unstable, short-lived RNAs prompted us to investigate whether nuclei extraction enriches for nascent transcripts originating from CREs. We focused on CAGE-derived core promoters classified by their overlap with gene-proximal or gene-distal DHSs.

As expected (**Fig. 2**), a greater number of gene-distal, CAGE-inferred core promoters showed markedly higher expression (|log_2_ fold-change| > 1) in nucCAGE than in whole-cell CAGE (**Fig. 3A-B**). Unexpectedly, gene-proximal DHSs showed a similar pattern, exemplified by the *RAPGEF2* promoter, which displayed substantially increased CTSS signal in nucCAGE samples (**Fig. 3A**, horizontal axis in **Fig. 3B**). Such increases in promoter-associated CTSSs suggest that many transcription initiation events do not result in stable, mature mRNAs^25,59,60^.

**Figure 3.**
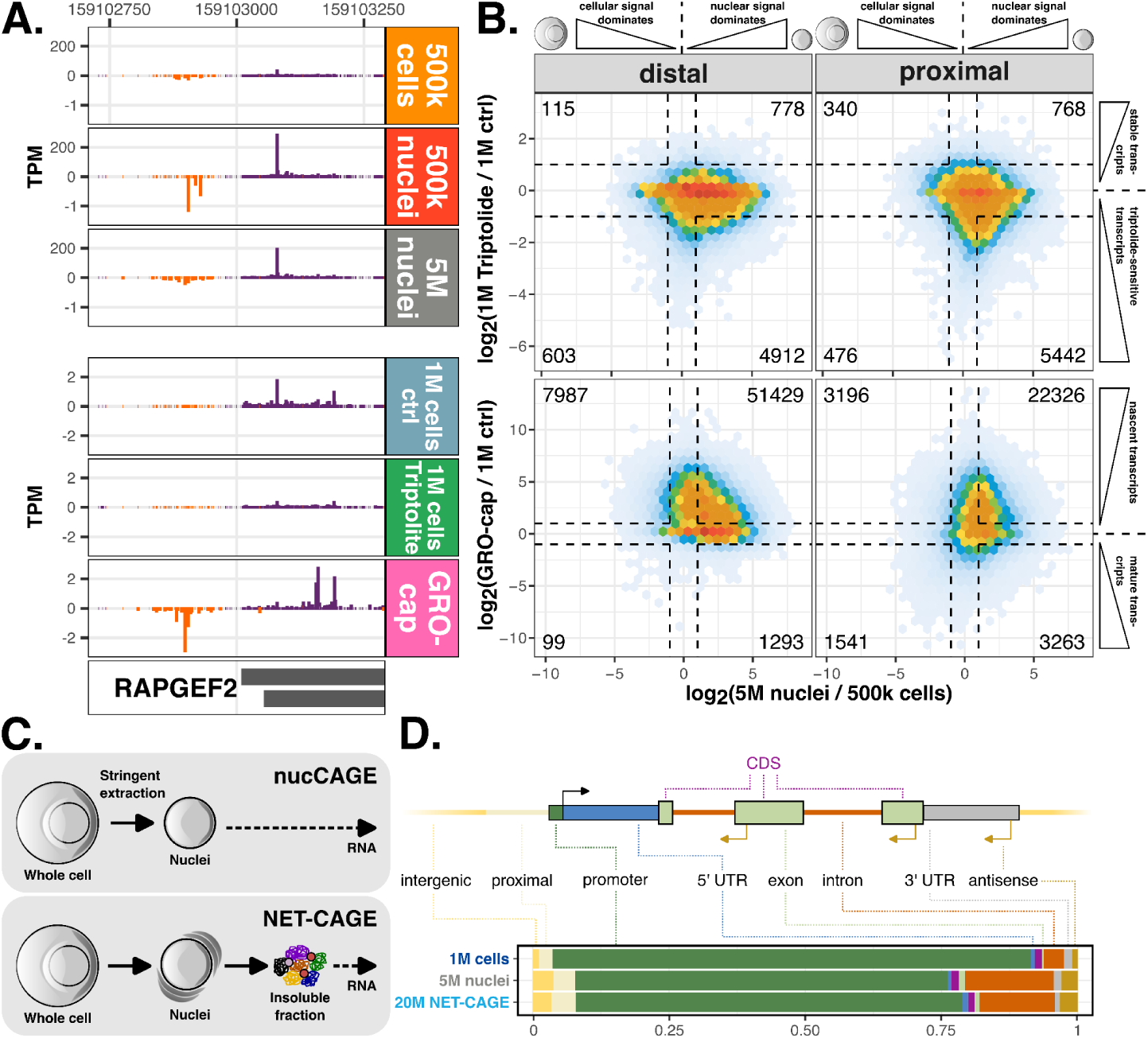
nucCAGE enables high-sensitivity detection of nascent transcripts. **A.** Genome tracks for nucCAGE, GRO-cap, and whole cell CAGE samples with or without triptolide treatment and varying GM12878 cell inputs, exemplifying signal enrichment across assays at the *RAPGEF2* promoter. Note the different TPM scales (vertical axes) to highlight PROMPTs and triptolide-induced signal depletion. **B.** 2D density plots contrasting enrichment of nuclear over whole-cell RNAs (horizontal axes) with the effects of triptolide (vertical axis, top row) and nascent RNAs captured by GRO-cap (vertical axis, bottom row). Numbers in plot corners represent the number of transcripts falling into the quadrants separated by dashed lines. Dashed lines represent log_2_ fold-changes of −1 and 1. **C.** Schematic illustration of the key methodological differences between NET-CAGE, which utilizes chromatin fractionation of nuclear RNA, and nucCAGE, which employs shear forces to perform stringent nuclei extraction. **D.** Genomic distribution of identified TSSs, visualized as stacked bars reflecting the proportion of detected TSSs across annotation categories. Shown for pooled GM12878 1 million (1M) whole-cell and 5M nuclei CAGE samples, and 20M NET-CAGE samples.

As a complementary test of nascent RNA enrichment, we assessed the sensitivity of nucCAGE to transcriptional arrest using the TFIIH inhibitor triptolide. Triptolide blocks the ATP-dependent helicase activity of TFIIH^61–63^, preventing transcription initiation and rapidly reducing nascent transcripts such as eRNAs and PROMPTs^27,28^. Both proximal and distal DHSs showed significantly reduced CAGE signal after triptolide treatment, with the strongest reduction observed for transcripts enriched in nucCAGE (**Fig. 3B**, top; p=0.024 for distal, p=9.12·10^-1^^72^ for proximal, χ^2^-test). This supports an increased sensitivity of nucCAGE to nascent transcripts.

We next benchmarked nucCAGE against GRO-cap, a nascent transcription assay. nucCAGE-enriched core promoters located in DHSs exhibited higher activity in GRO-cap than in whole-cell CAGE (**Fig. 3B**, bottom; p=8.15·10^-12^ for distal, p=9.59·10^-297^ for proximal, χ^2^-test), further validating the enrichment of nascent RNAs in nucCAGE over whole-cell CAGE. In agreement, nucCAGE exhibited comparable enrichment of CTSSs to NET-CAGE^42^ within intronic and intergenic regions (**Fig. 3C-D**), consistent with the expected genomic locations of nascent eRNAs and other long non-coding RNAs. However, nucCAGE requires considerably less input material and does not rely on transcriptional inhibitors such as α-amanitin. Of note, many nucCAGE-derived core promoters in distal DHSs were not detected by GRO-cap, suggesting that nucCAGE can capture transient or low-abundance initiation events that other nascent assays may miss.

To assess the detection of steady-state transcripts, we compared gene-associated CTSS expression with RNA-seq-derived gene expression^51^. Whole-cell CAGE samples showed strong correlations with RNA-seq (Pearson’s r = 0.81), while nucCAGE exhibited slightly reduced correlations (r = 0.72) (**Fig. S7**), consistent with its enrichment of unstable RNAs. Notably, nucCAGE libraries prepared from 5 million nuclei showed the highest correspondence with RNA-seq (r = 0.73), reflecting reduced noise and optimal sensitivity at this input level (**Fig. S5D**). GRO-cap, as expected for a nascent assay, showed the lowest correlation with RNA-seq among tested methods (r = 0.68).

In summary, nucCAGE captures both nascent and stable transcripts, enabling the detection of more putative enhancers and promoters than whole-cell CAGE. Using approximately 2.5–5 million nuclei provides an optimal input range that balances high sensitivity with low noise, yielding high promoter complexity, high detection of nascent transcripts, and RNA-seq correlations that exceed those of nascent transcript assays.

### PRIME performs data-driven computational detection of CREs from TSS data

CAGE has proven powerful for characterizing CREs by capturing transcription initiation with base-pair resolution^17,44,58^, and nucCAGE further increases TSS complexity (**Fig. 2**) and sensitivity to nascent transcripts (**Fig. 3**). However, defining active CREs *de novo* from TSS data typically relies on heuristic thresholds, such as the overall strand directionality of divergently transcribed loci^17^ or minimum expression cutoffs, which are highly sensitive to library complexity (**Fig. S8A**). As a result, stringent cutoffs can discard biologically relevant genomic loci, whereas lenient thresholds risk incorporating noise (**Fig. S8B**). Consistently, although stringent criteria for divergent transcription identify highly relevant loci^14^, they can fail to capture the continuum of promoter and enhancer activities represented in CAGE profiles (**Fig. 1E-F**). These limitations highlight the need for a data-driven method that can learn features of genuine regulatory activity directly from high-resolution TSS data.

We present PRIME, a LightGBM-based classifier designed to overcome these limitations by predicting active CREs directly from transcription initiation profiles (**Fig. 1C**). Although we trained PRIME specifically on CAGE CTSS signals, the resulting model is assay-agnostic and can be applied to any TSS data with base-pair resolution. Here, we focus on its application to CAGE and nucCAGE datasets. PRIME was trained to learn characteristic TSS signal patterns indicative of CREs, normalized to emphasize signal shape rather than absolute expression levels, using DHSs with detectable CTSS signal as positive examples, excluding annotated exonic regions, and genomic regions with CTSS signal outside of DHSs as negatives (**Fig. S9**, see Methods). PRIME was trained on GM12878 whole-cell and nucCAGE datasets across multiple input levels. Importantly, this training strategy enabled the model to learn characteristic initiation patterns of *bona fide* CREs independently of thresholding criteria or sample-specific TSS complexities, without explicitly distinguishing between promoter and enhancer activity, using DHS overlap not as ground truth but as a practical scaffold to expose the model to representative TSS signal patterns of regulatory activity.

For genome-wide prediction, PRIME evaluates sliding windows across CTSS-dense genomic loci, assigning prediction scores, which are subsequently thresholded and merged into contiguous loci, each with a single integrated PRIME (**Fig. S10**, see Methods). This prediction strategy significantly increases recall of CREs, while maintaining precision comparable to divergent loci defined by transcriptional directionality (**Fig. S11**). PRIME prediction scores (range: [0,1]) represent model-specific confidence measures and should not be interpreted as calibrated probabilities.

To interpret what PRIME had learned, we analyzed SHAP-based^64^ feature contributions across predicted CREs. SHAP values highlight the positional influence of TSS signal within each site, with the strongest contributions concentrated within the first ∼100 bp around the PRIME locus center (**Fig. 4A**), regardless of genomic location (**Fig. S12**). This mirrors the characteristic initiation patterns of active promoters and enhancers^17,65–67^, indicating that PRIME captures biologically meaningful initiation patterns that generalize across different TSS profiling methods.

**Figure 4.**
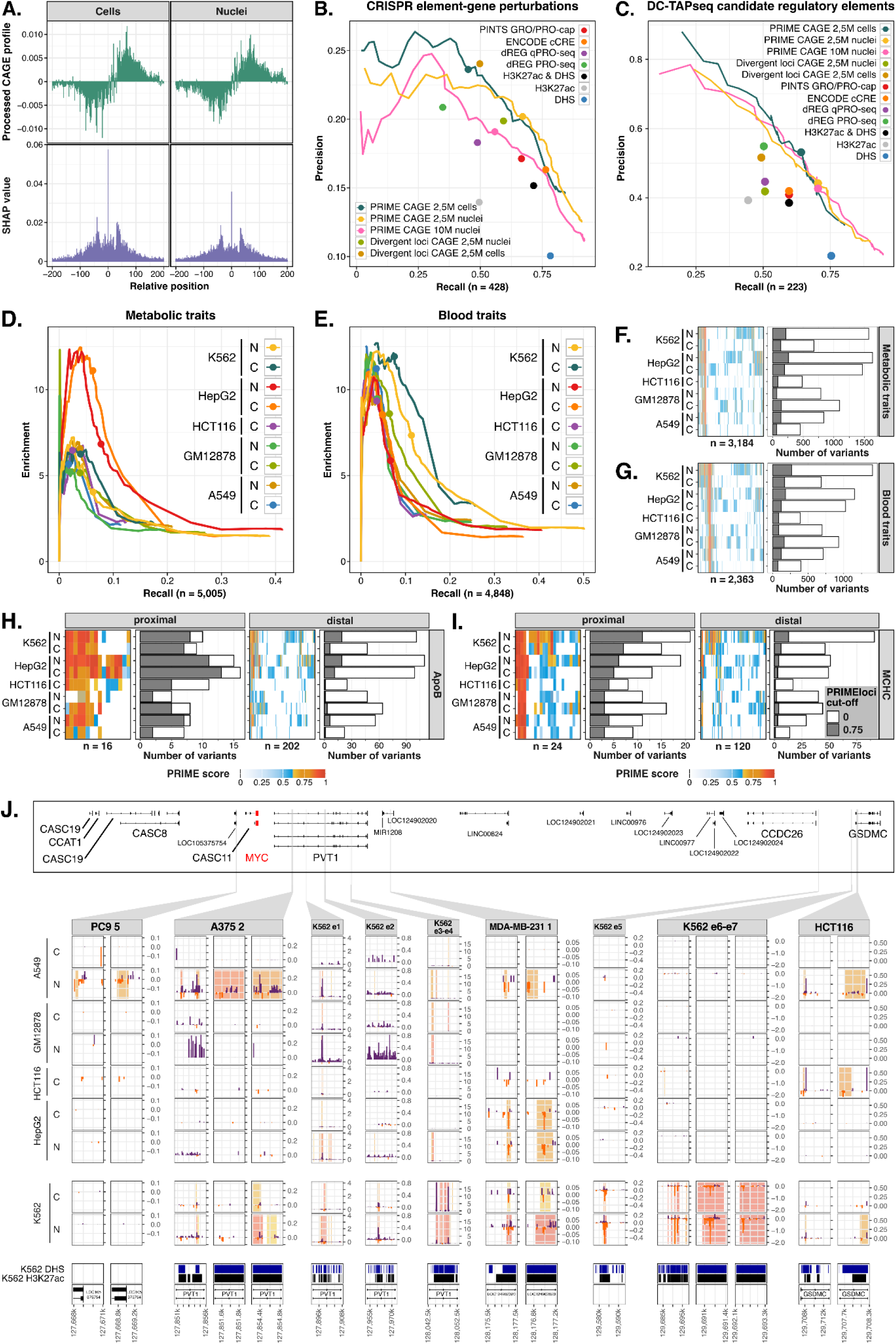
Functional validation and trait-association mapping of predicted CREs. **A.** Positional importance of transcription initiation. Histograms showing the average CTSS signal (top, positive values: plus strand, negative values: minus strand) and SHAP values (bottom) at base pairs relative to the PRIME locus center, illustrating the contribution of CTSS signal and position to PRIME classification. **B.** Benchmarking against CRISPRi screens. Precision-recall curves for PRIME and alternative CRE prediction methods (H3K27ac, DHS, FANTOM5-style divergent loci, ENCODE cCREs, dREG, and PINTS) evaluated against gold-standard CRISPRi-validated enhancers in K562 cells. Curves represent continuous predictors, while dots indicate binary predictors. **C.** Validation using DC-TAP-seq. Precision-recall curves for the same methods as in **B**, but evaluated against DC-TAP-seq data as the ground truth. **D.** Enrichment of metabolic trait-associated variants (see Methods). Enrichment-recall curves showing the enrichment of fine-mapped GWAS variants for metabolic traits relative to common variants as a function of recall across predicted CREs in K562, HepG2, HCT116, GM12878, and A549 nuclei (N) and cell (C) samples. **E.** Enrichment of blood trait-associated variants. Similar to **D**, but for blood trait variants (see Methods). **F.** Corresponding plot to **D**, displaying the specific cell types where predicted CREs overlap individual variants associated with metabolic traits. The heatmap displays the PRIME scores associated with each variant, while the bar plot displays the number of variants captured by PRIME loci in each cell type sample at no PRIME score threshold (white) and a PRIME score threshold of 0.75 (grey). **G.** Corresponding plot to **E**, displaying the specific cell types where predicted CREs overlap individual variants associated with blood traits. Heatmap and bar plot as in **F**. **H.** Mapping of ApoB-associated variants. Analysis of ApoB-associated variants stratified by whether the overlapping PRIME-derived CRE is located proximal or distal to annotated TSSs. Heatmaps and bar plots as in **F**. **I.** Mapping of MCHC-associated variants. Analysis of MCHC-associated variants stratified by whether the overlapping PRIME-derived CRE is located proximal or distal to annotated TSSs. Heatmaps and bar plots as in **F**. **J.** Regulatory architecture of the *MYC* gene locus showing zoom-ins of known CREs to *MYC*. CAGE CTSS signal tracks (TPM-normalized) are displayed for K562, HepG2, HCT116, GM12878, and A549 nuclei (N) and cell (C) samples and are overlaid with PRIME CRE predictions (vertical bars) colored by PRIME prediction score. For several CREs, we provide additional zoom-ins, illustrating the positional refinement of CREs achieved by PRIME. See **Supplementary** Figure 14 for a detailed overview.

### Benchmarking PRIME for CRE detection against CRISPRi perturbations

Encouraged by the variant-level benchmark (**Fig. 1E-F**), we next evaluated PRIME using a gold-standard dataset of CRISPRi-tested genomic elements from perturbation screens in K562 cells^43^. We compared PRIME to widely used CRE detection methods, including H3K27ac peaks^51^, DHSs, ENCODE cCRE^8^, and nascent transcript data-based computational methods dREG^49^ and PINTS^35^.

While chromatin-based approaches (H3K27ac, DHSs, and their combination) achieved high recall, they identified large numbers of CRISPRi-negative regions (**Fig. S13A**), resulting in low precision (**Fig. 4B**). In contrast, RNA-based methods, and PRIME in particular, substantially improved precision while maintaining high recall. Across the entire range of PRIME score thresholds, the model consistently outperformed all alternative methods, demonstrating a superior balance between precision and recall.

Although PRIME predictions derived from nucCAGE showed slightly lower precision than those derived from whole-cell CAGE at equivalent score thresholds, they exhibited markedly higher recall. Notably, many PRIME-predicted positive regions that were found negative by CRISPRi data were supported by alternative prediction methods (**Fig. S13B**), suggesting that these sites may represent functional CREs not detected in the K562 CRISPRi screens. Indeed, when benchmarking against an independent CRISPRi dataset with greater power to detect perturbations of smaller effect sizes^68^, PRIME achieved even higher precision in K562 cells (**Fig. 4C**), further supporting the biological relevance of PRIME predictions.

Together, PRIME achieved the best overall balance between enrichment and recall across all considered benchmarks (eQTL, ClinVar, CRISPRi; **Fig. 1E-F**, **Fig. 4B-C**).

### PRIME discovers cell type-specific trait enrichments

To assess PRIME’s ability to identify cell-type-specific CRE activity, we generated nucCAGE and whole-cell CAGE datasets from GM12878 (lymphoblast), K562 (myelogenous leukemia), A549 (lung adenocarcinoma), HepG2 (hepatoblastoma), and HCT116 (colon carcinoma) cell lines. Enrichment analysis of trait-associated SNVs confirmed the cell-type relevance of PRIME predictions on these datasets: metabolic and liver-related traits were most enriched in PRIME loci identified in HepG2 cells (**Fig. 4D,F,H**), while immune- and blood-trait SNVs showed the strongest enrichment in K562 cells (**Fig. 4E,G,I**).

Interestingly, we also observed enrichments across cell types suggestive of shared regulatory functions. For example, SNVs associated with Apolipoprotein B (ApoB) were enriched not only in HepG2 but also in K562 PRIME loci, including a candidate CRE identified in the first intron of the *ABO* gene (**Supplementary Note 2**, **Fig. SN1**). Conversely, a PRIME-inferred CRE within *ABCA7* (**Supplementary Note 2**, **Fig. SN2**), overlapping an SNV associated with mean corpuscular hemoglobin concentration (MCHC), was active in HepG2 cells despite the erythroid nature of the trait.

To further dissect cell-type-specific regulatory activity of disease relevance, we next examined CRE architecture across multiple cell types at the *MYC* locus (**Fig. 4J**, **Fig. S14**). PRIME recapitulated previously validated CREs while providing higher-resolution localization of regulatory elements. In K562 cells, we observed strong and selective activation of multiple CRISPRi-validated enhancers (e1–e7)^69^. Notably, elements e3 and e4 exhibited promoter-like initiation profiles characterized by strong, plus strand-biased transcription. In addition, we observed a cluster of PRIME CREs in K562 nuclei samples intragenic to *CCDC26*, co-localizing with the previously characterized blood enhancer cluster (BENC) locus^70^ (**Fig. S14**). Beyond hematopoietic contexts, nucCAGE and PRIME identified distinct patterns of epithelial CRE activity, including A549-specific expression of PC9-5, A549-biased activity of A375-2, and shared activity of HCT116-1 in A549 and HCT116 cells^71,72^. In contrast, MDA-MB-231-1 (overlapping MYC Lung Adenocarcinoma Super-Enhancer (MYC-LASE)^73^), exhibited broad activity across A549, HCT116, HepG2, and K562 cells, but remained inactive in GM12878.

Together, these results indicate that the combination of nucCAGE and PRIME provides the sensitivity and specificity required for detailed dissection of regulatory architecture at disease-associated loci.

### Building an atlas of CRE activities across human cell types and tissues

Given PRIME’s strong enrichment, recall, and cell-type resolution for variant-linked regulatory elements (**Fig. 1E-F**, **Fig. 4**), we next applied the GM12878-trained PRIME model at atlas-scale to the extensive CAGE datasets generated by the FANTOM5 consortium^44^. To derive a robust, genome-wide map of CRE activity across the atlas, we applied PRIME to the aggregated signal across a curated set of 760 FANTOM5 CAGE libraries (**Fig. 5A**, see Methods). A total of 2,849,385 regions (putative enhancers and promoters) with PRIME scores ≥ 0.75 were retained as cell-type agnostic focal points for downstream analyses. Then, for each of 141 considered cell types, tissues, and cell lines, herein referred to as ‘facets’ (ontology-derived groups of CAGE libraries, see Methods, **Supplementary Table 1**), we computed facet-specific PRIME scores within this fixed set of loci, enabling fine-grained evaluation of CRE activity across diverse cell types and tissues. Hierarchical clustering based on facet-specific PRIME scores revealed a clear organization within the atlas, including a separation between physiological tissues and cancer-derived samples (**Fig. 5B**, **Fig. S15**).

**Figure 5.**
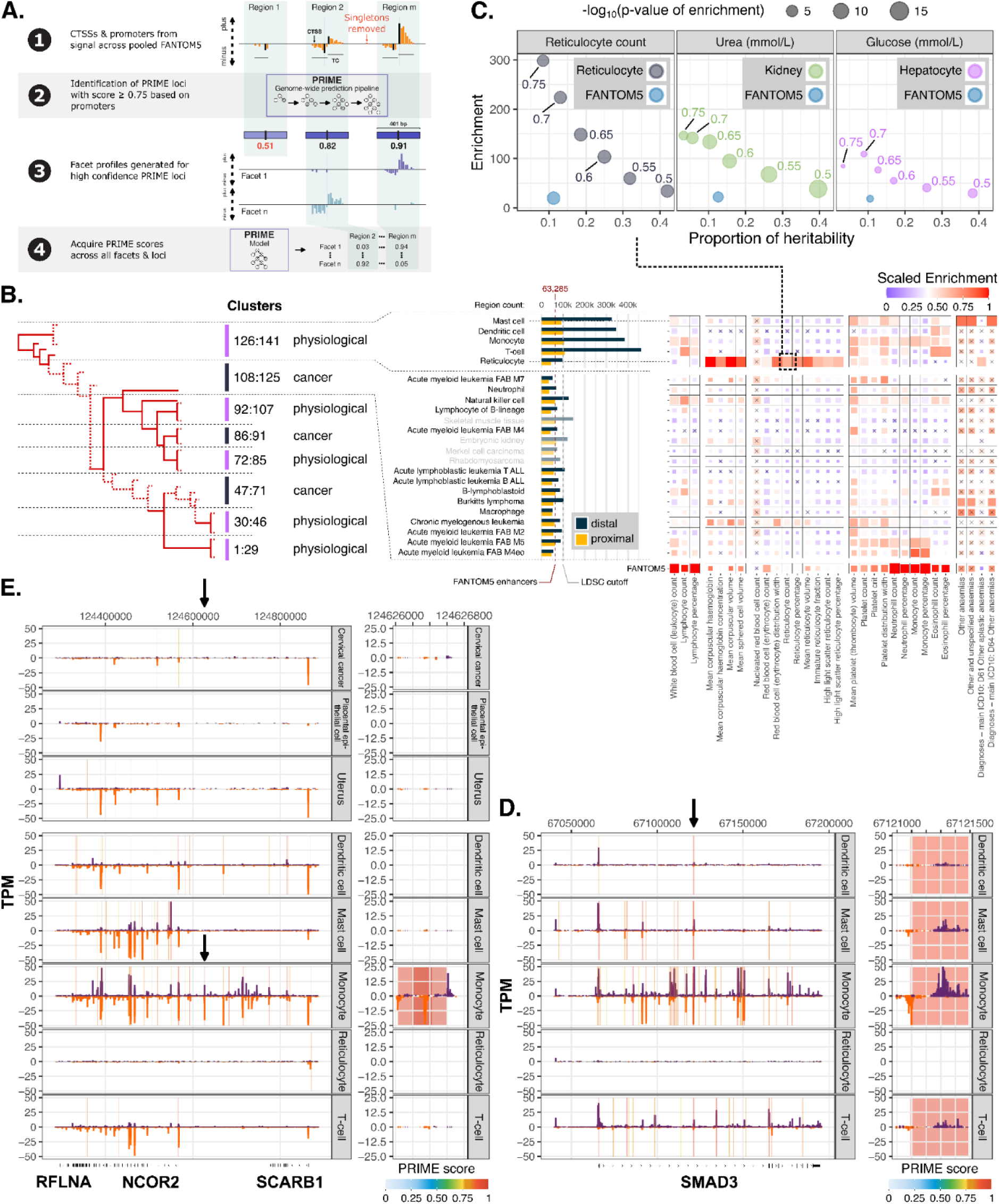
A PRIME-enhanced atlas of CRE activities reveals trait-cell-type associations. **A.** Schematic overview of the strategy for applying PRIME to 760 FANTOM5 CAGE libraries to construct an atlas of CREs across 141 cell, tissue, and cell line facets. **B.** Left: hierarchical clustering based on facet-specific PRIME scores revealing the organization of the atlas, including a separation between physiological and cancer-derived samples. Middle: barplot illustrating the number of distal and proximal CREs identified for each facet, at a minimum PRIME score threshold of 0.5. The number of enhancers identified in FANTOM5 and the maximum number of distal CREs considered for stratified LD score regression (s-LDSC) are depicted by red and grey dashed lines, respectively. Right: scaled heritability enrichment from s-LDSC for blood-related traits. Non-significant enrichments (nominal P value > 0.05) are marked with crosses. **C.** s-LDSC-derived enrichment (vertical axis) versus proportion of heritability (horizontal axis) for reticulocyte counts, urea, and glucose traits at different PRIME score thresholds in reticulocyte, kidney, and hepatocyte facets, respectively. Corresponding values for FANTOM5 enhancers are shown for reference. **D.** Regulatory architecture of the *SMAD3* gene locus, showing an intronic distal CRE (chr15:67121091-67121491, marked by black arrow, zoomed in to the right) with high PRIME activity in immune cell types. This CRE overlaps the variant rs17294280 (15_67121286_G_A), which is strongly associated with asthma, lung function (FEV/FVC ratio), and allergic rhinitis (see **Supplementary Note 3**, **Fig. SN3** for details). CAGE CTSS signal tracks (TPM-normalized) are displayed for FANTOM5 facets and are overlaid with PRIME CRE predictions (vertical bars) colored by PRIME prediction score. **E.** Regulatory architecture of the *NCOR2* gene locus, highlighting a monocyte-active distal CRE (chr12:124626205-124626605, marked by black arrow, zoomed in to the right). This CRE overlaps the variant rs763017549 (12_124626281_C_T), which is associated with premature separation of placenta (abruptio placentae) (see **Supplementary Note 3**, **Fig. SN4** for details). CAGE CTSS signal tracks (TPM-normalized) are displayed for FANTOM5 facets and are overlaid with PRIME CRE predictions (vertical bars) colored by PRIME prediction score.

Due to the increased sensitivity of PRIME over divergent loci calling, the model substantially increased the number of identified distal CREs in FANTOM5 samples. While the original FANTOM5 enhancer atlas contains 63,285 predicted enhancers^17,23^, PRIME identified a median of 73,151 promoter-distal CREs per facet at a PRIME score threshold of 0.5 (**Fig. 5B**, **Fig. S15, Supplementary Table 2**). Specifically, 2,450,372 (2,146,690 promoter-distal and 303,682 promoter-proximal) and 399,875 (263,685 promoter-distal and 136,190 promoter-proximal) of the ∼2.9 million cell-type agnostic PRIME loci were identified in at least one facet at a PRIME score threshold of 0.5 and 0.75, respectively (**Supplementary Data,** https://anderssonlab.org/PRIMEatlas/). Many PRIME-predicted loci were detected in only one or a few cell types, indicating a substantial increase in cell-type specificity relative to the original atlas (**Fig. S16**).

To evaluate their functional relevance, we performed stratified LD score regression (s-LDSC)^24^ across 4,178 UK Biobank traits^74,75^. For each facet, associated gene-distal PRIME loci were assessed for enrichment of trait heritability. PRIME-derived enrichments reliably recapitulated known relationships between traits and cell types and tissues, and revealed cell-type specific enrichments beyond those detectable using the original FANTOM5 enhancer set (**Fig. S17**). For example, we observed heritability enrichments for urea, C-reactive protein, and urate levels in kidney distal CREs; blood pressure in cardiac fibroblast distal CREs; spherical power in eye distal CREs; and various neurological traits in brain and neuronal distal CREs.

As previously noted^24^, the original FANTOM5 enhancer atlas is highly enriched for blood-and immune-related traits (**Fig. S17**, **Fig. 5B**), likely due to an overrepresentation of hematopoietic cell types within the dataset. Importantly, PRIME recapitulated these enrichments while providing far greater resolution across related immune cell types (**Fig. 5B**). Systematic s-LDSC analysis across multiple PRIME score thresholds further demonstrated that lenient thresholds (e.g., 0.5) capture broad sets of trait-associated distal CREs with reduced enrichment, whereas stricter thresholds (e.g., 0.75) trade recall for increased confidence, enabling tuning of the predictions to a desired balance of recall and precision (**Fig. 5C)**.

Among blood-related traits, PRIME correctly attributed lymphocyte-associated traits to T cell, natural killer cell, and lymphocyte cell facets; hemoglobin- and reticulocyte-related traits to reticulocyte facets; and monocyte traits to the monocyte cell facet as well as monocyte-related acute myeloid leukemia subtypes M4eo and M5 (**Fig. 5B**). More complex relationships between immune traits and immune cell types were also evident. For instance, neutrophil traits were enriched not only in the neutrophil facet but also in monocyte^76^ and monocyte-related leukemia facets, while eosinophil traits showed enrichment across both myeloid (e.g., mast cell, monocyte) and lymphoid^77^ (e.g., T cell, natural killer cell, lymphoblastoid) cell facets, consistent with the broader immunoregulatory roles of eosinophils across immune contexts^78,79^. Furthermore, monocyte-associated cell facets, including monocytes and AML subtypes M5 and M4eo, showed enrichment for variants associated with various myeloid-associated traits, consistent with the functional plasticity of monocytes in homeostasis and immunity^80–82^.

### Utilizing the PRIME-enhanced CRE atlas for variant interpretation

We next examined trait-associated variants listed by the Open Targets platform^83^. When combined with enhancer-gene regulatory predictions from scE2G^84^ and ENCODE-rE2G^43^, the PRIME atlas revealed several links between trait-associated variants, cell-type-specific enhancers, and candidate effector genes.

First, an intronic distal CRE (chr15:67121091-67121491) within the *SMAD3* locus overlaps a variant (rs17294280, 15_67121286_G_A) strongly associated with asthma^85^ (P = 5.77×10^-23^; PIP = 0.995, PICS), lung function (FEV/FVC ratio^86^; P = 1.18×10^-12^; PIP = 0.194, SuSiE-inf), and allergic rhinitis^87^ (P = 6.0×10^-12^; PIP = 0.911, PICS) (**Fig. 5D**, **Supplementary Note 3**, **Fig. SN3**). This putative enhancer shows highest PRIME activity in immune cell types, particularly T cells, monocytes, and mast cells. The variant is further supported by eQTL evidence linking it to *SMAD3* expression in blood (P=0.000011, effect size = −0.23, GTEx^53^ phs000424.v10.p2) and by enhancer-gene predictions from scE2G and ENCODE-rE2G, nominating *SMAD3* as the likely effector gene. Given the central role of SMAD3 in TGF-β signaling^88^ and immune regulation, and prior reports linking *SMAD3* methylation at birth to asthma^89^, this locus provides a plausible regulatory mechanism linking immune-cell enhancer activity to asthma susceptibility.

Second, PRIME identified a monocyte-active distal CRE (chr12:124626205-124626605) located ∼58 kb from the transcription start site of *NCOR2* that overlaps a variant (rs763017549, 12_124626281_C_T) associated with premature separation of placenta^90^ (abruptio placentae) (P = 9.80×10^-18^; PIP = 1.0, PICS) (**Fig. 5E**, **Supplementary Note 3**, **Fig. SN4**). The CRE shows strong activity in monocytes and is linked to *NCOR2* by scE2G predictions, suggesting a regulatory connection between immune-cell transcriptional programs and placental biology. *NCOR2* encodes a transcriptional co-repressor that recruits histone deacetylase complexes^91^ and has previously been implicated in trophoblast differentiation^92^ and preeclampsia^93^, providing a potential mechanistic explanation for this association.

The PRIME-enhanced CRE atlas also revealed less well-characterized biological connections. One example is the unexpected enrichment of variants related to non-iron deficiency anemia (e.g., “Other anemias”) within mast-cell PRIME loci (**Fig. 5B**). For example, a mast cell-active distal CRE (chr14:24517772-24518172) at the *CMA1* locus harbors a variant (rs76735545, 14_24518091_C_G) associated with pernicious anemia^94^ (P = 1.31 × 10^-11^; PIP = 1.0, SuSiE-inf) (**Supplementary Note 3**, **Fig. SN5**). The regulatory potential of this variant is supported by ENCODE_rE2G^43^ linking the putative enhancer to *CMA1* in HepG2 cells, which also expresses the gene. The variant (14-24518091-C-T) is also an eQTL for *SDR39U1* (skeletal muscle, P = 5.7 × 10^-9^, effect size = −0.370, GTEx^53^ phs000424.v10.p2). This suggests a mast cell-driven role in erythroid physiology and anemia pathogenesis.

Beyond these cases, additional examples include loci linking immune cell type distal CREs to anthropometric traits (*ACOT11*/*SSBP3*), platelet traits (*CD82*), and eosinophil counts (*IKZF3*) (**Supplementary Note 3**). Together, these loci demonstrate the utility of the PRIME atlas in providing cell-type-specific gene regulatory context for interpreting GWAS variants.

## Discussion

Identifying active cis-regulatory elements (CREs) and the cell types in which they are active remains a core challenge in genomics. Chromatin-based maps catalog candidate regulatory regions but may lack specificity for regulatory activity^2,10^. TSS-based assays provide a direct readout of regulatory activity but have been constrained by limited sensitivity, input requirements^13,54,55^, and heuristic enhancer definitions^2^. Here, we address these limitations by developing nucCAGE, a nuclear TSS assay enriching for unstable transcripts, and PRIME, a predictive model for identifying active CREs from initiation profiles, enabling sensitive and cell-type-resolved mapping of CRE activity.

nucCAGE offers a practical alternative to dedicated nascent transcription assays. Methods such as GRO-cap^34^ and PRO-seq^95^ provide detailed mechanistic insight but require large input amounts^40^ (50–100 million cells) and protocols that limit their use largely to cell lines^96^, while whole-cell CAGE accurately captures promoter usage but underrepresents rapidly degraded transcripts such as enhancer RNAs and promoter upstream transcripts^25^. In contrast, nucCAGE enriches unstable and nuclear transcripts through nuclear extraction while retaining modest input requirements compared to nascent transcription assays. By sampling a mixture of nascent and unstable, short-lived nuclear transcripts, nucCAGE increases sensitivity to initiation at promoters and enhancers while retaining a clear relationship to steady-state gene expression. Across cell types, nucCAGE preserves promoter-associated signal while markedly increasing sensitivity to intronic and intergenic initiation events, extending TSS-based profiling to primary cells and disease-relevant tissues and enabling analysis of low-abundance initiation events.

Interpreting increased TSS sensitivity requires computational approaches that generalize across library complexity and input. Existing methods for CRE detection from TSS data rely on fixed thresholds or heuristic features that are sensitive to sequencing depth and sample-specific properties^17^. PRIME overcomes these limitations by learning characteristic initiation patterns directly from TSS signal profiles, emphasizing signal shape rather than magnitude. This enables consistent CRE prediction across nucCAGE and whole-cell CAGE datasets. Across multiple benchmarks, including ClinVar^52^ variants, fine-mapped eQTLs^53^, and CRISPRi perturbation datasets^43,68^, PRIME achieves a superior balance of recall and precision relative to chromatin-based and nascent transcription-based approaches.

Notably, the high recall achieved by PRIME in nucCAGE datasets, at the expense of a slight decrease in precision, likely reflects the capture of a continuum of enhancer activities. This observation aligns with recent high-throughput perturbation results using DC-TAP-seq, which demonstrate that many functional enhancers exhibit small effect sizes^68^. By identifying these low-effect-size enhancers that more stringent methods may miss, PRIME provides a more comprehensive landscape of the regulatory genome, suggesting that a considerable fraction of predicted negatives in previous perturbation screens may indeed harbor functional, albeit subtle, regulatory potential.

At the atlas scale, PRIME substantially improves the resolution at which regulatory activity can be linked to cell types and traits. Applied to FANTOM5^44^ data, PRIME identifies a larger and more cell-type-specific set of distal CREs than the original FANTOM5 enhancer atlas^17^ while preserving strong enrichment for trait-associated genetic variation. Stratified LD score regression^24^ across thousands of traits recapitulates known relationships between traits and cell types and highlights additional, biologically plausible associations, including a putative link between mast cells and non-iron deficiency anemia. The adjustable PRIME score thresholds enable tuning between recall and precision, supporting the use of this resource for discovery-oriented analyses and targeted variant-to-function interpretation in a cell-type-specific manner.

Despite these advances, several limitations remain. nucCAGE does not capture a purely nascent RNA population but samples a mixture of nascent and unstable, short-lived nuclear transcripts, reflecting a deliberate trade-off between sensitivity and specificity and the aim to overcome constraints in input material^42^. Library complexity further constrains performance. While PRIME increases sensitivity to CRE detection, the number of detectable loci remains dependent on library complexity, making the increased transcriptional sensitivity provided by nucCAGE particularly important for maximizing the number of CREs identified. Increasing nuclear input improves sensitivity up to an optimal range (2.5–5 million nuclei), whereas excessive input leads to noise accumulation, possibly driven by the capture of decaying RNA fragments. The enhanced sensitivity to transcription initiation provided by nucCAGE enables more quantitative interrogation of regulatory element activity. Such data may support future functional classification of CREs based on initiation features such as frequency, strand balance^97^, and multi-peak modality^44,98^, rather than binary activity. Together, nucCAGE and PRIME provide tools that balance sensitivity, interpretability, and applicability across diverse cell types and input constraints.

## Acknowledgements

R.A. acknowledges support from the Novo Nordisk Foundation (NNF20OC0059796, NNF24OC0096127), the Novo Nordisk Foundation Center for Genomic Mechanisms of Disease (NNF21SA0072102), and the Independent Research Fund Denmark (6108-00038). R.K. acknowledges support from the Lundbeck Foundation (R370-2021-924), Villum Foundation (VIL72111) and the Independent Research Fund Denmark (5281-00295B).

We thank members of the Andersson lab, Tissue Gene Regulation lab, and the Novo Nordisk Foundation Center for Genomic Mechanisms of Disease for feedback on the methods and results.

## Author contributions statement

H.E. and N.N. co-led the project as co-first authors; H.E. and R.K. analyzed and interpreted CAGE data with respect to RNA localization, sample complexity, and nascent transcription; R.K. and L.J. developed the nuclei extraction protocol; C.S.V. and R.K. performed CAGE library preparation and sequencing; S.P., M.S., and R.A. developed initial methods for predicting regulatory elements and associated model training strategies; N.N. developed and trained the PRIME models and computed predictions across cell lines and FANTOM data; H.E., N.N., N.A., M.T., M.G.E., B.L.G., A.S., R.K., and R.A. developed the PRIME R package; H.E., N.A., M.T., M.G.E., A.S., R.K., and R.A. developed the PRIMEprep CAGE data processing pipeline; H.E., N.N., R.K., and R.A. analyzed and interpreted data in the development of PRIME and PRIMEmodel; H.E., W.Q., M.U.S., J.M.E., and R.A. developed the CRISPR, eQTL, and GWAS benchmarking pipelines; H.E., N.N., W.Q., R.K., and R.A. performed PRIME model benchmarking; H.E., N.N., T.K., H.T., P.C., R.K., and R.A. developed the strategy to analyze FANTOM CAGE data to build the PRIME atlas; H.E, N.N., D.A.S., B.E.B., R.K., and R.A. performed PRIME-guided regulatory architecture dissection; H.E., J.M.E., and R.A. developed the analysis strategy for FANTOM atlas-guided GWAS interpretation; H.E., R.K., and R.A. analyzed cell line and FANTOM data to interpret GWAS variants; R.K. and R.A. provided overall supervision for the project; H.E., N.N., R.K., and R.A. wrote the manuscript with input from all authors.

## Methods

### Cell culture

K562 [ATCC-CCL-243], HCT116 [ATCC-CCL-247], A549 [ATCC CCL-185], and HepG2 [ATCC-HB-8065] cells were acquired from the American Type Culture Collection via LGC Standards. GM12878 [CEPH/UTAH Pedigree 1463] cells were purchased from the NIGMS Human Genetic Cell Repository at the Coriell Institute. K562 cells were cultured in Advanced RPMI 1640 medium [Thermo Fisher Scientific (TFS), 12663012], HCT116 in DMEM with sodium pyruvate and low glucose [TFS, 11885084], A549 in Ham’s F-12 Nutrient Mix [TFS, 11765054], HepG2 in DMEM [TFS, 41965039] and GM12878 in RPMI 1640 without added L-Glutamine [TFS, 21870076]. Medium supplements comprised 10% (v/v) FBS [TFS, 10500056/10500064] and 1% (v/v) Penicillin-Streptomycin [TFS, 15140122] for all 5 cell lines, 1% (v/v) L-Glutamine [TFS, 25030081] for K562, HepG2, and GM12878 cells, as well as 1 g/l Glucose [TFS, A2494001] for A549 cells.

The confluent cell lines A549, HepG2, and HCT116 were cultured until 75-80% confluence before trypsinization with 0.25% trypsin-EDTA [TFS, 25200056] and splitting 1:5-1:8. The two suspension cell lines, K562 and GM12878, were cultured until a concentration of 7.5 × 10^6^ cells/ml was reached before splitting them 1:3.

All adherent and suspension cell lines were reared in Corning T-25 or T-75 flasks at 37 °C with 5% CO_2_.

### Triptolide treatment

5 ml GM12878 cells were seeded in 6 well plates [Greiner Bio-one, 657160] at a concentration of 2 × 10^5^ cells/ml and incubated for 16 h prior to triptolide treatment. Either 50 µl of a 10 mM triptolide [Sigma-Aldrich, T3652] solution or 50 µl of DMSO as control were added, and cells were incubated for 5 h prior to harvesting and lysis. Triplicates were prepared for both conditions.

### Nuclei extraction and CAGE-sample preparation

A549, HepG2, and HCT116 cells were washed in their culture flasks with DPBS without divalent cations [TFS, 14190136], trypsinized with 0.25% trypsin-EDTA [TFS, 25200056], harvested, and washed once again with the same DPBS solution. The K562 and GM12878 suspension cells were harvested by centrifuging cells at 1,000 rpm at room temperature for 2 min in an Eppendorf 5810R centrifuge and washed once with PBS [TFS, 10010023]. Cells of all lines were counted with the Countess II FL Automated Cell Counter [TFS, AMQAF1000] and disposable cell-counting chamber slides [TFS, C10228].

Three replicates each of 2.5⋅10^6^ K562, HCT116, A549 and HepG2 cells as well as 5⋅10^5^ and 10^6^ GM12878 cells were aliquoted, pelleted, and immediately dissolved in 300 µl of RNA lysis buffer from the PureLink RNA Mini Kit [TFS, 12183025] including 1% β-Mercaptoethanol [Sigma-Aldrich, M6250]. These samples constituted the whole cell-samples.

3 replicates each of 2.5⋅10^6^ and 10^7^ K562 cells; 5⋅10^5^, 10^6^, 5⋅10^6^, and 10^7^ GM12878 cells, 2.5⋅10^6^ A549 and 2.5⋅10^6^ HepG2 cells were, in preparation for the nuclei extraction, aliquoted and pelleted. Nuclei were extracted by dissolving these cell pellets in 500 µl hypotonic lysis buffer (10 mM Tris-HCl, pH 7.4; 10 mM NaCl; 3 mM MgCl_2_; 1% (v/v) BSA; 0.1% (w/v) NP-40 Substitute [Roche, 11754599001]; 0.1% (v/v) Tween20 [Roche, 11332465001]; 0,01% (v/v) Digitonin [Promega, G9441], 4 mU/µl SUPERase·In [TFS, AM2696]; 4 mU/µl Protector RNase Inhibitor [Sigma-Aldrich, 3335402001]). Samples were incubated for 5 min on ice (4°C) and mechanically sheared by passing them 10 times through a 1 ml syringe assembled with a 27G needle. The resulting nuclei suspension was diluted with 9.5 ml hypotonic resuspension buffer (10 mM Tris-HCl, pH 7.4; 10 mM NaCl; 3 mM MgCl_2_; 1% (v/v) BSA; 0.1% (w/v) Tween20 [Sigma-Aldrich, 655206]; 4 mU/µl SUPERase·In [TFS, AM2696]; 4 mU/µl Protector RNase Inhibitor [Sigma-Aldrich, 3335402001]) to dilute the detergents, and centrifuged for 10 min at 4°C and 500×g. The resulting nuclei pellet was dissolved in 300 µl of RNA lysis buffer from the PureLink RNA Mini Kit. These samples constitute the nuclei samples.

RNA extraction for all replicates of the nuclei and whole-cell samples was performed with the PureLink RNA Mini Kit according to the manufacturer’s protocol including on-column DNase digestion using the PureLink DNase Set [TFS, 12185010]. All samples were quality checked by automated gel electrophoresis of 2.5 ng per replicate using RNA 6000 Pico kits [Agilent, 5067-1513] on an Agilent 2100 Bioanalyzer.

### Staining and Flow cytometry

K562 nuclei were isolated from 2⋅10^6^ input cells using either the nuclei extraction as part of the Omni-ATACseq protocol^99^ (Omni) or the optimized protocol (N27) incorporating ten passes through a 27G needle as outlined above. Following extraction, nuclei were gently resuspended in 285 μl ice-cold flow cytometry buffer (FCB, 1% (w/v) BSA in DPBS).

Nuclei were incubated with rabbit monoclonal anti-TOMM20 antibody (ST04-72 clone, [TFS, MA5-32148]) at a final concentration of 5 μg/ml for 45 min on ice with gentle agitation. Following washing in FCB and centrifugation (500×g, 10 min, 4 °C), samples were incubated with Alexa Fluor 488-conjugated goat anti-rabbit IgG (H+L) secondary antibody [TFS, A-11034] at a final concentration of 1 μg/ml for 15 min on ice with gentle agitation and protected from light. Samples were washed again in FCB and subsequently stained with DAPI at a final concentration of 10 μg/ml for 5 min on ice with gentle agitation. After a final wash in FCB, samples were resuspended in 300 μl ice-cold FCB.

Control samples were acquired for both extraction protocols and included 1) whole K562 cells, 2) unstained nuclei, 3) nuclei only stained with DAPI and 4) nuclei only stained with anti-rabbit IgG 2nd antibody. Whole K562 cells were subjected to the same staining procedure but centrifugation was adapted to 1000 rpm for 2 min at RT instead.

Flow cytometric analysis was performed on a BD FACSMelody™ Cell Sorter (BD Biosciences). Alexa Fluor 488 and DAPI fluorescence were acquired using the FITC and BV421 detection channels, respectively. Detector voltages were established using unstained, single-stained and double-stained controls. A total of 10,000 events were recorded per sample. Data were analyzed in RStudio and visualized using ggcyto (v1.39.4) and ggplot2 (v4.0.3). Nuclei were identified by FSC-A/FSC-H singlet gating and DAPI positivity.

### CAGE library preparation

CAGE libraries were generated following the SLIC-CAGE^56^ and nAnT-iCAGE^55^ protocols. In brief, total RNA from individual nuclear or whole-cell replicates (up to 4,000 ng) was combined with prepared SLIC-CAGE carrier RNA to a total input of 5,000 ng RNA. These carrier-combined samples served as input material for first-strand cDNA synthesis using the SuperScript IV reverse transcriptase [TFS, 18090050] and random hexamer primers [IDT, 5’-N_6_-TCT-3’]. Next, the diol of 7-methylguanosine (m7G) in the RNA of the generated RNA:cDNA hybrids was oxidised via sodium periodate [NaIO_4_, Sigma-Aldrich, 311448]. As m7G are unique to capped RNA species, this oxidation enables selective biotinylation of capped RNAs at the 5’ end with Biotin (Long-Arm) Hydrazide [VectorLaboratories, SP-1100] in the next step.

3’ single stranded-RNA stretches not reverse transcribed and incorporated into the RNA:cDNA hybrids, but biotinylated at their 3’ nucleotide were removed through RNase ONE [Promega, M4261] digestion to ensure the subsequent pull-down using paramagnetic M-270 Streptavidin beads [TFS, 65306] selectively captures the biotinylated m7Gs (i.e., cap-trapping). Subsequent RNaseH [Takara, 2150A] and RNase ONE [Promega, M4261] digestion ensured the release of cDNAs representing exclusively capped RNAs from the streptavidin beads and RNA:cDNA-hybrids.

This was followed by ligation using MightyMix [Takara, 6023] of the Illumina-compatible 5’ and 3’ nAnT-iCAGE linkers and second-strand cDNA synthesis with the nAnT-iCAGE second primer and Deep Vent (exo-) DNA polymerase [NEB, M0259S]. Carriers underwent digestion with I-Scel [NEB, R0694S] and I-CeuI [NEB, R0699S], after which the resulting library was size-selected and PCR amplified.

The average library length was determined from the fragment distribution measured on the Agilent 2100 Bioanalyzer using the High Sensitivity DNA kit [Agilent, 2067-4626], and libraries were quantified using the Quant-iT PicoGreen dsDNA kit [TFS, P7589]. Equimolar pooling of sets of eight samples was performed prior to single-end sequencing on a NextSeq 550 Illumina sequencer with High Output v2.5 (75 cycles) reagents [Illumina, 20024906] utilizing standard Illumina primers. Libraries were spiked with 3% PhiX from the NextSeq PhiX Control Kit [Illumina, FC-110-3002].

### CAGE data preprocessing and mapping

For processing of CAGE sequencing data, we developed the PRIMEprep preprocessing and mapping pipeline (https://github.com/anderssonlab/PRIMEprep, v0.1.0), which integrates and standardizes all steps described below to ensure high-quality data for downstream analysis.

Initial quality control of the raw sequencing reads was performed using FastQC (v0.12.1). Raw FASTQ files were trimmed using fastp^100^ (v0.23.4) to remove the first 3 bases from each read to eliminate barcode sequences (i.e., --trim_front1 3). Reads with fewer than 50% of bases having a Phred quality score ≥20 were discarded (--length_required 30 --qualified_quality_phred 20 --unqualified_percent_limit 50). FastQC was run both before and after trimming and filtering. Ribosomal RNA reads were removed using rRNAdust^101^ (v1.02) (an optional step in PRIMEprep) using the U13369.1 rRNA reference prior to genome mapping.

Filtered reads were mapped to the GRCh38 ENCODE GRCh38_no_alt_analysis_set_GCA_000001405.15 reference genome using STAR^102^ (v2.7.3a) in single-end mode with parameters optimized for CAGE data, including extension of alignments at the 5′ end of reads (--alignEndsType Extend5pOfRead1). Only uniquely mapped reads were retained (--outFilterMultimapNmax 1). STAR was run to produce coordinate-sorted BAM files.

Post-mapping QC involved assessment of mapping statistics using samtools^103^ (v1.21, using htslib 1.21) and library complexity using preseq^104^ (v2.0). Unmatched non-templated guanine additions at the 5′ ends of reads were corrected in a strand-specific manner by identifying 5′ terminal G residues not supported by the reference genome based on read sequence and alignment mismatch information encoded in the MD tag. When detected, the affected reads were adjusted by removing the non-templated G and shifting the corresponding 5′ end coordinate by one or two bases, depending on the number of non-templated Gs detected.

CAGE-inferred TSSs (CTSSs) were defined as the genomic positions corresponding to the 5′ ends of uniquely mapped reads after G correction. CTSSs were collapsed by genomic position and strand to generate CTSS BED files. Strand-specific bedGraph files were generated from CTSS counts and converted to bigWig format using bedGraphToBigWig for downstream analysis using PRIME.

### PRIME: an R package for data-driven regulatory element analysis from TSS data

Downstream CAGE analysis was performed in R (v4.2.2) using PRIME (https://github.com/anderssonlab/PRIME, v0.1.0) and CAGEfightR^57^ (v.1.18.0). We developed PRIME to extend CAGEfightR with additional utilities for quantitative analyses of transcription initiation data, with a focus on regulatory element characterization. PRIME operates on CTSS-level and CTSS cluster (tag cluster)-level data represented as SummarizedExperiment and GRanges objects and is fully compatible with Bioconductor genomic infrastructure. PRIME provides utilities for quantifying transcription initiation complexity, signal saturation, and background noise, enabling robust quantitative comparison between libraries. The package further supports identification and characterization of divergently transcribed loci and core promoter architectures, including core promoter decomposition and analyses of strand balance, positional dispersion, and sequence-associated initiator patterns. In addition, PRIME implements functions for subsampling, normalization, and profile-based summarization of CTSS signal around genomic features, facilitating systematic regulatory element analyses from CAGE data or other TSS assay data. PRIME also includes an interface to the PRIME LightGBM (LGBM) model (https://github.com/anderssonlab/PRIMEmodel, v1.0.0), developed in this study, for model-based regulatory element scoring based on CTSS data.

### TSS level analysis

Strand-specific CTSS bigWig data generated by PRIMEprep were imported into R as SummarizedExperiment objects using CAGEfightR^57^ and filtered to remove singletons supported by a single read. Replicate pools were constructed by aggregating CTSS counts across samples. CTSS signal was analyzed either at single-nucleotide resolution, by aggregation into strand-specific positional clusters of CTSSs (tag clusters; see section ‘*Core promoter level analysis*’), or by aggregating counts within 401 bp windows centered on DHS^9^ summits (±200 bp).

Library complexity was assessed after library size-matched subsampling, in which CTSS libraries were subsampled without replacement to the minimum total CTSS count among the samples included in each comparison, ensuring that differences in sequencing depth do not confound complexity estimation. Cumulative CTSS signal distributions across DHSs, computed by ranking DHSs by total CTSS signal and calculating cumulative fractions as a function of rank, were used to compare complexity and signal saturation between samples. In this context, library complexity reflects the number of DHSs required to account for a given fraction of CTSS signal, while signal saturation reflects the degree to which total CTSS signal is concentrated among high-signal DHSs. Analogous complexity analysis was performed for CTSS genomic locations, quantifying the signal distribution among CTSSs.

Genomic annotation of CTSSs was performed hierarchically relative to UCSC knownGene transcript models (TxDb.Hsapiens.UCSC.hg38.knownGene, v.3.16.0), assigning CTSSs to (by decreasing priority) promoter, promoter-proximal, 5′ untranslated region (UTR), 3′ UTR, coding sequence (CDS), exon, intron, antisense, or intergenic categories. Fractions of CTSS signal mapping to intergenic, intronic, and exonic regions were computed, and relative changes in intergenic-to-intronic and intronic-to-exonic CTSS signal fractions were compared across samples and conditions.

For visualization, CTSS signals were converted to tags per million (TPM) and used exclusively for visualization. Strand-specific CTSS signals were scaled such that plus-strand CTSS values were plotted as positive values and minus-strand CTSS values as negative values. Normalized strand-specific CTSS signals were visualized using Gviz (v.1.42.1) for track-level plotting.

### Nascent transcription-associated analysis

For validation against independent nascent transcription assays, DHS-associated CTSS signal from pooled GM12878 replicates was compared with publicly available GRO-cap data (ENCODE; ENCFF638SZH, ENCFF740NAF). In addition, genomic annotation distributions of CTSSs were compared with NET-CAGE^42^ (GSE118075) GM12878 data.

For each DHS, total CTSS signal was quantified for nucCAGE and whole-cell CAGE libraries, including triptolide-treated and control samples, and TPM normalized. Differences in the proportions of DHSs showing increased versus decreased CAGE expression between conditions were evaluated using chi-squared tests, with statistical significance assessed by permutation testing (10,000 iterations) to derive empirical null distributions.

Specifically, we calculated the log_2_ fold-change of expression between nuclei and whole-cell CAGE, as well as the log_2_ fold-change between triptolide-treated and control whole-cell samples. To evaluate the robustness of the association between nucCAGE enrichment and transcriptional arrest, we performed 10,000 permutations. In each iteration, the relationship between nucCAGE/whole-cell ratios and the inhibitor response (triptolide log_2_ fold-change) or nascent signal (GRO-cap log_2_ fold-change) was randomized to generate an empirical distribution of Chi-square statistics. Empirical p-values were then determined by comparing the observed Chi-square statistic against this null distribution.

Additionally, we employed Wilcoxon rank sum tests to compare the distribution of triptolide sensitivity and GRO-cap signal between nuclear-enriched and whole-cell-enriched transcripts. DHSs were stratified into gene-proximal (±500 bp) and gene-distal (> 500 bp) classes based on proximity to annotated UCSC knownGene TSSs to assess nascent enrichment across different regulatory contexts.

### Core promoter level analysis

CTSSs on each strand were grouped into strand-specific tag clusters, in which adjacent CTSSs were merged if separated by no more than 20 bp, defining CAGE tag clusters, herein referred to as core promoters. Core promoter expression was quantified as the sum of CTSS counts within each tag cluster, followed by TPM normalization.

Core promoters were assigned to genes based on genomic proximity (within 1000 bp upstream and 100 bp downstream) to UCSC knownGene TSSs. For gene-level analyses, core promoter-associated CTSS signal was summed across all core promoters assigned to the same gene, yielding gene-level expression estimates. GM12878 CAGE-derived gene expression estimates were compared with GM12878 bulk RNA-seq (ENCODE^105^; ENCFF345SHY) and GRO-cap (ENCODE; ENCFF638SZH, ENCFF740NAF) datasets, using Entrez gene identifiers derived from Ensembl annotation to enable for cross-modality gene mapping.

Initiator sequence composition was analyzed at dominant CTSS positions (summits) within each core promoter. For each promoter, the CTSS with the highest expression was identified, and the underlying genomic sequence was extracted. Initiators were classified according to nucleotide composition at the −1/+1 positions relative to the CTSS, distinguishing canonical YR (CA, TA, CG, TG) and YC (CC, TC) initiator classes, as well as other sequence contexts. The impact of 5′ G-correction was evaluated using paired core promoter-level analyses on identical libraries processed with and without correction, with a focus on changes in initiator sequence composition at dominant CTSS positions.

Core promoter architecture was characterized using CTSS positional complexity metrics calculated using pooled replicate CTSS signal. For each core promoter, pooled strand-specific CTSS signals were first normalized to relative frequencies across all CTSS positions within the core promoter. Core promoter breadth was quantified as the interquantile width (10-90%) of the cumulative CTSS signal distribution, corresponding to the genomic span containing the central 80% of initiation events^44,57^. Initiation dispersion was quantified using shape entropy^44,57^, computed as Shannon entropy over the normalized CTSS position frequencies. Shape entropy values were normalized by the maximum possible entropy given the number of CTSS positions with non-zero signal per promoter, yielding a value between 0 and 1. This normalization ensures that entropy values are directly comparable across core promoters with differing numbers of initiation sites.

### Identification and characterization of divergently transcribed loci

Divergently transcribed loci were identified from tag clusters derived from pooled replicate CTSS signal, as previously defined^17^. Briefly, candidate divergent loci were defined by first identifying pairs of opposing-strand, divergently oriented core promoters separated by ≤400 bp. These pairs were merged into divergent loci based on shared core promoter usage, with prioritization during merging determined by tag cluster expression level. For each merged divergent locus, the resulting plus-strand and minus-strand regions were used to define a single locus midpoint, which was then used to summarize strand-specific CTSS signal within 200 bp windows upstream and downstream of this midpoint for minus strand and plus strand expression, respectively. Transcriptional directionality was quantified as the normalized imbalance between plus- and minus-strand expression. Directionality was defined as (P2 - M1) / (P2 + M1), where P2 and M1 represent the aggregated expression of the 200 bp windows immediately flanking the midpoint downstream (plus strand) and upstream (minus strand), respectively. This directionality index ranges from -1 to 1, where values near 0 represent balanced bidirectional initiation and values approaching the extremes indicate a strong strand bias. Loci with strong strand bias were excluded using an absolute value directionality cutoff of ≤ 0.8, retaining bidirectionally transcribed regions. These divergent loci were used for benchmarking purposes regardless of their divergent signature, as defined below. For downstream analyses, both strand CTSS signals were aggregated, yielding divergent locus-level expression estimates.

Each divergent locus was classified as having a divergent signature if (M1 > P1) & (P2 > M2), where P1 and M2 represent the expression of the corresponding opposite strands in windows defined for M1 and P2, respectively. Loci were further annotated by genomic context (proximal vs. distal) based on TxDb.Hsapiens.UCSC.hg38.knownGene. We evaluated how the directionality score varied across different genomic regions and samples. To validate the accessibility of divergent loci, stratified by directionality score and divergent signature and expression level, the loci were overlapped with single-nucleus ATAC-seq (snATAC-seq) data^106^. We further compared these properties between loci with detectable chromatin accessibility (DHS-positive) and those without (DHS-negative).

### Noise and background analysis

Sample-specific technical noise in CTSS signal was estimated empirically from genomic regions unlikely to represent transcription initiation. A total of 1,000,000 regions were sampled from 1,781,993 genomic background windows, each 200 bp in width. Background windows were required to be intergenic or intronic with respect to UCSC knownGene annotations, to not overlap ENCODE blacklist regions (ENCFF356LFX), and to be be positioned ±300 bp away from ENCODE hotspot regions derived from a previously compiled diverse set of 706 human DNase-seq samples^98^. Windows were further limited to autosomal uniquely mappable regions, defined based on a CAGE read length of 73 bp.

For each sample, CTSS signal was aggregated within background windows, and the resulting background signal distribution was used to empirically characterize condition- and input level-specific noise levels.

We further computed CTSS signal fingerprints to assess the distribution of CTSS signal concentration across ranked CTSSs. For each replicate, CTSS counts were extracted and sorted by expression, and cumulative CTSS signal was calculated as a function of CTSS rank. Cumulative signal was normalized by the total CTSS signal per sample to obtain relative cumulative fractions. To enable efficient comparison across samples, cumulative profiles were downsampled to 100,000 uniformly spaced rank positions, retaining the final cumulative value.

### Dataset construction for PRIME model training

A schematic overview of the dataset construction and model training is shown in **Figure S7**. Positive and negative training regions were defined using CAGE and chromatin accessibility data. GM12878 500K and 1M whole-cell CAGE and 1M, 5M, and 10M nucCAGE were included. DHSs for GM12878 were obtained from the NIH Roadmap Epigenomics Consortium^9^ (sample E116, narrowPeak format) and lifted from GRCh37 to GRCh38 using LiftOver^107^. DHSs overlapping ENCODE Blacklist^108^ regions were excluded.

Initial positive regions were defined based on DHSs. DHS summits were extended by ±200 bp, and only regions containing CAGE signals within these windows were retained, resulting in 401 bp regions centred on DHS summits. Initial negative regions were defined from tag cluster summits, extended by ±200 bp, and restricted to regions not overlapping the initial positive set, thereby selecting CAGE-associated regions lacking DHS support. A filtering step was applied to exclude regions supported by fewer than 4 CAGE reads. Positive regions overlapping GENCODE (v31) annotated exons, CDS, and 3′ UTRs were further removed.

To increase the size of the training dataset, filtered positive regions were augmented by applying normally distributed random shifts (±30 bp) around the region midpoint, followed by removal of redundant regions. The filtered positive regions and the augmented regions were pooled to form the combined positive set.

The combined negative set consisted of augmented negative regions and near-positive negative regions. Initial negative regions were augmented using uniformly distributed random shifts to off-center regions relative to tag cluster summits. Near-positive negative regions were defined as sequences flanking the 151 bp core regions of filtered positive regions, with less than 50% overlap with any region in the positive set. The combined negative set was randomly subsampled to match the size of the combined positive set while preserving the original distribution of annotation types.

The final PRIME model was trained on a dataset comprising 4,428,109 regions.

### PRIME model training

CAGE signal profiles within all regions in the positive and negative sets were locally normalized prior to model training. For each region, strand-specific CTSS signals were represented as forward (*f*) and reverse (*r*) vectors and transformed into a normalized strand-difference CAGE profile, defined as:

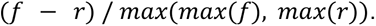

This transformation yields a 401 position profile vector per region, with values ranging between –1 to 1, preserving local strand asymmetry while removing dependence on global signal amplitude. Processed CAGE profiles were used as input for model training. For model evaluation, chromosomes 2-4 were held out as a test set. The remaining chromosomes were randomly split into 80% training and 20% validation sets, while preserving the natural distribution of annotation classes.

The PRIME model is a binary classification light gradient boosting machine (LGBM) implemented in Python (v3.9.9) using LightGBM^109^ and scikit-learn^110^. Hyperparameter tuning was performed using Bayesian optimization implemented with Weights & Biases (wandb) (https://www.wandb.com/), resulting in the final model configuration: *n_estimators* = 1686, *num_leaves* = 1463, *learning_rate* = 0.09377, *max_depth* = 30, *min_split_gain* = 0, *reg_alpha* = 0, *reg_lambda* = 50, *boosting_type* = ‘gbdt’. The model was trained using binary cross-entropy loss. Performance was evaluated on validation and independent test sets using accuracy, precision, recall, and F1-score. All runs used fixed random seeds, and no class reweighting was required due to the balanced training dataset.

### Using PRIME model for genome-wide predictions

A genome-wide prediction pipeline was developed to apply the trained PRIME model to TSS data, including CAGE, for identification of likely CREs (including both enhancers and promoters).

Genome-wide CTSSs were first identified from CAGE bigWig files using CAGEfightR, followed by identification of strand-specific tag clusters. Candidate regions were defined from tag cluster summits extended by ±200 bp to define 401 bp regions, which were reduced to obtain a set of non-overlapping regions. Within these regions, 401 bp sliding windows with a step size of 20 bp were generated to tile the entirety of the reduced non-overlapping regions. For each sliding window, strand-specific CTSS signals were transformed into normalized strand-difference CAGE profiles as described above. These profiles were provided as input to the PRIME model to obtain a prediction score for each window.

Predicted windows and their scores were subsequently post-processed using a segmentation strategy to identify high-confidence, non-overlapping regions, and to decompose broad regions of transcription initiation into subregions. In this procedure, sliding windows were first filtered using a global score threshold (default = 0.75). Overlapping windows passing this threshold were clustered. Within each cluster, only the 151 bp core region of each window was considered. The core with the maximum prediction score within a cluster (*max_score*) was identified, and all other cores overlapping this core were evaluated using a local stringency threshold defined as *max_score – d* (default *d* = 0.1). Cores passing this threshold were retained and merged to define a region of interest. Cores already processed, as well as cores overlapping the resulting regions, were excluded from further consideration. This procedure was iteratively repeated within each cluster until all cores had been evaluated. The final output consisted of non-overlapping regions of at least 151 bp, each assigned a representative score corresponding to the maximum prediction score, and a summit point at the midpoint of the highest-scoring core.

All steps of the genome-wide prediction pipeline were implemented using R and Python, and the complete R package and associated bash pipeline are available in the PRIMEmodel GitHub repository (https://github.com/anderssonlab/PRIMEmodel, v1.0.0).

We would like to note that PRIME assigns prediction scores to genomic windows based on initiation patterns, and does not inherently distinguish between promoters and enhancers. The final classification into proximal or distal categories was performed post-hoc using the TxDb.Hsapiens.UCSC.hg38.knownGene annotation.

Performance of genome-wide PRIME predictions was evaluated on the held-out test set comprising chromosomes 2-4 by comparison with predictions obtained using divergent loci calling, a FANTOM5-style method^17^ described above. Predicted sites from both methods were intersected with DHS regions defined as 401 bp windows centered on DHS summits, and only predictions whose summits fell within these DHS windows were retained for evaluation. Precision was defined as the fraction of prediction sites overlapping DHS windows with detectable CAGE signal, and recall was defined as the fraction of DHS windows with CAGE signal that were recovered by predictions.

### External regulatory element prediction datasets

External regulatory element predictions for K562 cells were collected from multiple published sources. PINTS peaks^35^ from GRO-cap and PRO-cap datasets were downloaded from the PINTS web portal (https://pints.yulab.org/). ENCODE candidate cis-regulatory elements (cCREs)^8^ for K562 were obtained from the SCREEN web portal (https://screen.wenglab.org/), retaining promoter-like signatures (PLS), proximal enhancer-like signatures (pELS), and distal enhancer-like signatures (dELS). dREG peaks derived from K562 PRO-seq and qPRO-seq data (GSE150625) were also included^96^. K562 H3K27ac ChIP-seq peaks were downloaded from ENCODE (ENCFF038DDS), and DHSs were obtained from the NIH Roadmap Epigenomics Consortium^9^ and lifted from GRCh37 to GRCh38.

All regulatory element datasets were standardized prior to benchmarking. Genomic regions were resized to a fixed width of 200 bp, either centered on the original region (PINTS, dREG, cCREs, DHSs, PRIME loci, divergent loci) or on the signal summit (H3K27ac). Reference promoters were defined using the UCSC knownGene annotation as ±500 bp around TSSs. Regions were classified as promoter-proximal or distal relative to these promoters, overlapping regions were merged to avoid redundancy, and regions overlapping coding sequences were excluded. This standardization procedure was applied uniformly across all datasets to ensure comparability.

### CRISPR benchmarks

Predictive performance was evaluated using CRISPRi enhancer perturbation data in K562 cells. The CRISPRi benchmark dataset^43^ was previously compiled from three enhancer perturbation screens^46,111,112^, in which targeted repression of candidate regulatory elements was followed by measurement of gene expression effects.

For benchmarking, genome-wide PRIME predictions and comparator regulatory element annotations were intersected with CRISPRi-tested regions. Precision-recall curves were computed by comparing overlaps between predicted regulatory regions and CRISPRi-validated elements across prediction score thresholds.

To further evaluate performance on perturbation datasets with higher sensitivity to small-effect regulatory elements, PRIME predictions were also benchmarked against the DC-TAP-seq K562 enhancer perturbation dataset^68^.

### eQTL benchmark

Enrichment and recall of eQTLs within predicted distal regulatory elements were evaluated using GTEx^53^ v8 whole-blood eQTLs fine-mapped with SuSiE^113^ and FINEMAP^114^ by the Finucane lab (https://www.finucanelab.org/data). Only noncoding variants were retained, and variants were filtered to a posterior inclusion probability (PIP) greater than 0.5 to focus on high-confidence fine-mapped associations. Analyses were restricted to distal regulatory elements.

Predicted regulatory elements were intersected with fine-mapped eQTL variants. Enrichment was calculated relative to a background set of distal noncoding variants from the 1000 Genomes Project obtained from the Price group^115^, and recall was defined as the fraction of eQTL variants overlapping predicted regions. Enrichment-recall curves were generated across prediction score thresholds.

### GWAS benchmark

GWAS benchmarking was performed using UK Biobank variants fine-mapped with SuSiE^113^ and FINEMAP^114^ by the Finucane Lab (https://www.finucanelab.org/data). Traits were grouped into blood-related (HbA1c, Hb, RBC, MCV, MCH, MCHC, Plt), immune-related (WBC, Neutro, Lym, Mono, Eosino, Baso, CRP, IBD, AID_Combined, Asthma, Hypothyroidism, Fibroblastic_Disorders), and liver/metabolic (ApoB, TC, LDLC, TG, Alb, SHBG, TBil, ALT, AST, GGT, Urea, Glucose, IGF1) categories. Only distal noncoding variants with a PIP greater than 0.2 were retained for analysis.

Because CRE annotations were defined on GRCh38, UK Biobank fine-mapped variant coordinates (provided on GRCh37/hg19) were converted to GRCh38/hg38 using UCSC liftOver (hg19ToHg38 chain). Variants that failed liftOver were excluded (mapping rate=0.9978).

Predicted regulatory regions were tested for overlap with GWAS variants. Enrichment was computed relative to a background set of distal noncoding 1000 Genomes Project variants^115^, consistent with the eQTL benchmark. Enrichment–recall curves were calculated across prediction score thresholds to assess the ability of PRIME and comparator annotations to recover trait-associated noncoding variants.

### ClinVar benchmark

ClinVar benchmarking was performed using ClinVar^52^ noncoding variants (variant_summary.txt.gz; ClinVar release 2 March 2026) to assess recovery of disease-associated regulatory variation. Variants were restricted to single-nucleotide variants (SNVs) mapped to GRCh38 and filtered to retain only noncoding variants based on UCSC knownGene annotations.

ClinVar variants were filtered by clinical significance annotation to retain only records annotated as likely pathogenic, pathogenic, or of uncertain significance. Overlap between predicted regulatory elements and ClinVar variants was evaluated. Enrichment was calculated relative to a background set of distal noncoding 1000 Genomes Project variants^115^, consistent with the eQTL and GWAS benchmarks. Enrichment–recall curves were computed across prediction score thresholds.

### PRIME model interpretation

To interpret the features learned by PRIME during training on GM12878 CAGE data, attribution analysis was performed on PRIME-predicted regions in the K562 cell line. Positional importance was quantified using SHAP^64^ (SHapley Additive exPlanations). Specifically, the SHAP TreeExplainer^116,117^ was utilized, optimized for tree-based models such as LightGBM.

SHAP values were computed at single-base resolution across the 401 bp input window of each PRIME predicted positive. To identify consistent positional signatures driving the model’s performance, these values were aggregated to generate mean positional SHAP importance profiles (mean absolute SHAP values per position)..

### PRIME-enhanced human functional annotation map

The analysis of the FANTOM5 dataset was based on the PRIME genome-wide prediction pipeline with additional steps. First, CAGE CTSSs from all FANTOM5 samples were pooled across libraries to construct a global map of transcription initiation. Singleton CTSSs were removed to reduce noise from low-confidence signals.

The first stage of the analysis consisted of applying the PRIME genome-wide prediction pipeline (starting from the tag cluster identification step, as described above). Predicted regions were retrieved using the default pipeline parameters (score threshold = 0.75, d = 0.10), resulting in 2,849,384 regions derived from the pooled FANTOM5 dataset.

In the second stage, facets were constructed by grouping CAGE libraries based on ontology terms representing specific tissues, organs, and primary cell types as previously defined^17^. To complement these physiological facets, we created cell-line-specific facets based on library metadata. For each facet, individual libraries within the corresponding ontology group were aggregated via replicate pooling.

In the third stage, pooled predictions from the first stage were used as a fixed set of candidate regulatory regions for facet-specific scoring. The predicted regions from the first stage were centered on their summit positions and extended by ±200 bp to define 401 bp windows matching the PRIME model input. For each FANTOM5 facet, strand-specific CTSS signals were assigned to these regions and transformed into normalized CAGE profiles as described above. These profiles were then provided as input to the PRIME model to obtain facet-specific prediction scores for each region, which were used for downstream analyses.

This third-stage procedure was implemented as an R function in the PRIMEmodel package (PRIMEmodel::predictFocal()), which enables efficient scoring of predefined genomic regions without re-identification of tag clusters or genome-wide scanning.

### s-LDSC regression analysis across UKBB traits and distal regulatory elements

To evaluate the functional relevance and trait associations of the PRIME-enhanced regulatory atlas, we performed stratified linkage disequilibrium score regression (s-LDSC)^24^. We utilized summary statistics from 4,178 UK Biobank (UKBB) traits, restricted to GWAS phenotypes with high-quality summary statistics provided by the Neale group^74^ (https://nealelab.github.io/UKBB_ldsc/).

Facet-specific PRIME scores were first quantile-normalized to the average facet distribution to ensure comparability across different cell types and tissues. PRIME CREs were stratified into promoter-proximal (overlapping annotated TSSs, 5’ UTRs, or promoters) and promoter-distal (intronic and intergenic) classes. For each facet, we generated binary annotation files by selecting distal regions meeting a PRIME confidence score of 0.5 and 0.75. To account for potential biases in sequencing depth, we restricted each facet set to covering at most the top 100,000 distal regions based on their normalized PRIME scores. As an additional annotation category, FANTOM5 enhancers in GRCh38 coordinates^23^ (obtained from https://zenodo.org/records/556775) were included and processed analogously for comparison.

Partitioned heritability was calculated for each trait across all facets. We estimated trait heritability enrichment, defined as the proportion of heritability explained by the annotation divided by the proportion of SNPs it contains (*h^2^* / *Prop. SNPs*). To enable comparisons across traits, the enrichment scores were mean-centered and scaled within each trait. Significant cell-type-trait relationships were identified by evaluating the enrichment p-values and standard errors across facets.

Relationships between facets were assessed using binary Jaccard distances calculated from the overlap of active (PRIME score > 0.5) distal regulatory elements. Facets were organized via hierarchical clustering with average linkage.

### Interpretation of trait-associated genetic variants

Trait-associated variants and their statistical fine-mapping data were retrieved from the Open Targets Platform using Google BigQuery. We performed a genomic interval-based join between PRIME-defined distal CREs and the credible_set table. GWAS variants were filtered for those overlapping CRE coordinates. To ensure high-confidence associations, we extracted the Posterior Inclusion Probability (PIP) for each variant-study-locus combination. Only variants with a PIP > 0.3 were retained for downstream analysis, increasing the likelihood that the variants overlapping PRIME CREs were likely causal of the phenotypic trait.

To link fine-mapped variants within PRIME CREs to their most probable target genes, we utilized the Enhancer-to-Gene (E2G) portal (available at https://e2g.stanford.edu/) containing links by scE2G^84^ and ENCODE-rE2G^43^. Prioritized loci were further cross-referenced with linked molecular Quantitative Trait Loci (molQTLs), specifically eQTLs.

## Data availability

All CAGE sequencing data generated in this study have been uploaded to the Gene Expression Omnibus Database under accession number GSE306952.

## Supplementary Data

PRIME predictions in K562, GM12878, HepG2, HCT116, A549 cells and FANTOM5 facets as well as facet CAGE signal tracks are available at Zenodo (doi: 10.5281/zenodo.19712783, v0.2.0).

An interactive genome browser for visualizing all PRIME predictions is available at https://anderssonlab.org/PRIMEatlas/.

**Supplementary Table 1** is available at: https://github.com/anderssonlab/nucCAGE_PRIME_paper/blob/main/0.External_resources/F ANTOM5/FANTOM5_sample_ontology_mapping.tsv sampleName: sample name; sampelID: CNhs ID; ontology: facet name; data: facet group

**Supplementary Table 2** is available at: https://zenodo.org/records/19712783/files/facet_statistics.tsv?download=1 facet: FANTOM5 cell type or tissue category; proximal_0.5: number of CREs located in proximal regions (score >= 0.5); proximal_0.75: number of CREs located in proximal regions (score >= 0.75); distal_0.5: number of CREs located in distal regions (score >= 0.5); distal_0.75: number of CREs located in distal regions (score >= 0.75); all_0.5: number of CREs (score >= 0.5); all_0.75: number of CREs (score >= 0.75); mean / median: global averages and midpoints across all facets

**Publicly available datasets** used in this study are available from the following sources: reference fasta file for mapping from ENCODE GRCh38_no_alt_analysis_set_GCA_000001405.15; CAGE data from FANTOM5 (https://fantom.gsc.riken.jp/5/datafiles/phase2.0/); GM12878 GRO-cap from ENCODE (ENCFF638SZH, ENCFF740NAF); GM12878 NET-CAGE (GSE118075); GM12878 bulk RNA-seq (ENCFF345SHY); ENCODE blacklist regions (ENCFF356LFX); PINTS peaks from GRO-cap and PRO-cap datasets obtainec from the PINTS web portal (https://pints.yulab.org/); ENCODE candidate cis-regulatory elements (cCREs) for K562 from the SCREEN web portal (https://downloads.wenglab.org/Registry-V4/ENCFF414OGC_ENCFF806YEZ_ENCFF849T DM_ENCFF736UDR.bed); dREG peaks derived from K562 PRO-seq and qPRO-seq data (GSE150625); K562 H3K27ac ChIP-seq peaks from ENCODE (ENCFF038DDS); DNase hypersensitive sites (DHSs) from the NIH Roadmap Epigenomics Consortium (GM12878: E116, K562: https://egg2.wustl.edu/roadmap/data/byFileType/peaks/consolidated/narrowPeak/E123-DNase.macs2.narrowPeak.gz); SNPs from the 1000 Genomes Project obtained from the Price group (https://alkesgroup.broadinstitute.org/LDSCORE/baseline_v1.1_hg38_annots/); K562 CRISPR perturbation data obtained from the Engreitz group (https://github.com/EngreitzLab/CRISPR_comparison/blob/main/resources/crispr_data/EPCri sprBenchmark_combined_data.training_K562.GRCh38.tsv.gz); summary statistics from 4,178 UK Biobank (UKBB) traits obtained from the Neale group (https://nealelab.github.io/UKBB_ldsc/); ClinVar noncoding variants (https://ftp.ncbi.nlm.nih.gov/pub/clinvar/tab_delimited/variant_summary.txt.gz; release 2 March 2026); fine-mapped UK Biobank GWAS variants obtained from Finucane group (https://www.finucanelab.org/data); FANTOM5 enhancers in GRCh38 coordinates (https://zenodo.org/records/556775); fine-mapped GTEx eQTLs obtained from Finucane group (https://www.finucanelab.org/data).

## Code availability

CAGE sequencing data were preprocessed using the PRIMEprep pipeline (v0.1.0) (https://github.com/anderssonlab/PRIMEprep/releases/tag/v0.1.0), incorporating FastQC (v0.12.1), fastp (v0.23.4), rRNAdust (v1.02), STAR (v2.7.3a), samtools (v1.21; htslib v1.21), and preseq (v2.0).

Downstream analyses were performed in R (v4.2.2) using PRIME (v0.1.0) (https://github.com/anderssonlab/PRIME/releases/tag/v0.1.0) and CAGEfightR (v1.18.0). Genome annotation and processing utilized TxDb.Hsapiens.UCSC.hg38.knownGene, GenomicRanges (v1.50.2), and related Bioconductor infrastructure. Visualization was performed using Gviz (v1.42.1).

The PRIME model was implemented in Python (v3.9.9) using LightGBM and scikit-learn, with hyperparameter optimization performed using Weights & Biases (wandb). The PRIME model (v1.0.0) and genome-wide prediction pipeline is available at https://github.com/anderssonlab/PRIMEmodel/releases/tag/v1.0.0.

All code required to reproduce the analyses is available at https://github.com/anderssonlab/nucCAGE_PRIME_paper/releases/tag/v1.0.0 (v1.0.0).

