## Supplementary Material for "Mapping active cis-regulatory elements from transcription initiation events"

#### Contents


Supplementary Figures 2
- Supplementary Figure 1 | Optimization of nuclei extraction protocol. 2
- Supplementary Figure 2 | Individual nucCAGE replicates consistently confirm higher sensitivity. 3
- Supplementary Figure 3 | Benchmarking CRE predictions by genetic variants stratified by gene proximity. 4
- Supplementary Figure 4 | Characterization of RNA yield in GM12878 and K562 CAGE libraries. 5
- Supplementary Figure 5 | Characterization of library complexity and noise estimates in GM12878 and K562 CAGE libraries. 6
- Supplementary Figure 6 | Assessment of regulatory element detection capacity and core promoter complexity. 7
- Supplementary Figure 7 | Correlation of gene-associated expression across nucCAGE, whole-cell CAGE, GRO-cap, and RNA-seq. 8
- Supplementary Figure 8 | Characterization of CAGE-derived divergently transcribed loci. 9
- Supplementary Figure 9 | Construction of training dataset and strategy for PRIME model training. 10
- Supplementary Figure 10 | PRIME genome-wide prediction pipeline. 11
- Supplementary Figure 11 | PRIME model performance on held-out test dataset. 12
- Supplementary Figure 12 | Positional importance of transcription initiation.. 13
- Supplementary Figure 13 | Intersections of true and false positive regulatory element predictions across benchmarking methods. 14
- Supplementary Figure 14 | Regulatory architecture of the MYC gene locus. 15
- Supplementary Figure 15 | Hierarchical organization and quantification of the PRIME-enhanced FANTOM5 regulatory element atlas. 16
- Supplementary Figure 16 | Cell-type specificity of PRIME FANTOM5 CREs. 17
- Supplementary Figure 17 | Systematic mapping of trait heritability enrichment across UK Biobank traits. 18

Supplementary Notes 19
- Supplementary Note 1: Assays to detect eRNAs and divergent transcription at CREs. 19
- Supplementary Note 2: Trait-associated CREs in non-canonical cellular contexts 20
  - Supplementary Figure N1 | Regulatory architecture of the ABO gene locus. 20
  - Supplementary Figure N2 | Regulatory architecture of the ABCA7 gene locus. 21
- Supplementary Note 3: PRIME-guided interpretation of GWAS variants. 22
  - Supplementary Figure N3 | Regulatory architecture of the SMAD gene locus. 22
  - Supplementary Figure N4 | Regulatory architecture of the NCOR2 gene locus. 23
  - Supplementary Figure N5 | Regulatory architecture of the SDR39U1-CMA1 gene locus. 24
  - Supplementary Figure N6 | Regulatory architecture of the SSBP3-ACOT11 gene locus. 25
  - Supplementary Figure N7 | Regulatory architecture of the CD82 gene locus. 26
  - Supplementary Figure N8 | Regulatory architecture of the IKZF3 gene locus. 27

References 29

### Supplementary Figures

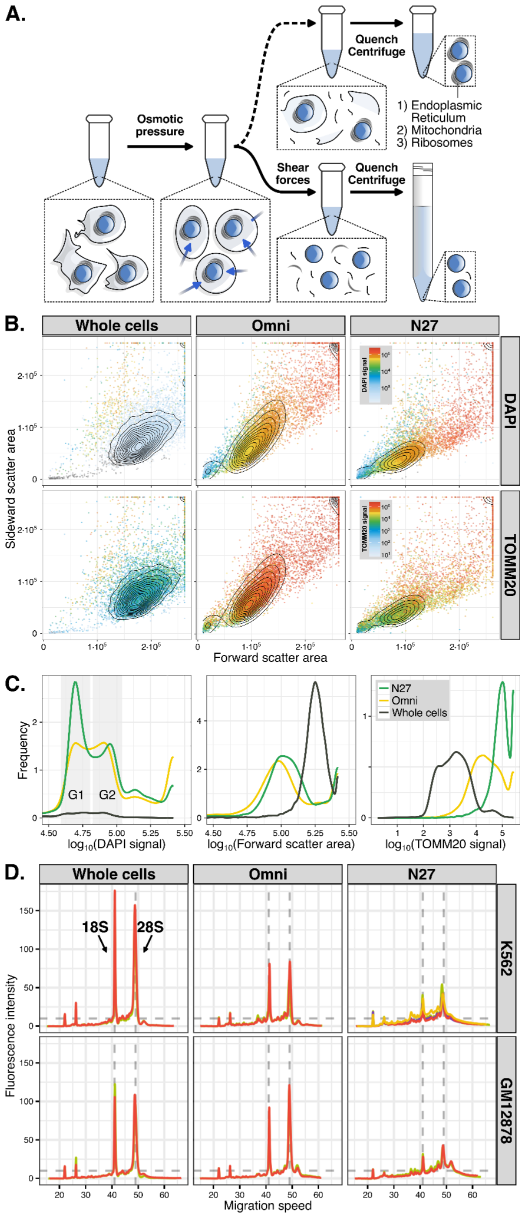


**Supplementary Figure 1 | Optimization of nuclei extraction protocol.**
**A.** Schematic overview of the extraction procedure to acquire nuclei depleted of attached mitochondria and ribosomes. Similar to the nuclei extraction as part of the Omni-ATAC protocol (Omni), osmotic pressure was applied by incubating cells in a hypotonic buffer [ref. 1]. In addition to this purely chemical extraction (upper branch), swollen nuclei were rushed through a 27G-needle to apply shear forces (N27) to remove the endoplasmic reticulum (ER) that is in a membranous continuum with the nuclear envelope. Detaching the ER will also deplete ribosomes and mitochondria from the nuclear extracts (lower branch). **B.** Flow cytometry performed on K562 nuclei extracted with Omni and N27 extraction procedures (see A.) and non-fixed, non-permeabilized whole cells as controls. All nuclei and cells were stained with DAPI and a monoclonal anti-TOMM20 antibody to quantify nuclei sizes and the presence of mitochondria. Unstained and single-stained controls were used to establish detector voltages, but are not shown. **C.** Distribution of forward scatter area, DAPI and TOMM20 signal across whole cells, extracted N27 and Omni nuclei. Data derived from FACS as shown in B. Despite a moderately reduced fraction of N27 nuclei in G1 compared to Omni nuclei (left), they are overall smaller (middle) and possess less attached mitochondria (right). **D.** Distribution of fragment sizes from total, extracted RNA as acquired by automated electrophoresis for GM12878 and K562 whole cells and nuclei. Across all samples and replicates, equal RNA masses were separated for comparability. 2 replicates were processed for all sample types apart from K562 nuclei extracted with N27 procedure, where 4 replicates are shown.

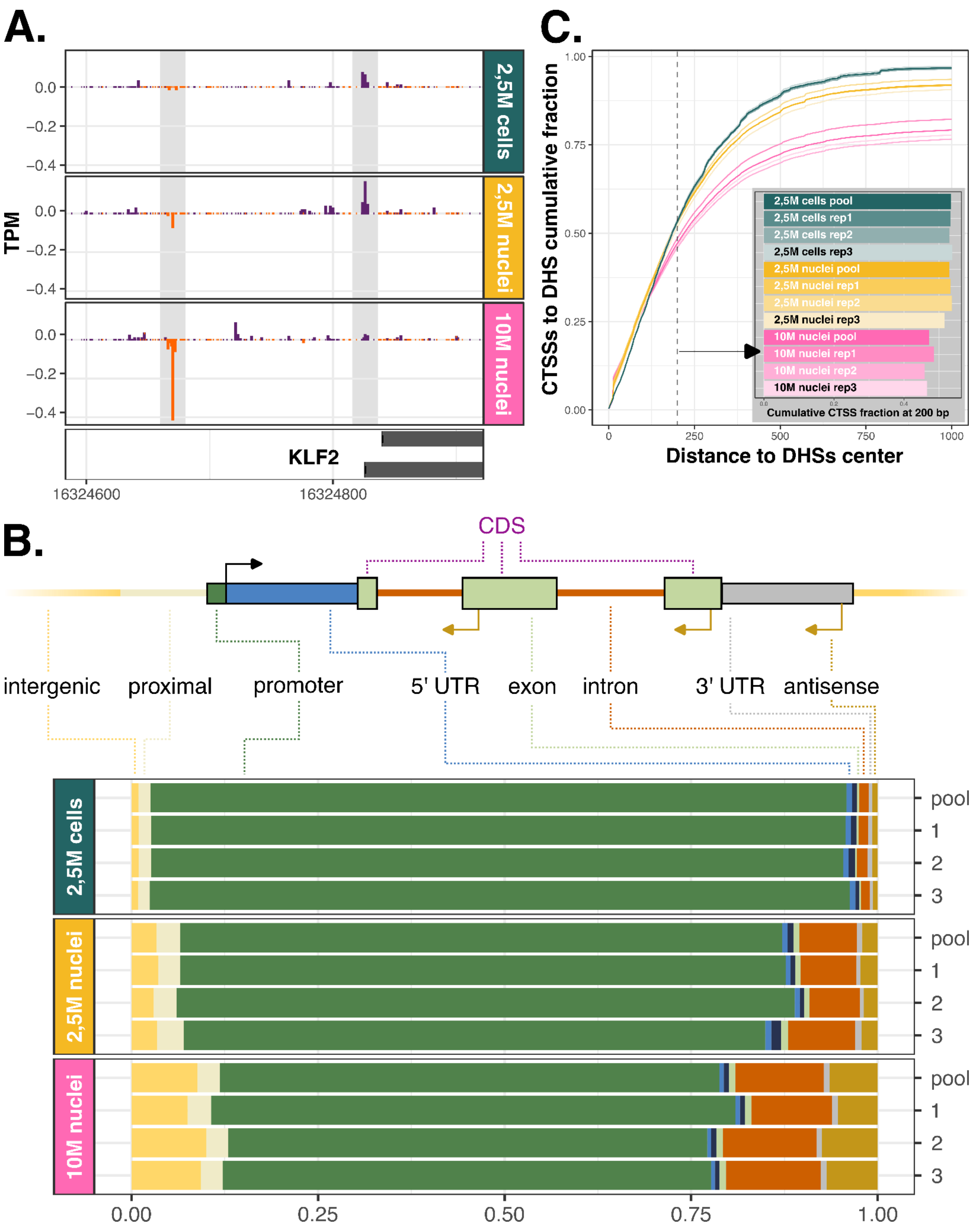


**Supplementary Figure 2 | Individual nucCAGE replicates consistently confirm higher sensitivity.**
**A**. TPM normalized K562 CTSS signal (vertical axes) at the KLF2 mRNA core promoter and its upstream PROMPT (both regions are marked in grey). CTSS signal on the minus strand is displayed as negative TPM values. PROMPTs are detected in nucCAGE samples but are almost absent in whole-cell samples. Conversely, a reduction in mRNA CTSS expression is seen in 10M nuclei samples. **B.** Genomic distribution of identified CTSSs, visualized as stacked bars reflecting the proportion of detected TSSs across annotation categories for individual K562 replicates. The annotations of detected CTSSs replicate the increase of intronic and intergenic fractions in nucCAGE libraries compared to whole-cell samples. **C.** Cumulative fraction (vertical axis) of DHSs covered by CTSSs, as a function of increasing distance between CTSSs and DHS summits.

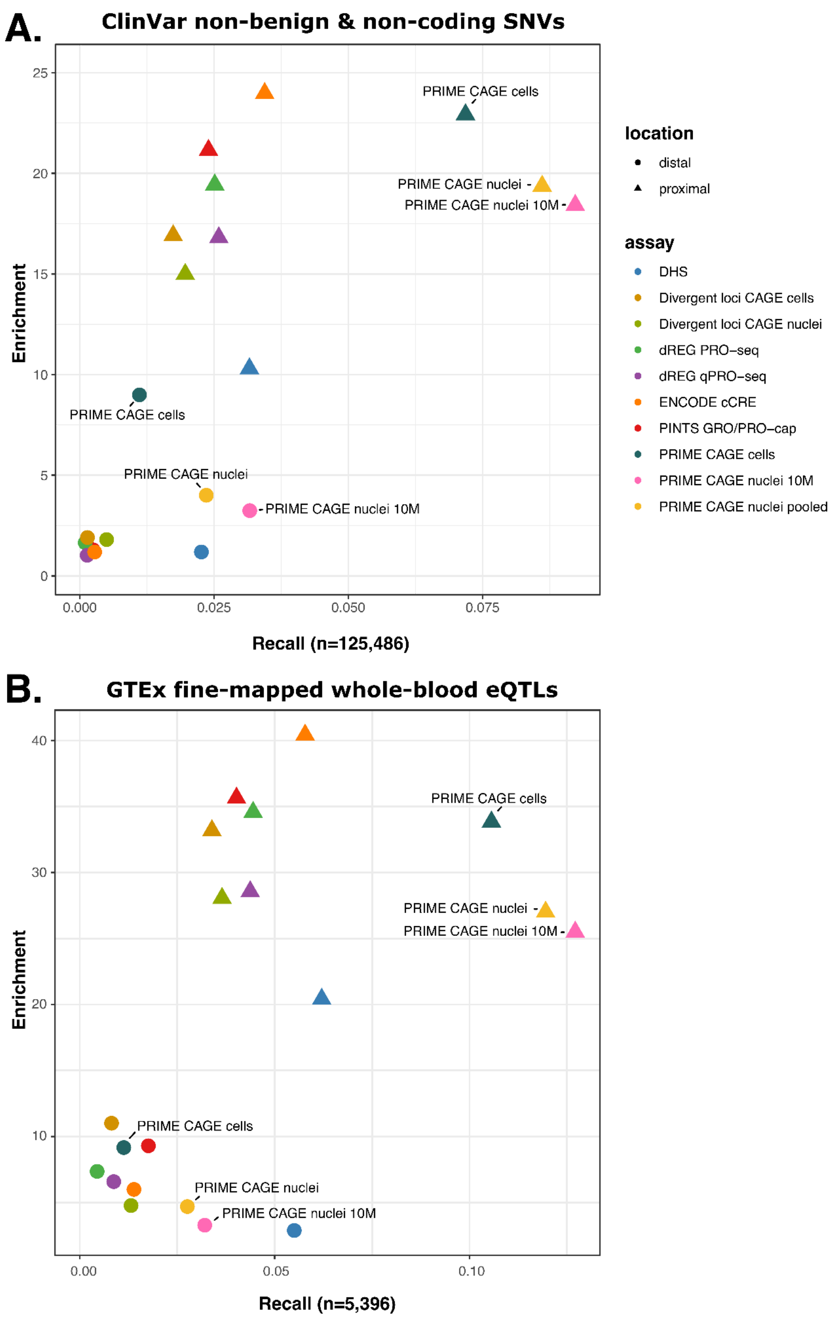


**Supplementary Figure 3 | Benchmarking CRE predictions by genetic variants stratified by gene proximity.** **A**. Enrichment (vertical axis) versus recall (horizontal axis) for predicting pathogenic and likely pathogenic ClinVar single nucleotide variants (SNVs) as well as variants of unknown significance. Enrichment is defined as the ratio of noncoding, non-benign ClinVar SNVs to noncoding common variants overlapping predicted CREs. Recall is the fraction of noncoding, non-benign ClinVar SNVs overlapping predicted CREs. Points indicate performance at recommended thresholds. Results are stratified into gene-proximal (±500 bp) and gene-distal (> 500 bp) classes based on proximity to annotated UCSC knownGene TSSs. **B**. Enrichment (vertical axis) versus recall (horizontal axis) for predicting fine-mapped eQTLs. Enrichment is defined as the ratio of fine-mapped noncoding eQTLs (PIP > 50%) to noncoding common variants overlapping predicted CREs. Recall is the fraction of eQTL variants overlapping CREs. Points and stratification are defined as in **A**.

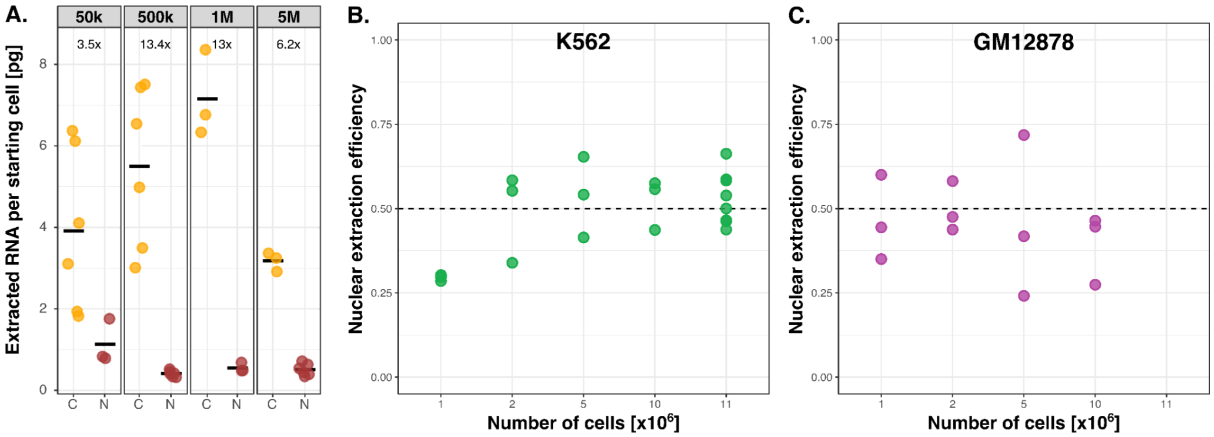


**Supplementary Figure 4 | Characterization of RNA yield in GM12878 and K562 CAGE libraries.**
**A.** Extracted RNA per cell (vertical axes) from whole-cell (C, yellow) and nuclei (N, red) samples, stratified by GM12878 input cell number (horizontal panels). Each dot represents the average yield per starting cell from a separate RNA extraction experiment. **B-C.** Nuclear extraction efficiency (vertical axes), measuring the ratio of isolated nuclei to the total cell population at different number of starting cells (horizontal axis) for K562 (**B.**) and GM12878 (**C.**) cells.

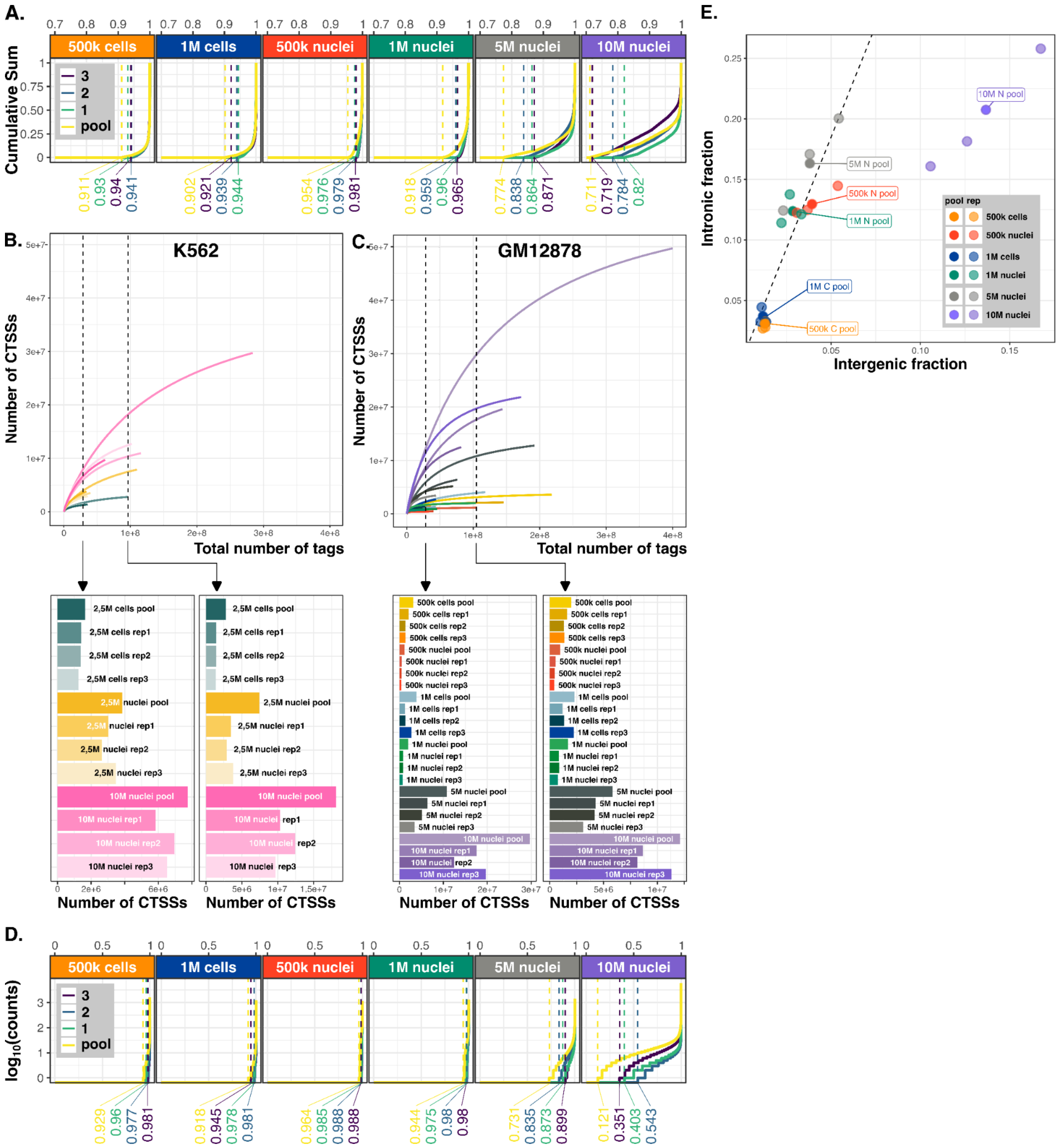


**Supplementary Figure 5 | Characterization of library complexity and noise estimates in GM12878 and K562 CAGE libraries.**

**A.** Fingerprint plots for all replicates and pools across nuclei and cell samples display the amount of genomic bins covered by signal from the corresponding libraries. The dashed vertical lines mark, for each sample, the rank at which the highest-signal CTSSs begin contributing detectable cumulative signal, providing a proxy for signal concentration and library complexity. **B.** Library complexity curves for pooled and individual replicate K562 samples, showing the number of unique CTSSs against the total number of CAGE reads (library size) to estimate the transcriptional diversity and signal-to-noise ratio. **C.** Library complexity curves for pooled and individual replicate GM12878 samples, showing the number of unique CTSSs against the total number of CAGE reads (library size) to estimate the transcriptional diversity and signal-to-noise ratio. Vertical dashed lines in both B. and C. indicate the minimum number of total CTSSs across all replicates (left) or across all pools (right) used to subsample and thus size-match all libraries. **D.** Noise estimates for the same sets of samples as in **A** representing the accumulation of random signal across the genome. The dashed vertical lines mark, for each sample, the rank (quantile) at which background signal first becomes detectable (value > 0) across randomly sampled, mappable, non-masked genomic windows, providing a proxy for noise onset and library background levels. **E.** Fractions of intergenic versus intronic CTSSs for pooled samples and individual replicates. All samples exhibit a linear, scaled increase in these fractions commensurate with input material, with the exception of the 10 million nuclei samples, which deviate from this trend.

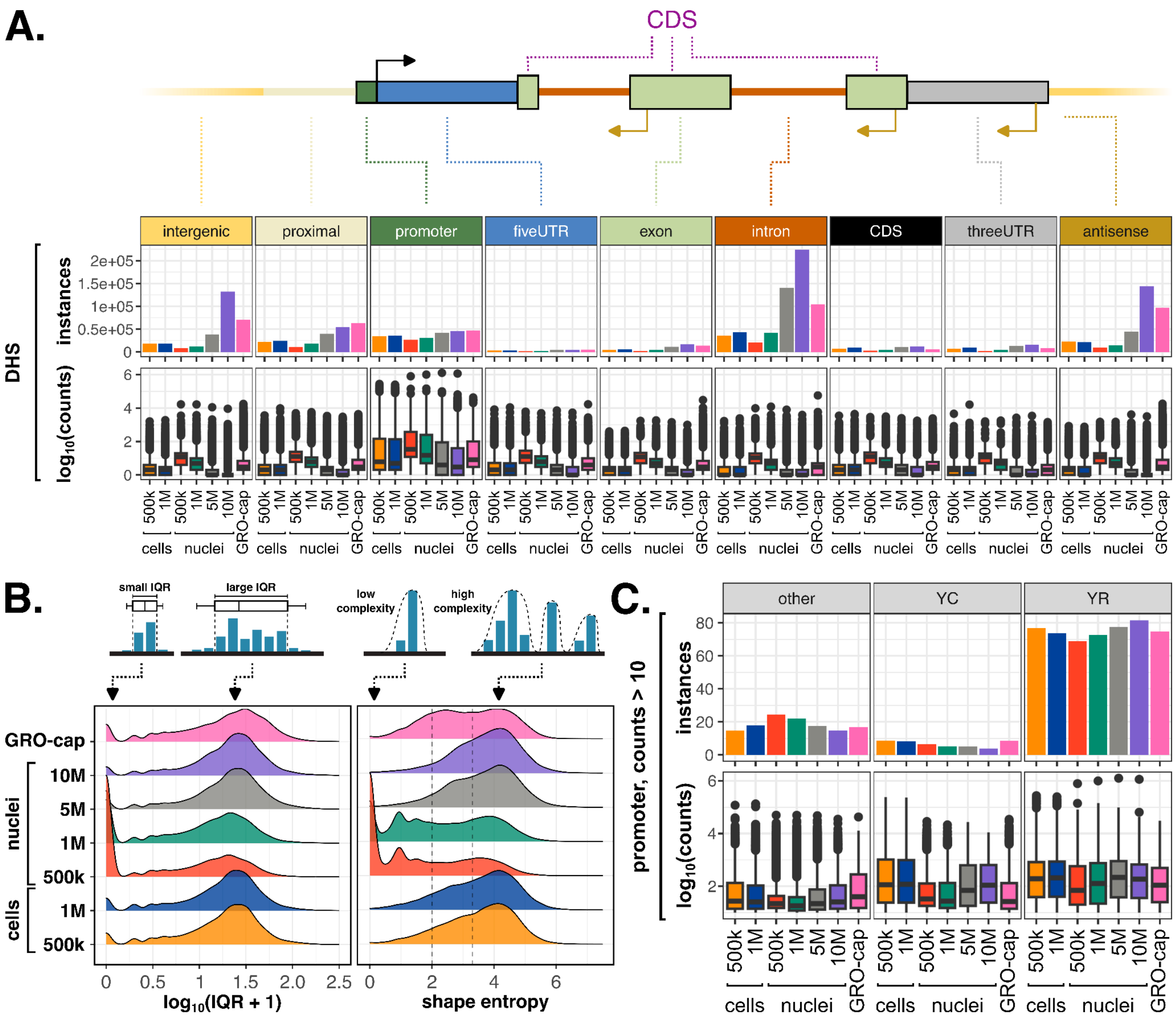


**Supplementary Figure 6 | Assessment of regulatory element detection capacity and core promoter complexity.**

**A.** Number of instances and expression level of core promoters (tag cluster) overlapping DNase I hypersensitive sites (DHSs) across genomic features and GM12878 sample types. Increasing nucCAGE input to 5 million nuclei significantly increases the number of expressed DHSs, exceeding the levels identified by GRO-cap. **B.** CTSS positional complexity of inferred core promoters as measured by 10-90% interquartile range (IQR, left) and shape entropy (right). Low-input nucCAGE samples (500,000 and 1 million nuclei) show reduced positional complexity, while 5 million nucCAGE samples closely resemble the positional complexities of whole-cell CAGE libraries. **C.** Number of instances and expression level of promoter-annotated CAGE-inferred core promoters with >10 counts that carry canonical YC or YR initiator or other motifs across sample types. Five million and 10 million nucCAGE samples display a higher expression of canonical YC initiator-associated TSSs than low-input samples.

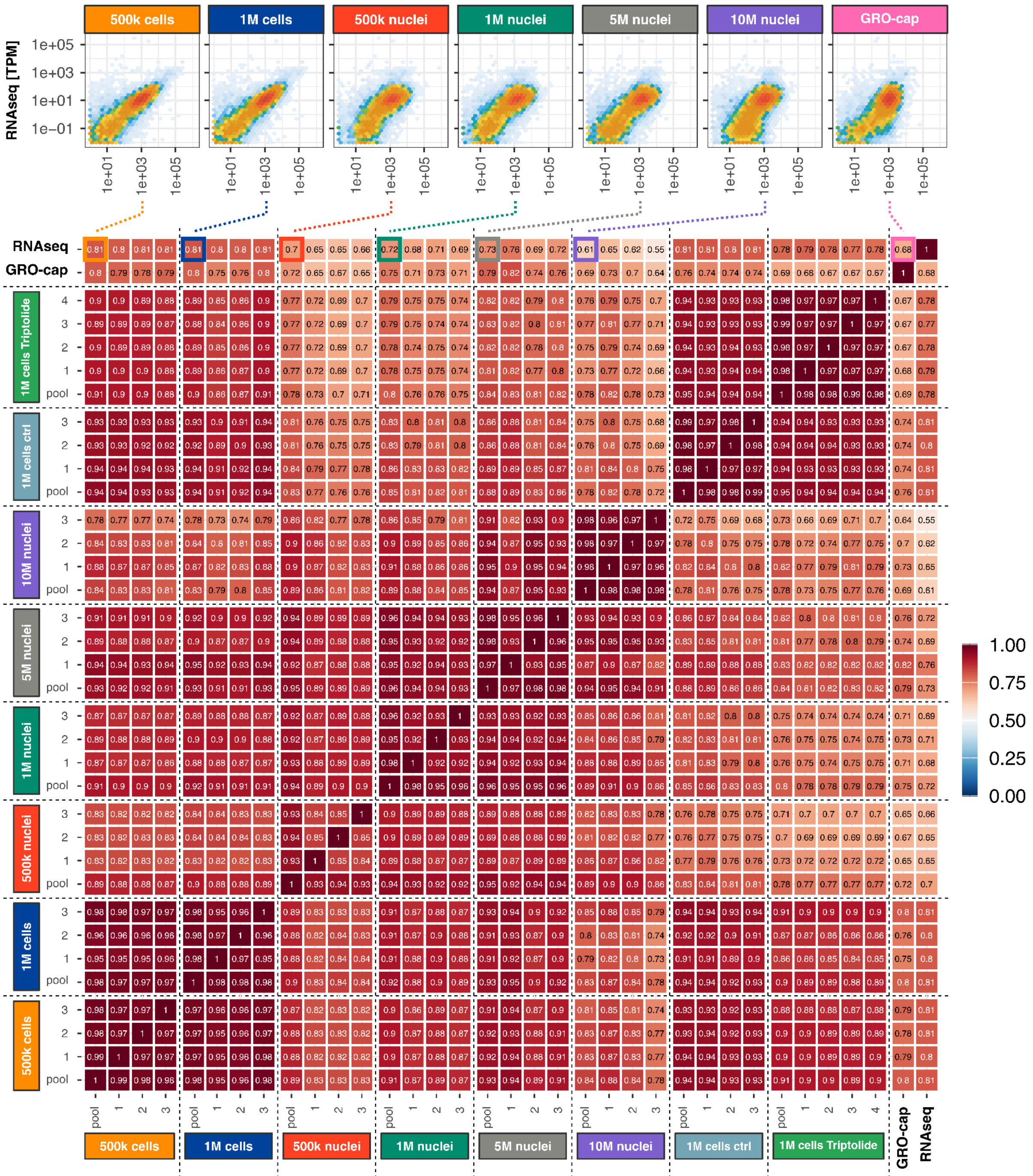


**Supplementary Figure 7 | Correlation of gene-associated expression across nucCAGE, whole-cell CAGE, GRO-cap, and RNA-seq.**
Correlation matrix displaying Pearson correlation coefficients for pairwise comparisons between sample types. Comparisons between RNAseq and nucCAGE, whole cell CAGE, and GRO-cap samples are highlighted through 2D density plots (top).

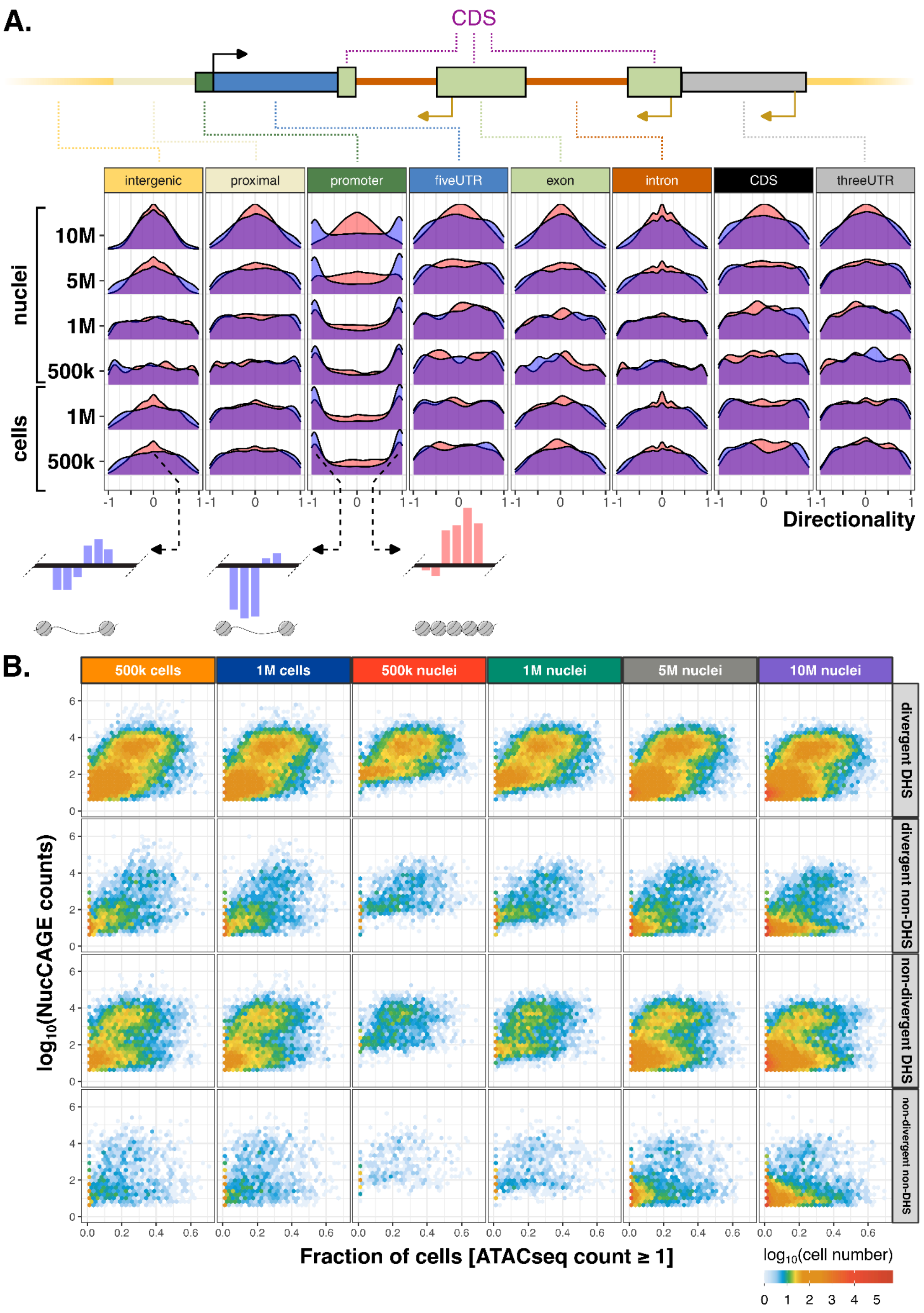


**Supplementary Figure 8 | Characterization of CAGE-derived divergently transcribed loci.**

**A.** Density plots of directionality scores for identified divergent loci separated by genomic features and overlapping (blue) or not overlapping (red) with DHSs. Directionality is calculated as *(P2-M1) / (P2+M1)*, where *P2* is the aggregated plus strand expression of the 200bp genomic window immediately flanking the divergent locus downstream of the inferred mid point and *M1* is the aggregated minus strand expression of the 200bp genomic window immediately flanking the divergent locus upstream of the inferred mid point. **B.** 2D density plot of divergent loci comparing the frequency of snATAC-seq support (fraction of cells with fragment support, horizontal axis) to CAGE expression level (vertical axis), stratified by overlap with DHSs or divergency classification. Color indicates the amount of sites falling into the respective bin. Divergent is classified as *(M1>P1) & (P2>M2)*, where *P2* and *M1* are defined as in **A,** *P1* is defined as the aggregated plus strand expression of the 200bp genomic window immediately flanking the divergent locus upstream of the inferred mid point, and *M2* is defined as the aggregated minus strand expression of the 200bp genomic window immediately flanking the divergent locus downstream of the inferred mid point.

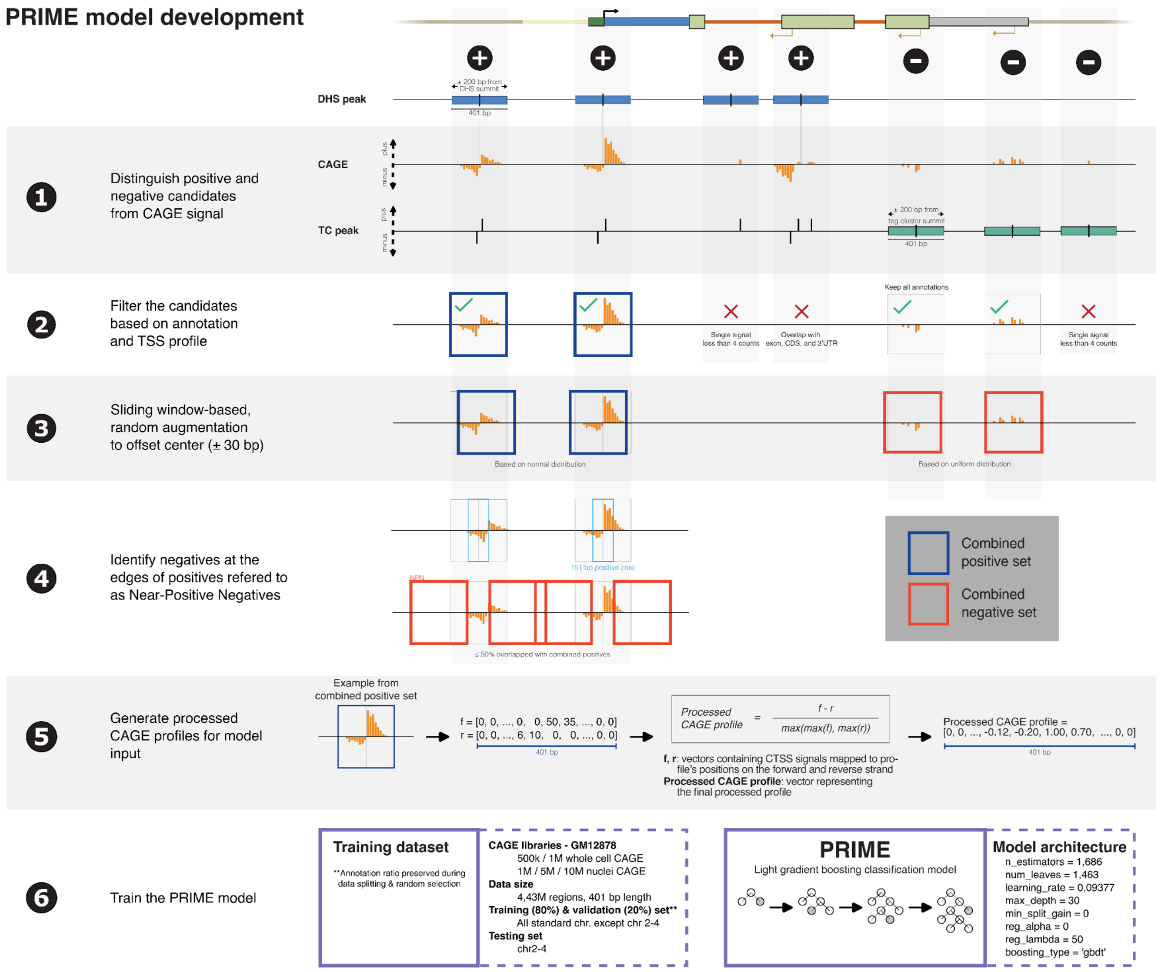


**Supplementary Figure 9 | Construction of training dataset and strategy for PRIME model training.** Schematic illustrating the six major steps for generating the training dataset and training the PRIME LightGBM classifier.

1. **Initial candidate definition**: Positive regions were defined as 401 bp windows centered on DHS summits that contained CAGE signal. Negative regions were defined as 401 bp windows centered on CAGE tag cluster summits that did not overlap any initial positive regions.
2. **Candidate filtering**: All candidates were filtered to exclude regions supported by fewer than four CAGE reads. Positive regions were also filtered to remove overlaps with GENCODE-annotated exons, coding sequences (CDS), and 3′ untranslated regions (3’UTRs).
3. **Random augmentation**: Filtered positive regions were augmented by applying normally distributed random shifts (±30 bp) around the region midpoint to form the combined positive set. Initial negative regions were augmented using uniformly distributed random shifts.
4. **Near-positive negatives**: Near-positive negative regions, defined as sequences flanking the 151 bp core of filtered positive regions with minimal overlap with the positive set, were added to the combined negative set. The combined negative set was randomly subsampled to match the size of the combined positive set.
5. **Profile generation**: Strand-specific CTSS signals (forward, *f*; reverse, *r*) for each region were locally normalized and transformed into a 401-position normalized strand-difference CAGE profile, defined as *(f - r) / max(max(f), max(r))*. This profile preserves local strand asymmetry while removing dependence on global signal amplitude.
6. **Model training**: The processed CAGE profiles were used as input to train the PRIME model, a binary classification LightGBM (LGBM). The final model was trained on 4,428,109 regions, with chromosomes 2-4 held out as an independent test set.

**PRIME genome-wide prediction**

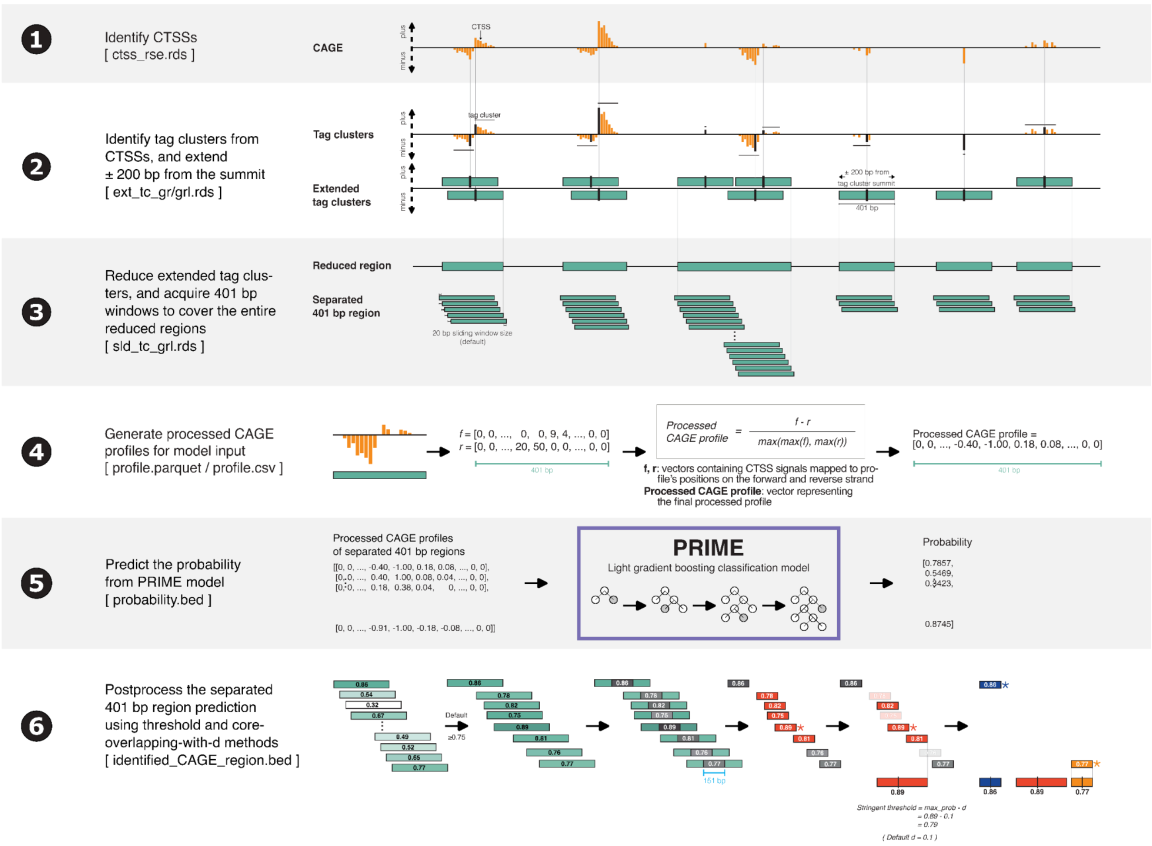


**Supplementary Figure 10 | PRIME genome-wide prediction pipeline.**
Schematic illustrating the six major steps developed to apply the trained PRIME model to TSS data for genome-wide identification of CREs.

1. **Identify CTSSs**: Genome-wide CAGE-inferred Transcription Start Sites (CTSSs) are identified from CAGE bigWig files using CAGEfightR[2].
2. **Identify TCs and extend**: Strand-specific tag clusters (TCs) are identified from CTSSs, and candidate regions are defined by extending the tag cluster summits by ±200 bp to create 401 bp regions.
3. **Reduce and tile regions**: The extended 401 bp regions are reduced to obtain a set of non-overlapping regions, which are then tiled using 401 bp sliding windows with a step size of 20 bp to cover the entire reduced regions.
4. **Generate processed CAGE profiles**: For each sliding window, strand-specific CTSS signals are transformed into a normalized strand-difference CAGE profile. This profile preserves local strand asymmetry while removing dependence on global signal amplitude.
5. **Predict score**: The processed CAGE profiles for each window are provided as input to the PRIME model to obtain a prediction score.
6. **Post-process predictions**: Predicted windows are post-processed using a segmentation strategy, which involves filtering by a global score threshold (default = 0.75), clustering overlapping windows, and iteratively merging retained 151 bp core regions using a local stringency threshold (defined as *max_score* – *d*, where default *d* = 0.1) to define high-confidence, non-overlapping regulatory regions.

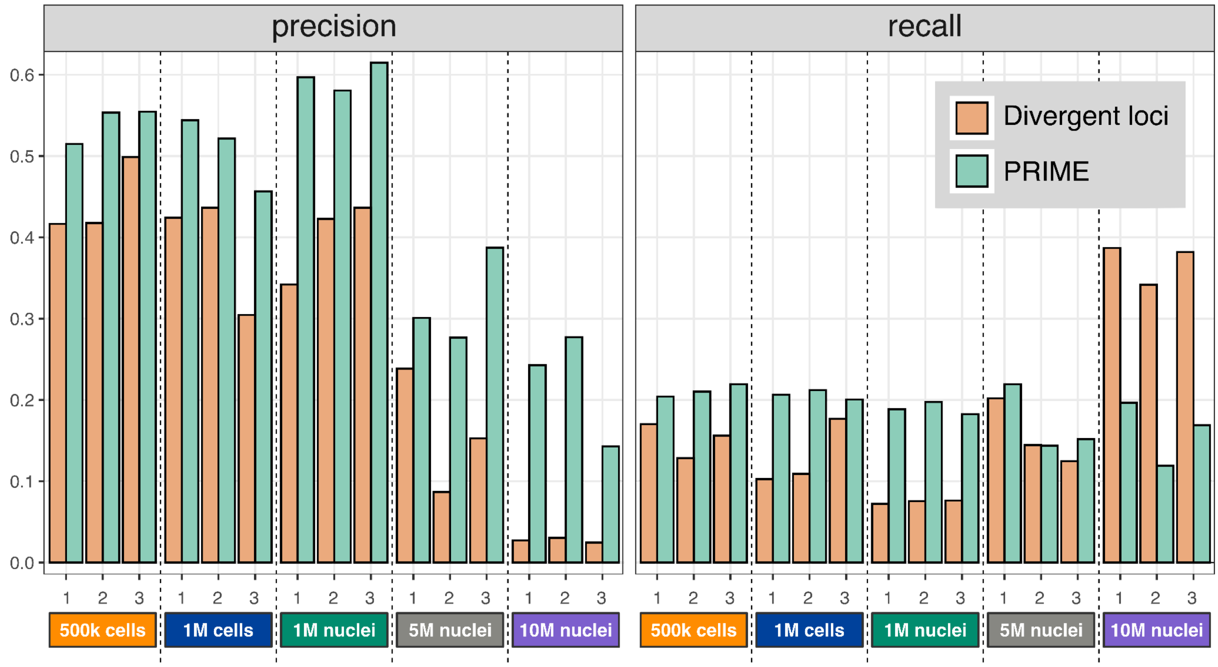


**Supplementary Figure 11 | PRIME model performance on held-out test dataset.**
Precision and recall values for PRIME genome-wide predictions (PRIME score ≥ 0.75, d=0.10) on the independent held-out test set (chromosomes 2-4) compared to divergent loci. PRIME increased both the recall and precision of CREs compared to divergent loci. Precision is defined as the fraction of prediction sites overlapping DNase I hypersensitive sites (DHS) windows of 401 bps with detectable CAGE signal. Recall is defined as the fraction of DHSs with CAGE signal that were recovered by predictions.

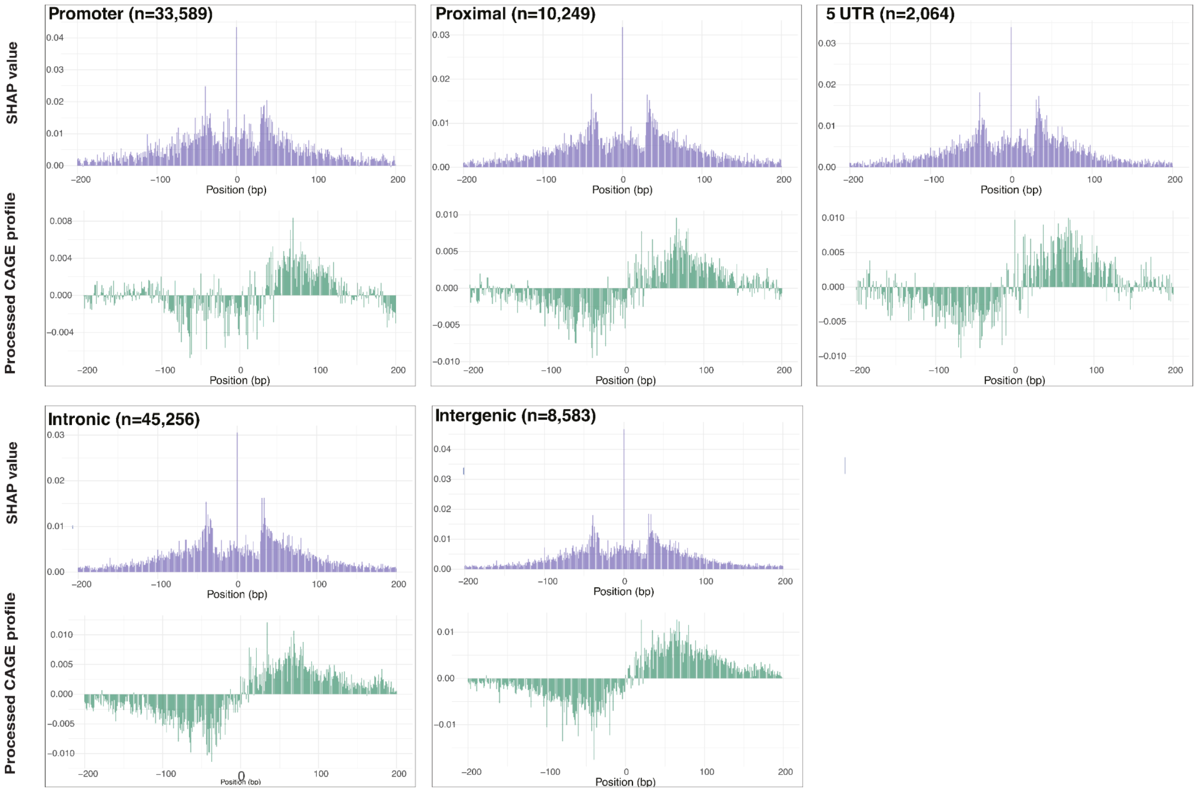


**Supplementary Figure 12 | Positional importance of transcription initiation..**

Histograms showing the average SHAP values (top) and K562 nucCAGE CTSS signal (bottom, positive values: plus strand, negative values: minus strand) at base pairs relative to the K562 PRIME locus center, illustrating the contribution of CTSS signal and position to PRIME classification, stratified by genomic annotation (promoter, proximal region, 5’ UTR, intronic, intergenic).

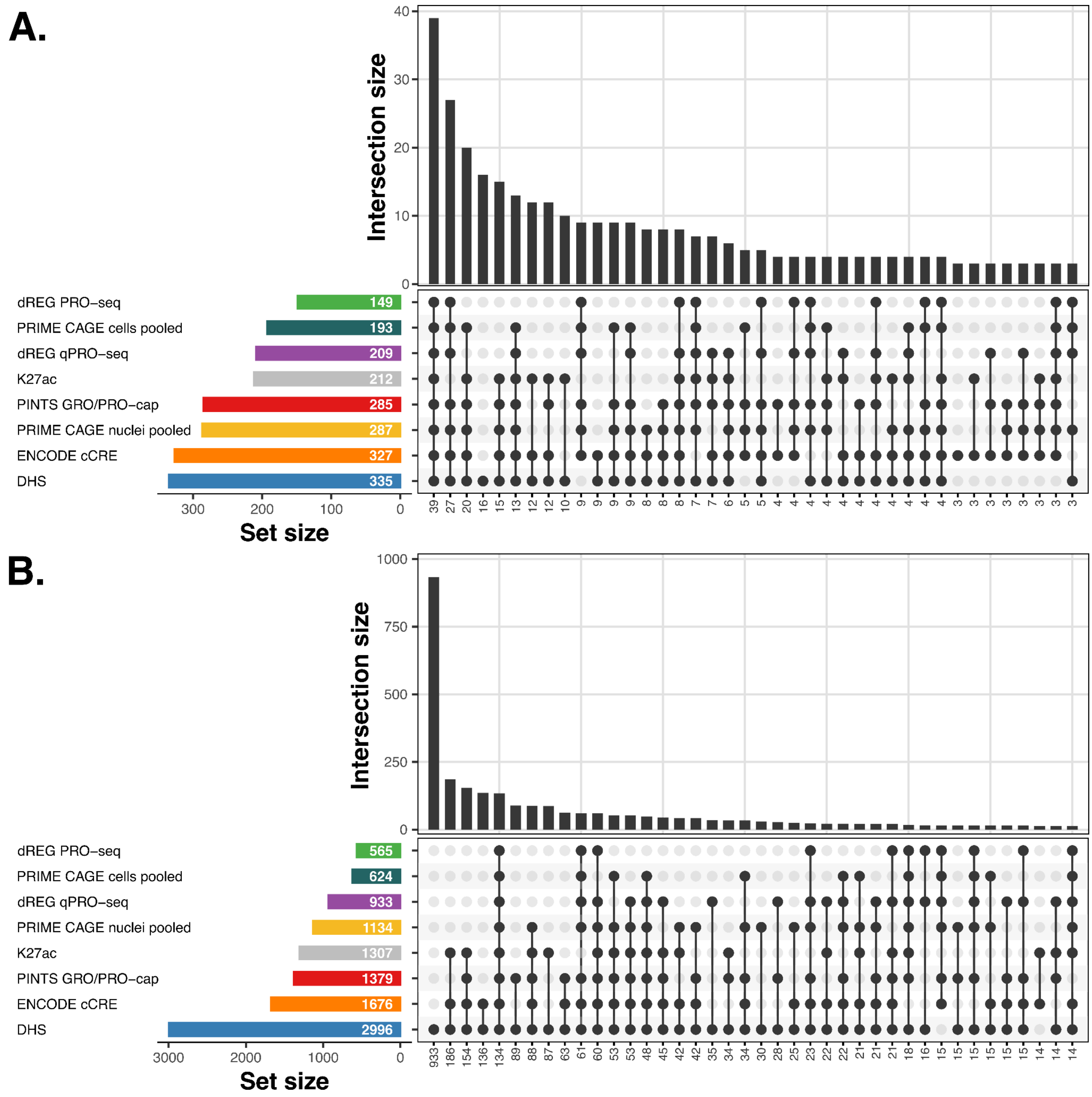


**Supplementary Figure 13 | Intersections of true and false positive regulatory element predictions across benchmarking methods.**

**A:** Upset plot illustrating the overlap of true positive predictions by PRIME and alternative CRE prediction methods (H3K27ac, DHS, ENCODE cCREs, dREG, and PINTS) in K562 cells. True positives are predicted regions that overlap a CRISPRi-validated element. **B.** Upset plot illustrating the overlap of false positives by PRIME and alternative CRE prediction methods in K562 cells. False positives are predicted regions that overlap a CRISPRi-tested negative element.

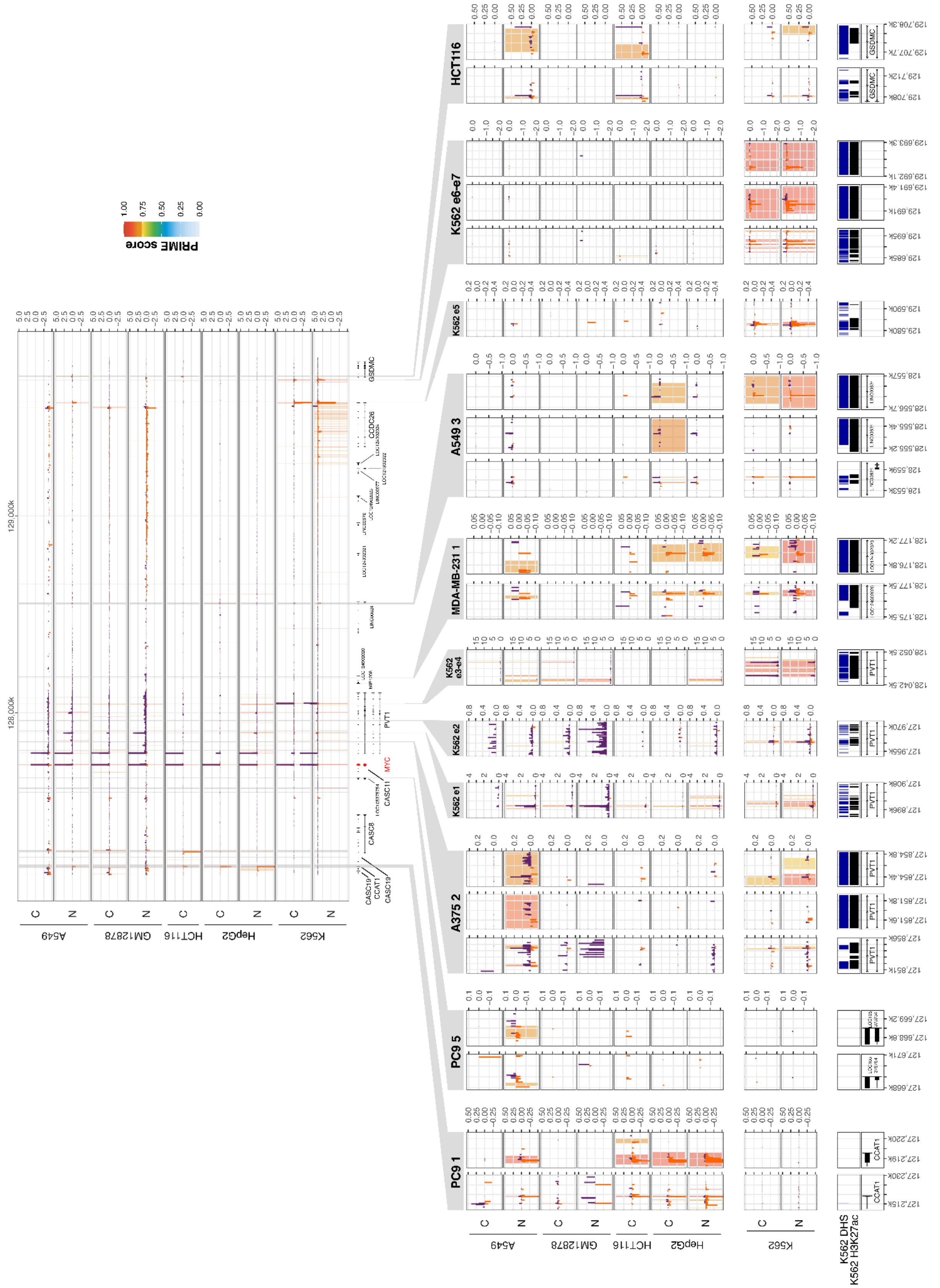


**Supplementary Figure 14 | Regulatory architecture of the *MYC* gene locus.**
CAGE CTSS signal tracks (TPM-normalized) are displayed for K562, HepG2, HCT116, GM12878, and A549 nuclei (N) and cell (C) samples at known CREs to *MYC* and are overlaid with PRIME CRE predictions (vertical bars) colored by PRIME prediction score. For several known CREs, we provide additional zoom-ins, illustrating the positional refinement of CREs achieved by PRIME.

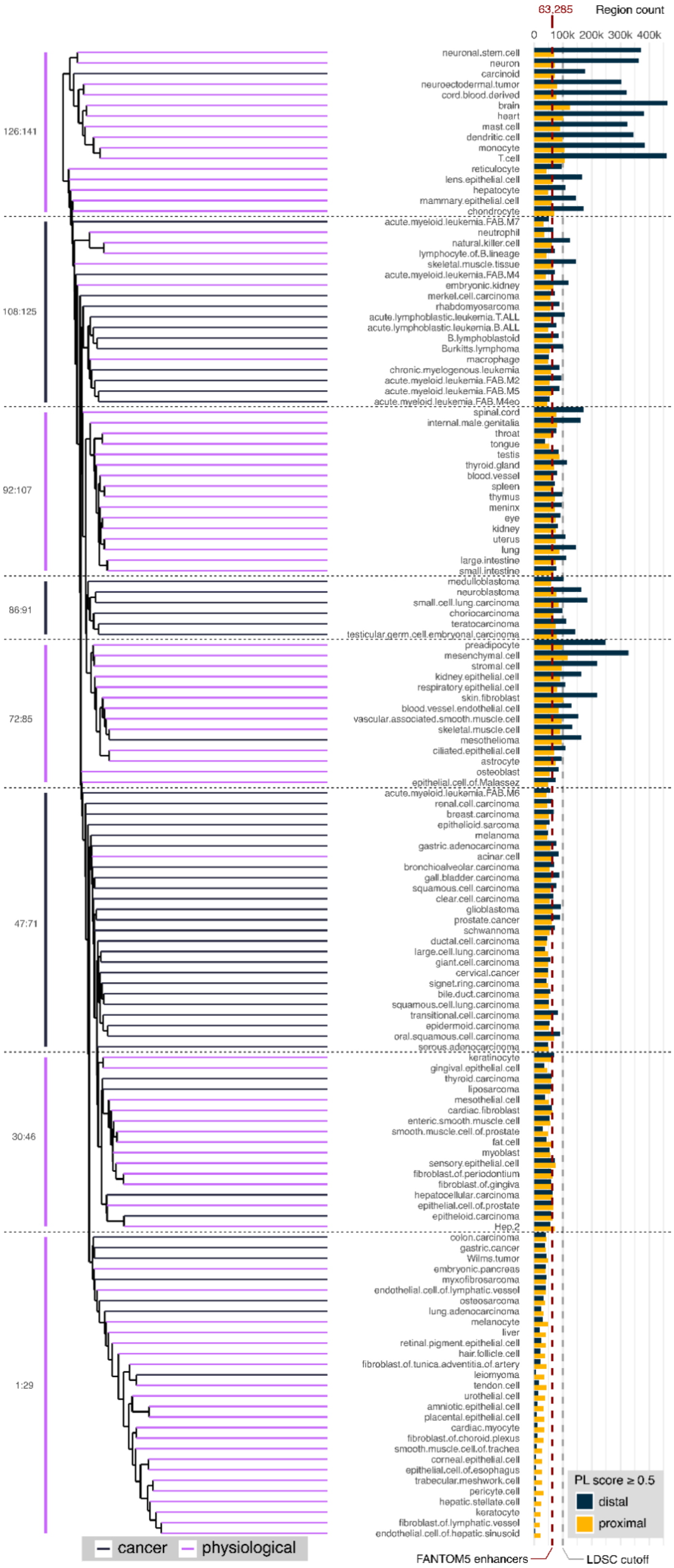


**Supplementary Figure 15 | Hierarchical organization and quantification of the PRIME-enhanced FANTOM5 regulatory element atlas.**

Left: hierarchical clustering based on facet-specific PRIME scores across all 141 cell, tissue, organ, and cell line facets, revealing the organization of the atlas, including a separation between physiological and cancer-derived samples. Right: barplot illustrating the number of distal and proximal CREs identified for each facet, at a minimum PRIME score threshold of 0.5. The number of enhancers identified in FANTOM5 and the maximum number of distal CREs considered for stratified LD score regression (s-LDSC) are depicted by red and grey dashed lines, respectively.

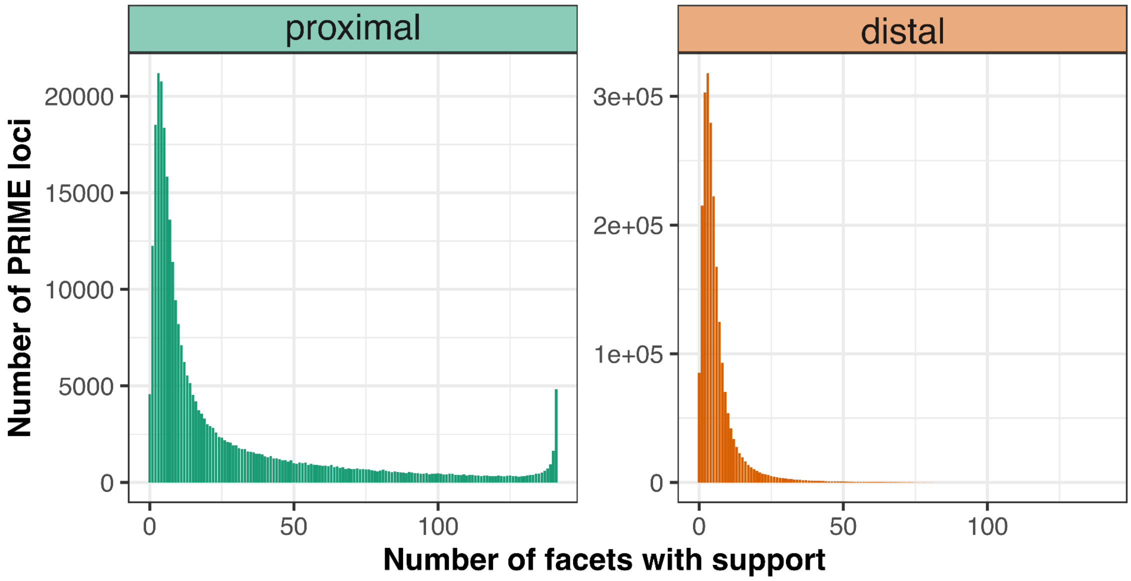


**Supplementary Figure 16 | Cell-type specificity of PRIME FANTOM5 CREs.**
Histogram showing how many facets are supporting each proximal (left) and distal (right) PRIME locus at a score threshold >= 0.5.

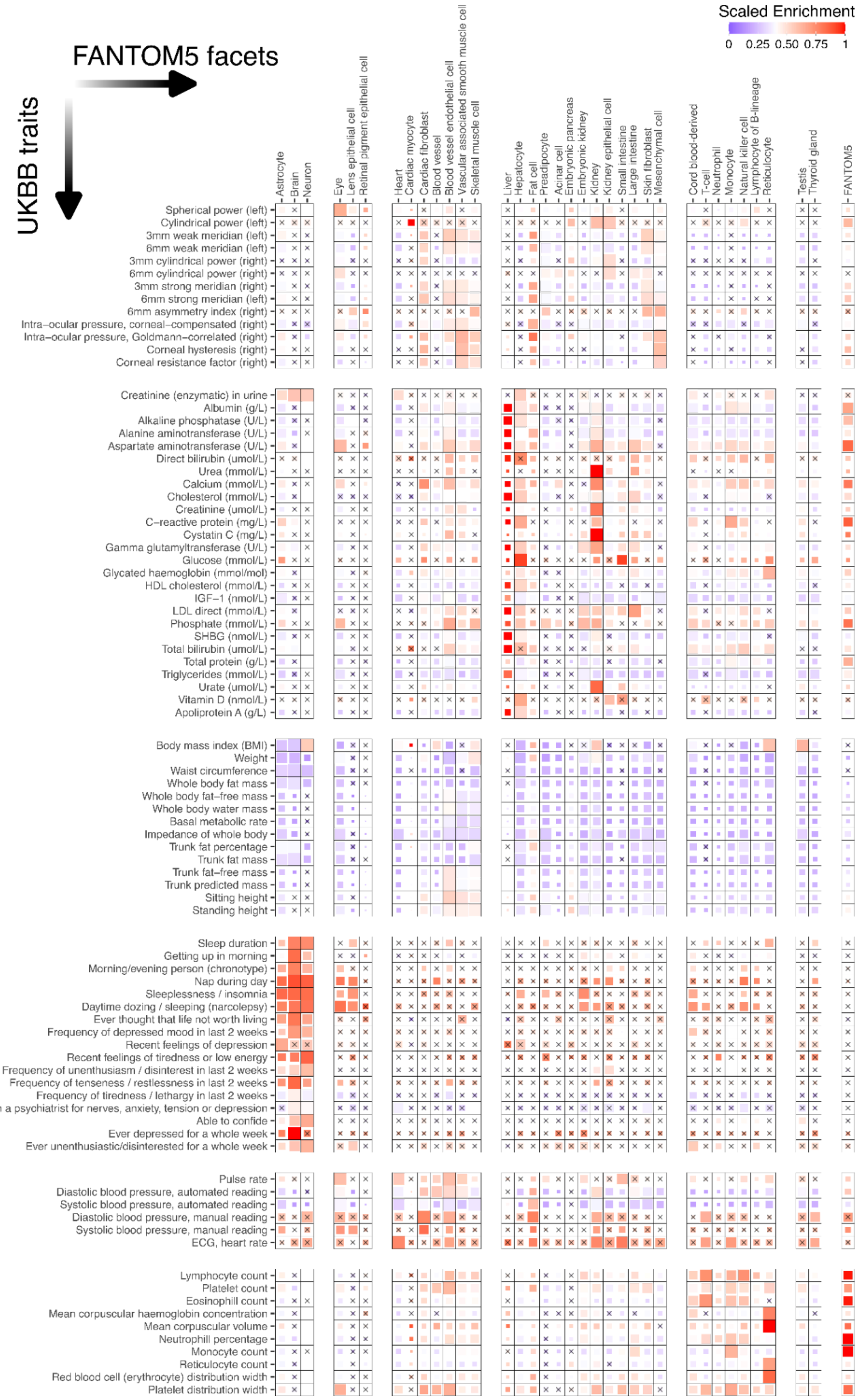


**Supplementary Figure 17 | Systematic mapping of trait heritability enrichment across UK Biobank traits.**
**A.** Scaled heritability enrichment from stratified LD score regression (s-LDSC) for selected UK Biobank traits across a selection of the 141 cell, tissue, organ, and cell line facets in the PRIME-enhanced FANTOM5 atlas. Heritability enrichment is defined as the proportion of heritability explained by the annotation divided by the proportion of SNPs it contains (h^2 / Prop. SNPs). Enrichment scores were mean-centered and scaled within each trait to enable robust comparison. Promoter-distal CREs were selected, and each facet was restricted to a maximum of 100,000 distal regions based on PRIME scores. Non-significant enrichments (nominal p-value > 0.05) are marked with a cross.

#### Supplementary Notes

##### Supplementary Note 1: Assays to detect eRNAs and divergent transcription at CREs.

Mounting evidence supports for regulatory active CREs to be divergently transcribed[3,4]. Similar to mRNA promoters[5,6], enhancers are divergently transcribed during their activation, too, resulting in the synthesis of unstable enhancer RNAs (eRNAs)[3,7–9]. To protect RNA molecules from active degradation by the cellular machinery, 5' 7-methylguanosine caps (5'-m7G-caps) and 3' poly(A)-tails can seal their ends[10,11]. Unlike mRNAs, the majority of eRNAs lack a poly(A) tail and are rapidly degraded[12–18]. Given their short half-life, eRNAs are rare compared to gene-derived mRNAs.

Two types of assays are used to detect eRNAs and thus attempt to overcome their scarcity:

1. Nascent transcript assays, including PRO-cap, profiling polymerase occupancy (i.e., GROseq, PROseq, GRO/PRO-cap, mNETseq, BRUseq, BruUVseq[6,19–23]), and
2. Transcription start site (TSS) assays, including CAGE, enriching for 5' ends of capped RNAs (i.e., CAGE, RAMPAGE, NET-CAGE, CoPRO, Start-seq, csRNAseq, STRIPEseq[24–30]).

Whereas nascent transcript assays measure only transcripts physically located at their DNA locus, TSS assays quantify the pool of nascent, mature or presently degraded RNA species. The rapid degradation of eRNAs reduces the likelihood of their detection with TSS assays[31,32]. As this degradation is however not instant, eRNAs can accumulate moderately, leading to TSS assays scaling differently from nascent transcript assays and thereby influencing the amount of starting material per library required from either assay [33].

It remains open to what degree TSS assays capture nascent transcripts and whether they even miss identifying transiently or dynamically activated CREs. In parallel, while nascent transcript assays measure polymerase occupancy, they can't distinguish between pausing and frequent re-initiation due to transcription termination before entering productive elongation[34–37]. These issues influence the use of the results from either assay type as a proxy for enhancer or promoter activities.

Shared among both types of assays is the basepair-resolution of TSSs at enhancers and promoters. Transcription at enhancers is inherently bidirectional with eRNAs being produced on both strands and thus TSSs mapped divergently[38]. However, gene promoters also give rise to promoter upstream elements (PROMPTs), short-lived transcripts produced on the strand opposite to the mRNA-initiating TSSs[5,6]. Despite the quantitative imbalance between PROMPTs and mRNAs, gene promoters thus share a common architecture of divergent transcription initiation with enhancers[39]. The advantage of TSS assays such as CAGE is their ability to annotate and distinguish enhancers and promoters *de novo* due to the quantitative differences between transcripts on both strands at promoters and the lack thereof at enhancers[3,40,41].

#### Supplementary Note 2: Trait-associated CREs in non-canonical cellular contexts

To illustrate how PRIME can reveal unexpected regulatory contexts, we highlight examples where variants linked to specific traits overlap CREs active in cell types not classically associated with those traits.

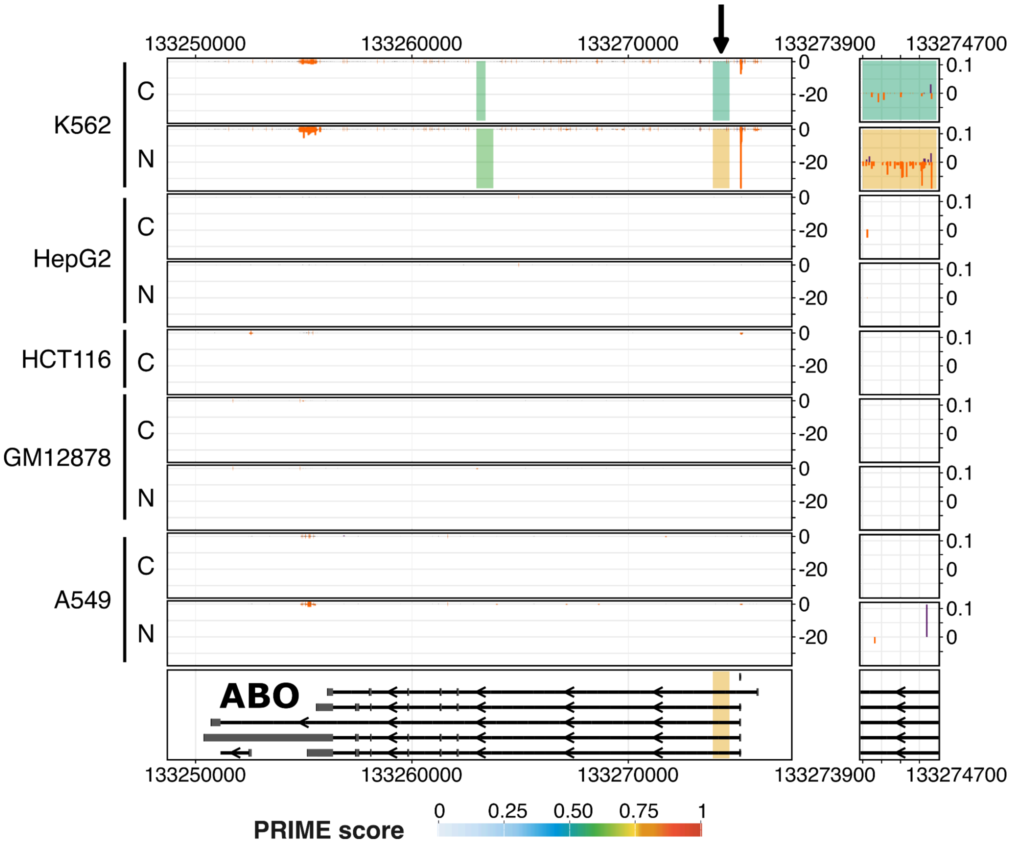


**Supplementary Figure N1 | Regulatory architecture of the *ABO* gene locus.**
CAGE CTSS signal tracks (TPM normalized) for K562, HepG2, HCT116, GM12878, and A549 nuclei (N) and cell (C) samples at the *ABO* gene locus. PRIME CRE predictions (vertical bars) are overlaid and colored by PRIME prediction score. A CRE cluster in the first intron of *ABO* (zoomed in to the right) overlaps a variant (rs115478735, 9_133274295_A_T) associated with ApoB levels.

SNVs associated with Apolipoprotein B (ApoB) were enriched not only in HepG2 (**Fig. 4E**) but also in K562 PRIME loci. Candidate CREs (chr9:133273984:133274255, chr9:133274684-133274835) identified by PRIME in the first intron of the *ABO* gene (**Fig. SN1**), active in K562 and flanking an ApoB-associated SNV (rs115478735, 9_133274295_A_T), support a mechanistic connection between erythroid regulation and lipid metabolism. Variants affecting ABO glycosyltransferase activity could influence the capacity of erythrocytes to bind ApoB-containing lipoproteins, such as LDL[42,43], on their surfaces[44]. This mechanism also offers a potential explanation for the well-established associations between the *ABO* gene and coronary artery disease[45–48], as well as other vascular diseases such as thromboembolism[49,50].

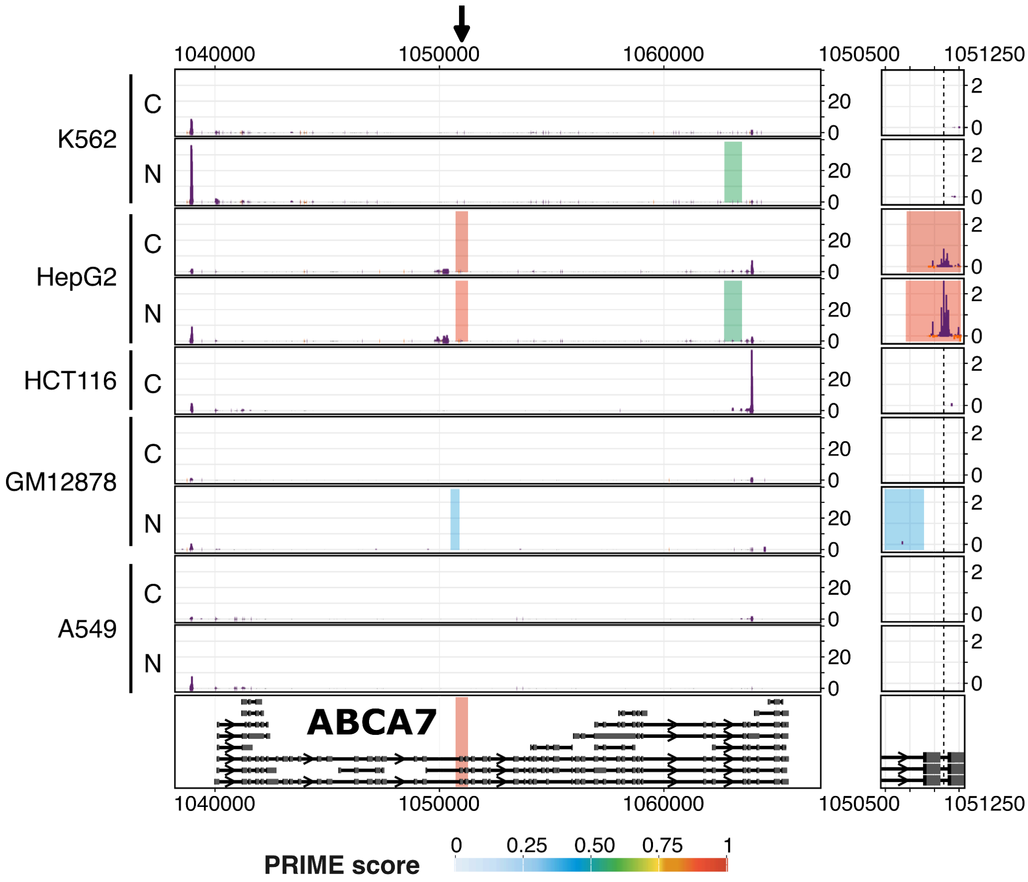


**Supplementary Figure N2 | Regulatory architecture of the *ABCA7* gene locus.**
CAGE CTSS signal tracks (TPM normalized) for K562, HepG2, HCT116, GM12878, and A549 nuclei (N) and cell (C) samples at the *ABCA7* gene locus. PRIME CRE predictions (vertical bars) are overlaid and colored by PRIME prediction score. An intronic PRIME CRE of *ABCA7* (zoomed in to the right) overlaps a variant (rs12151021, 19_1050875_A_G) associated with MCHC levels.

Conversely, a PRIME-inferred CRE within *ABCA7* (**Fig. SN2**), overlapping an SNV (rs12151021, 19_1050875_A_G) associated with mean corpuscular hemoglobin concentration (MCHC), was active in HepG2 cells, even though MCHC reflects the average concentration of hemoglobin in erythrocytes. Because ABCA7 regulates cholesterol efflux[51] from hepatocytes, and erythrocytes participate in reverse cholesterol transport, variants affecting hepatocyte ABCA7 activity may influence hemoglobin-associated cholesterol[52–54] or erythrocyte physiology, for instance through excess cholesterol binding to hemoglobin[55] (i.e., Hb-cholesterol adducts). This interpretation is supported by the importance of cholesterol for *in vitro* erythrocyte culture and by known associations between defects in cholesterol biosynthesis and anemia[56].

#### Supplementary Note 3: PRIME-guided interpretation of GWAS variants.

To illustrate how PRIME facilitates biological interpretation of non-coding GWAS signals, we highlight several trait-associated variants listed by the Open Targets platform[57], where CRE annotations, cell-type activity profiles, and enhancer-to-gene links nominate plausible effector genes and regulatory mechanisms.

##### PRIME locus chr15:67121091-67121491 links lymphoid and myeloid regulation of *SMAD3* to asthma

**Supplementary Figure N3 | Regulatory architecture of the *SMAD* gene locus.**
Regulatory architecture of the *SMAD* gene locus, highlighting a distal CRE (chr15:67121091-67121491, marked by black arrow, zoomed in to the right) with high PRIME activity in immune cell types. Top: ENCODE-rE2G[58] element-to-gene predictions in $CD14^+$ monocytes and activated T cells. Bottom: CAGE CTSS signal tracks (TPM-normalized) for selected FANTOM5 facets overlaid with PRIME CRE predictions (vertical bars) colored by PRIME prediction score.

PRIME identified a CRE (chr15:67121091-67121491) within an intron of the *SMAD3* gene, ~110kb from its transcription start site (**Fig. SN3**). The promoter-distal CRE shows strongest activity in immune cell types, achieving highest PRIME scores in T cells (~0.94), monocytes (~0.90), and mast cells (~0.90). The CRE overlaps the variant rs17294280 (15_67121286_G_A), which is associated with several traits: asthma[59] ($P = 5.77\times10^{-23}$; PIP = 0.995, PICS), FEV/FVC ratio[60] ($P = 1.18\times10^{-12}$; PIP = 0.194, SuSiE-inf), allergic rhinitis[61] ($P = 6.0\times10^{-12}$; PIP = 0.911, PICS). Supporting its regulatory potential, the variant is a known eQTL for *SMAD3* in whole blood (P=0.000011, effect size = -0.23, GTEx[62] phs000424.v10.p2) and is strongly linked to *SMAD3* by scE2G[63] and ENCODE-rE2G[58] in T-cells and monocytes. This regulatory role aligns with the known function of SMAD3 as a transcription factor in the TGF-β signaling pathway[64] and is consistent with prior reports linking *SMAD3* methylation at birth to asthma[65].

##### PRIME locus chr12:124626205-124626605 links monocyte regulation of *NCOR2* to premature separation of placenta (abruptio placentae)

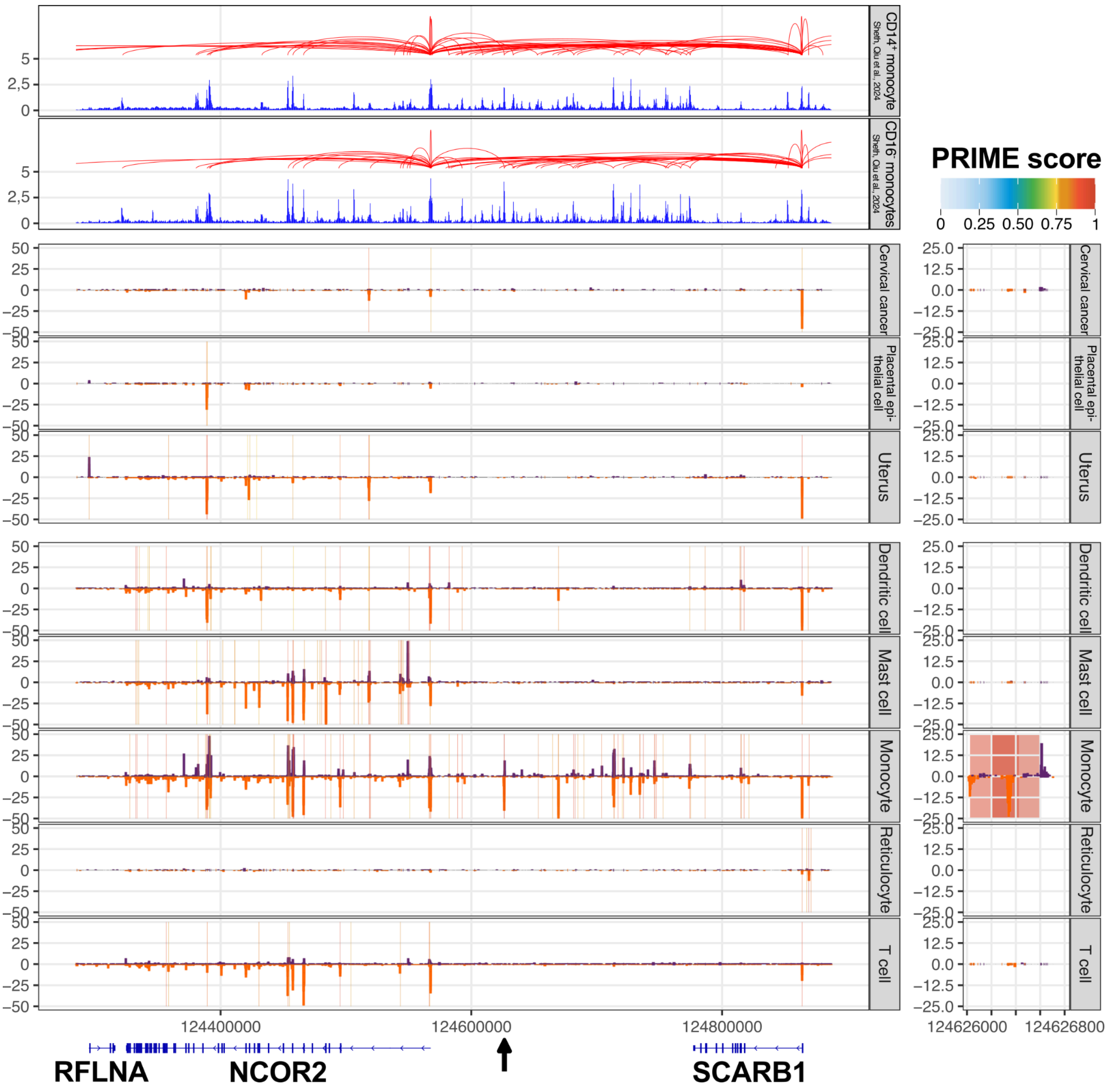


**Supplementary Figure N4 | Regulatory architecture of the *NCOR2* gene locus.**
Regulatory architecture of the *NCOR2* gene locus, highlighting a monocyte-active distal CRE (chr12:124626205-124626605, marked by black arrow, zoomed in to the right). Top: scE2G[63] element-to-gene predictions in $CD14^+$ and $CD16^-$ monocytes. Bottom: CAGE CTSS signal tracks (TPM-normalized) for selected FANTOM5 facets overlaid with PRIME CRE predictions (vertical bars) colored by PRIME prediction score.

PRIME identified a distal CRE (chr12:124626205-124626605), located ~58kb from the *NCOR2* transcription start site (**Fig. SN4**). The CRE shows strongest activity (~0.98) in monocytes and overlaps the variant rs763017549 (12_124626281_C_T), which is associated with premature separation of placenta[66] (abruptio placentae) ($P = 9.80\times10^{-18}$; PIP = 1.0, PICS). Supporting its regulatory potential, the candidate enhancer is strongly linked to *NCOR2* by scE2G[63] in monocytes. NCOR2 is a transcriptional co-repressor that recruits histone deacetylase complexes[67], is known to be involved in regulating trophoblast cell fate[68], and has previously been linked to preeclampsia[69].

**PRIME locus chr14:24517772-24518172 links mast cell regulation of *CMA1* and *SDR39U1* to pernicious anemia**

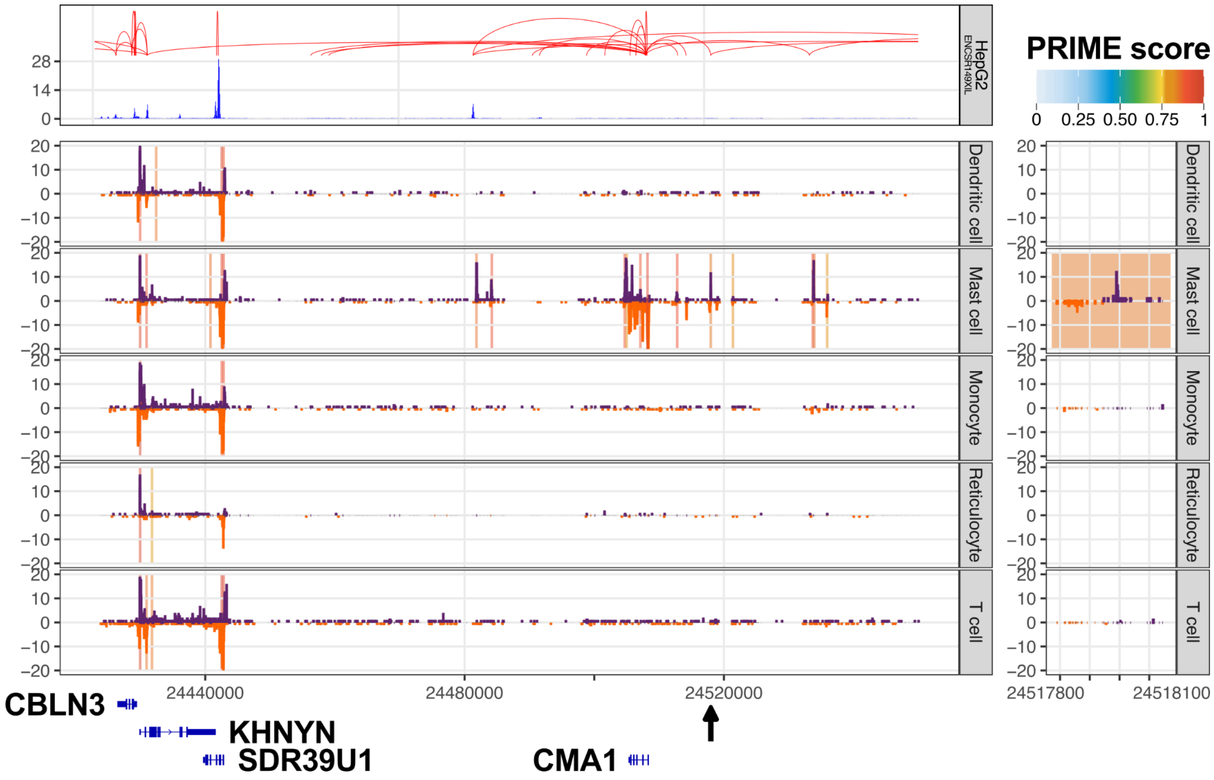


**Supplementary Figure N5 | Regulatory architecture of the *SDR39U1-CMA1* gene locus.**
Regulatory architecture of the *SDR39U1-CMA1* gene locus, highlighting a distal CRE (chr14:24517772-24518172, marked by black arrow, zoomed in to the right) with high PRIME activity in mast cells. Top: ENCODE-rE2G[58] element-to-gene predictions in HepG2 cells. Bottom: CAGE CTSS signal tracks (TPM-normalized) for selected FANTOM5 facets overlaid with PRIME CRE predictions (vertical bars) colored by PRIME prediction score.

We identified a PRIME CRE (chr14:24517772-24518172) within the *CMA1* gene locus, which is most highly scored in mast cells (PRIME score ~0.84) (**Fig. SN5**). This region harbors the variant rs76735545 (14_24518091_C_G), which is associated with pernicious anemia[70] (P = 1.31 × $10^{-11}$; PIP = 1.0, SuSiE-inf), a condition typically caused by vitamin B12 deficiency. The CRE is located ~10kb upstream of the *CMA1* transcription start site and ~75kb from *SDR39U1*. The regulatory potential of this variant is supported by evidence linking it to *CMA1* through ENCODE_rE2G[58] in HepG2 cells, which expresses the gene. Furthermore, variant rs76735545 (14-24518091-C-T) is an eQTL for *SDR39U1* in several tissues, including skeletal muscle (P = 5.7 × $10^{-9}$, effect size = -0.370, GTEx[62] phs000424.v10.p2) and subcutaneous adipose tissue (P = 0.0000031, effect size = -0.34, GTEx[62] phs000424.v10.p2). This suggests that mast-cell regulatory activity at this locus, potentially mediated by mast cell-derived signals such as chymase (encoded by *CMA1*), may be involved in erythroid physiology and the pathogenesis of anemia.

**PRIME locus chr1:54488214-54488614 links lymphoid and myeloid regulation of *ACOT11 / SSBP3* to height**

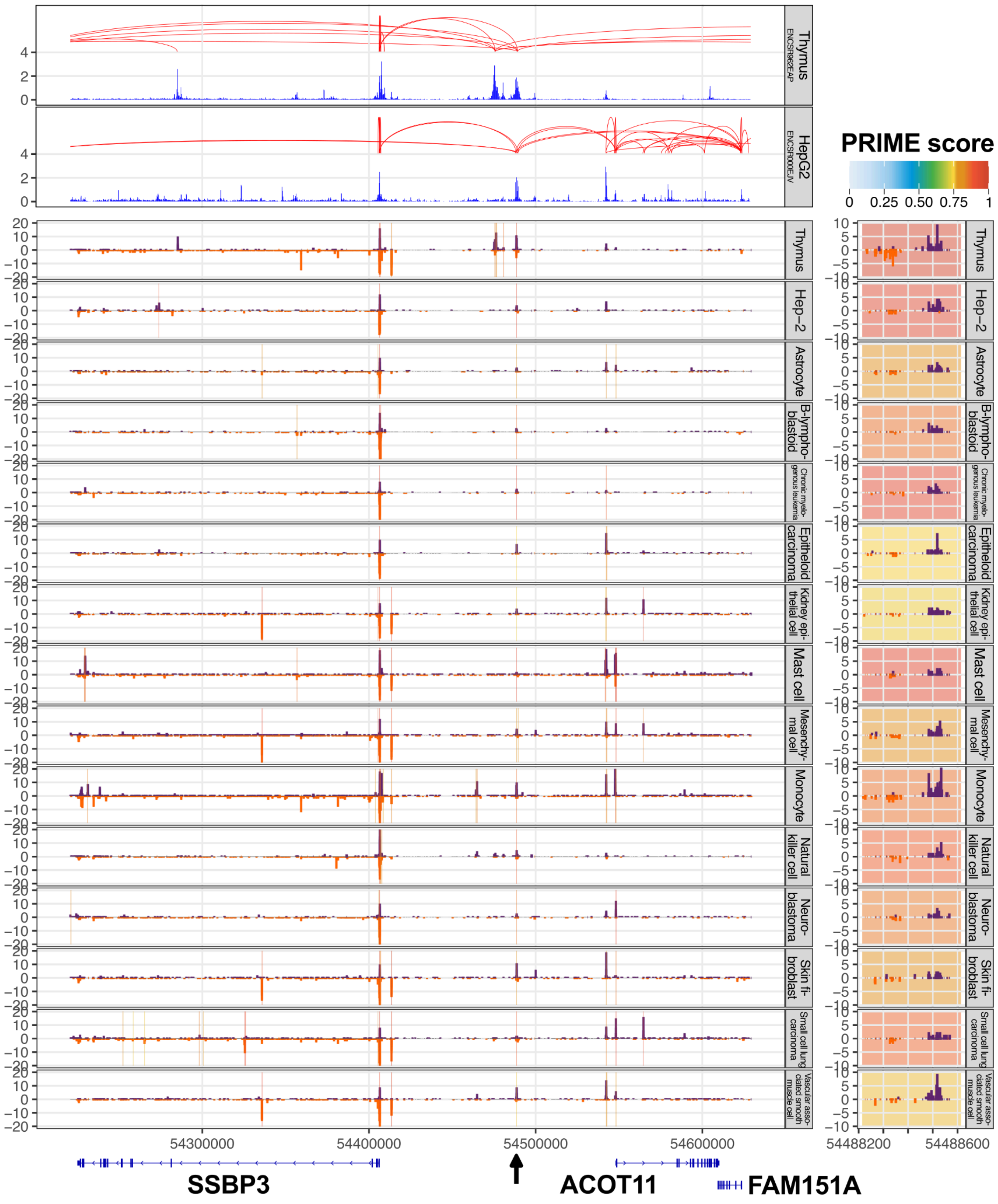


**Supplementary Figure N6 | Regulatory architecture of the *SSBP3-ACOT11* gene locus.**
Regulatory architecture of the *SSBP3-ACOT11* gene locus, highlighting a distal CRE (chr1:54488214-54488614, marked by black arrow, zoomed in to the right) with PRIME activity in lymphoid and myeloid cells. Top: ENCODE-rE2G[58] element-to-gene predictions in Thymus and HepG2 cells. Bottom: CAGE CTSS signal tracks (TPM-normalized) for selected FANTOM5 facets overlaid with PRIME CRE predictions (vertical bars) colored by PRIME prediction score.

PRIME identified a distal CRE (chr1:54488214-54488614), located ~60kb from the ACOT11 transcription start site and ~82kb from the SSBP3 transcription start site (**Fig. SN6**). The CRE exhibits broad activity, scoring most strongly in the thymus (~0.98), as well as in myeloid and lymphoid cell types. The CRE overlaps the variant rs6691924

(1_54488572_C_T), which is strongly associated with height (P = $3.00\times10^{-210}$; PIP = 1, PICS) and also linked to heart rate (P = $3.11\times10^{-19}$; PIP = 0.387, SuSiE-inf). The variant is strongly linked to both ACOT11 and SSBP3 by scE2G[63] in several cell types. This dual regulatory link is notable as both genes have previously been implicated in human anthropometric traits, specifically height[71,72]. Functionally, ACOT11 regulates lipid metabolism and energy expenditure[71]. SSBP3, which is expressed in the thymus (unlike ACOT11), acts as a co-regulator with Ldb1 and is crucial for glucose homeostasis, pancreatic islet architecture, and the maturation of β-cells[73].

**PRIME locus chr11:44570516-44570916 links myeloid regulation of CD82 to platelet count and triglyceride measurement**

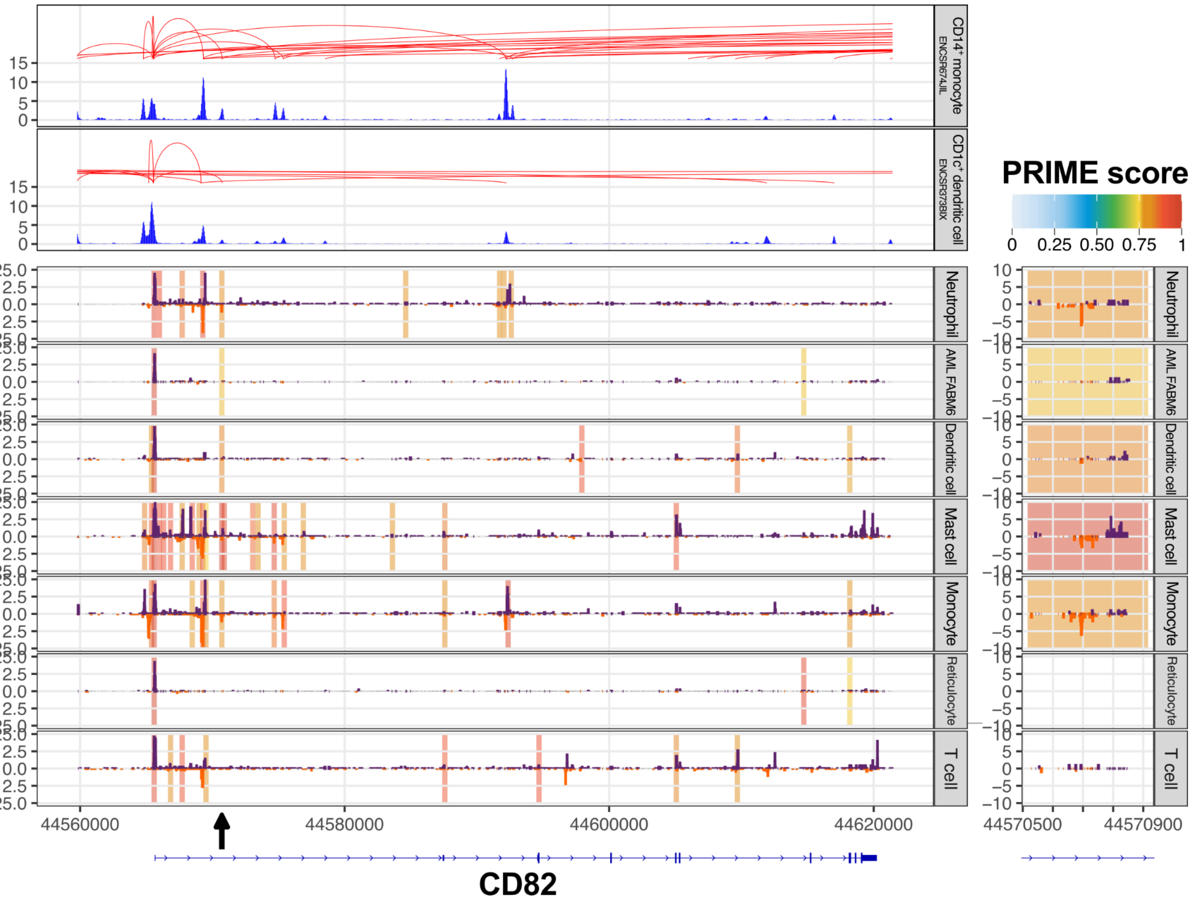


**Supplementary Figure N7 | Regulatory architecture of the *CD82* gene locus.**
Regulatory architecture of the *CD82* gene locus, highlighting a distal CRE (chr11:44570516-44570916, marked by black arrow, zoomed in to the right) with high PRIME activity in myeloid cells. Top: ENCODE-rE2G[58] element-to-gene predictions in CD14$^+$ monocytes and CD1c$^+$ dendritic cells. Bottom: CAGE CTSS signal tracks (TPM-normalized) for selected FANTOM5 facets overlaid with PRIME CRE predictions (vertical bars) colored by PRIME prediction score.

PRIME identified a CRE (chr11:44570516-44570916) in an intron of the CD82 gene, located ~5kb downstream of its transcription start site (**Fig. SN7**). The CRE exhibits strong activity in myeloid cell types, scoring highest in mast cells (~0.98), dendritic cells (~0.84), and neutrophils (~0.83). The CRE overlaps the variant rs56073922 (11_44570719-TTTA-T), which is associated with platelet count (P = $2.28\times10^{-9}$; PIP = 0.985, PICS) and triglyceride measurement (P = $3.46\times10^{-9}$; PIP = 0.310; lead variant 11_44562057_T_A, 95% credible set). Supporting its regulatory potential, the variant is a known eQTL for CD82 in adipose tissue (P = 2.87×10$^{-41}$, effect size = 1.06, twinsUK[74]; PIP = 0.95, SuSiE) and is strongly linked to CD82 by scE2G[63] in myeloid cells. This regulatory role aligns with the known function of CD82 as a tetraspanin protein that acts as a key regulator of myeloid cells in adipose and other peripheral tissues[75].

**PRIME locus chr17:39813772-39814172 links lymphoid and myeloid regulation of IKZF3 to eosinophil count**

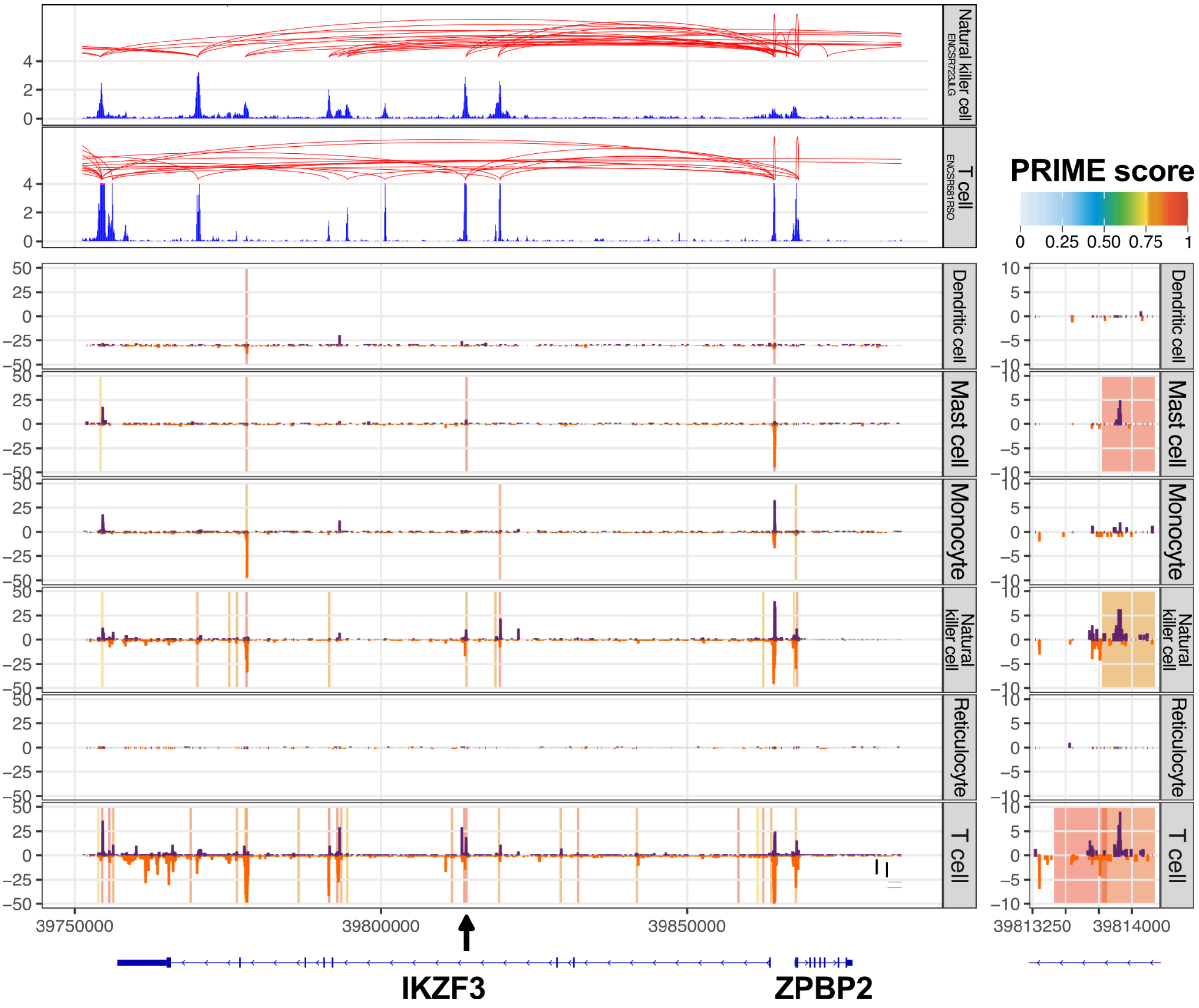


**Supplementary Figure N8 | Regulatory architecture of the *IKZF3* gene locus.**
Regulatory architecture of the *IKZF3* gene locus, highlighting a promoter-distal CRE (chr17:39813772-39814172, marked by black arrow, zoomed in to the right) with high PRIME activity in lymphoid and myeloid cells. Top: ENCODE-rE2G[58] element-to-gene predictions in natural killer cells and T cells. Bottom: CAGE CTSS signal tracks (TPM-normalized) for selected FANTOM5 facets overlaid with PRIME CRE predictions (vertical bars) colored by PRIME prediction score.

The PRIME locus chr17:39813772-39814172 is a CRE situated within an intron of the *IKZF3* gene, located approximately 50kb from its transcription start site (**Fig. SN8**). The candidate enhancer exhibits strong activity in several immune cell types, achieving high PRIME scores in mast cells (~0.89), T cells (~0.87), and natural killer cells (~0.81). It overlaps the variant rs12943633 (17_39814112_C_T), which is strongly associated with multiple traits, including eosinophil count[59] (P = 1.74×10$^{-41}$; PIP = 1.0, PICS), erythrocyte count[59] (P = 1.10×10$^{-15}$; PIP = 0.98, SuSiE-inf), and low density lipoprotein cholesterol measurement[76] (P = 3.50×10$^{-12}$; PIP = 1.0, PICS). Furthermore, the variant is strongly linked to *IKZF3* by scE2G in natural killer cells and T cells. This regulatory element's cell-type specificity aligns with the known function of IKZF3 as a transcription factor that regulates eosinophil development, maturation, and migration[77].
